# Short-chain fatty acid metabolites propionate and butyrate are unique epigenetic regulatory elements linking diet, metabolism and gene expression

**DOI:** 10.1101/2024.01.11.575111

**Authors:** Michael Nshanian, Joshua J. Gruber, Benjamin S. Geller, Faye Chleilat, Samuel Lancaster, Shannon M. White, Ludmila Alexandrova, Jeannie Marie Camarillo, Neil L. Kelleher, Yingming Zhao, Michael P. Snyder

**Affiliations:** Department of Genetics, Stanford University, School of Medicine, Stanford, CA; Vincent Coates Foundation Mass Spectrometry Laboratory, Stanford University, Stanford, CA; Department of Chemistry, Molecular Biosciences and Proteomics Center of Excellence, Northwestern University, Evanston, IL; Department of Biochemistry and Molecular Genetics, Feinberg School of Medicine, Northwestern University, Evanston, IL; Ben May Department of Cancer Research Committee on Cancer Biology, University of Chicago; Chicago, IL; Center for Genomics and Personalized Medicine, Stanford University School of Medicine, Stanford, CA

## Abstract

The short-chain fatty acids (SCFA) propionate and butyrate have beneficial health effects, are produced in large amounts by microbial metabolism and have been identified as unique acyl lysine histone marks. In order to better understand the function of these modifications we used ChIP-seq to map the genome-wide location of four short-chain acyl histone marks H3K18pr, H3K18bu, H4K12pr and H4K12bu in treated and untreated colorectal cancer (CRC) and normal cells, as well as in mouse intestines *in vivo*. We correlate these marks with open chromatin regions along with gene expression to access the function of the target regions. Our data demonstrate that propionate and butyrate bind and act as promoters of genes involved in growth, differentiation as well as ion transport. We propose a mechanism involving direct modification of specific genomic regions, by SCFA resulting in increased chromatin accessibility, and in case of butyrate, opposing effects on the proliferation of normal versus CRC cells.

## INTRODUCTION

Histone post-translational modifications (PTMs) may mediate crucial interplay between epigenetics and metabolism with important consequences for human health and disease. In addition to canonical lysine acetylation, eight types of short-chain lysine acylations have been recently identified on histones, such as propionylation (Kpr), butyrylation (Kbu), 2-hydroxyisobutyrylation (Khib), succinylation (Ksucc), crotonylation (Kcr) and β-hydroxybutyrylation (Kbhb)^1–3^. A growing body of evidence points to a unique epigenetic regulatory role for each of these modifications^4–6^. Their presence on histones is determined by the cellular metabolic state and the availability of various forms of acyl-CoA, linking metabolism to epigenetic regulation^4,7^. The cellular concentrations of non-acetyl acyl-CoAs in turn are dependent on the presence of SCFAs. Treating cells with heavy isotope-labeled SCFAs leads to heavy acyl labeling on histone proteins, pointing to a conversion of SCFAs to their cognate acyl-CoAs that are used as cofactors in histone acylation reactions^4,8–10^. This is accompanied by concomitant increases in the steady-state level of respective histone acylations in a dose-dependent manner^4,9,11^.

Recent studies show that major families of histone acetyl transferases (HATs) can catalyze histone acylation using acetyl-, propionyl- and butyryl-CoA cofactors with similar efficiencies^12–14^. For most HATs, the preference for the competing cofactor largely depends on the size of the acyl donor chain^13^. Furthermore, although almost all HATs strongly prefer acetyl-CoA, bulk levels of lysine acylations can be induced in a dose-dependent manner, in response to increasing levels of given acyl-CoA metabolites^14^. Moreover, cellular concentrations of various acyl-CoAs span orders of magnitude and are closely correlated with relative abundances of acyl marks identified *in vivo*^14^. There is also evidence that acyl-CoAs can acylate histones non-enzymatically *in vitro*^14^. These findings point to a direct link between cellular metabolism and epigenetic regulation where differential acylation is driven by cellular concentrations of respective metabolic substrates^14–16^.

Histone acetyl marks are generally associated with active regulatory elements (effects in *cis*) that promote gene expression by neutralizing Lys positive change, leading to electrostatic and structural changes in chromatin and the recruitment of readers of acylation (effects in *trans*). Chromatin immunoprecipitation followed by sequencing (ChIP-seq) experiments on Kbu, Khib, Kbhb, and Kcr show an association of active regulatory elements with histone acylations and their relative levels^2,8,9,11,17^. Recent evidence demonstrates that differential acylation states correlate with distinct physiological states and biological processes involving signal dependent gene activation, development and metabolic stress^2,8,9,11,18,19^. With respect to metabolic regulation of histone acylation, evidence points to acetyl-CoA metabolism as key determinant of histone acetylation in cancer cells, with changes in acetyl-CoA availability mediated by oncogenic metabolic reprogramming ^20^. Under the latter, the levels of acetyl-CoA will be reduced leading to ketogenesis, high NAD+/NADH ratio and activation of ACSS2, a known source of acyl-CoAs^11,21,22^. Acylation then occurs with acyl-CoAs other than acetyl-CoA.

Two SCFAs involved in histone acylation, propionate and butyrate, are generated in large quantities by microbial metabolism (as high as 70 – 100 mM in the gut lumen) and contribute to a wide array of cellular processes^23,24^. They can act as sources of energy, or as substrates for histone acylation and chromatin modification by directly targeting sites or recruiting remodeling proteins^5^. While the underlying regulatory mechanisms are largely unknown, the histone PTM state plays a key role^4,5^. Importantly, the abundance of SCFAs in the microbiome from dietary fiber metabolism makes them very attractive natural therapeutic agents, especially in the context of CRC. In cancer cells, butyrate and to a lesser extent propionate have been shown to have anti-proliferative properties that are generally attributed to inhibition of histone deacetylation (HDAC)^25,26^. According to this model, the antiproliferative, apoptotic properties of SCFAs in cancer cells are caused by histone hyperacetylation resulting from HDAC inhibition^27^.

In this study, we sought to determine the potential regulatory function of several short-chain acyl lysine histone marks, namely propionyl and butyryl H3K18 and H4K12 in CRC versus normal cells, and *in vivo*, as well as the effect of propionate and butyrate supplementation on chromatin accessibility and transcription. We also examined global acetylation levels as a function of propionate supplementation. We show that unique histone marks H3K18pr/bu and H4K12pr/bu are associated with genomic regions distinct from their acetyl counterparts. On a genome-wide level, they associate with targets controlling growth, differentiation and unfolded protein response.

Furthermore, they result in a more open chromatin structure and lead to the recruitment of a broad range of transcription factors (TFs). These results yield insights into how dietary factors, microbial metabolism and epigenetics are integrated to modulate tissue physiology and cancer susceptibility.

## RESULTS

### Identification of CRC H3K18pr/bu and H4K12pr/bu marks

We focused on H3K18 and H4K12, since acetylation at these sites has been associated with poor outcome in CRC^28–30^. To test for propionylation and butyrylation at these sites we treated CRC cells with increasing levels (0 – 10 mM) of sodium propionate (NaPr) and sodium butyrate (NaBu), consistent with physiological levels of these SCFAs in fiber-supplemented colon^31,32^ (range 1 – 100 mM). We then probed for the presence of these marks in acid-extracted histones using modification-specific antibodies. Immunoblots indicate the presence of propionylation and butyrylation on H3K18 and H4K12 at 10 mM, and in case of H3K18pr, at 1 and 10 mM propionate supplementation (Supplementary Fig. 1a, b). The results confirmed the conversion of propionate and butyrate to their cognate acyl-CoAs and their deposition as propionyl and butyryl marks on H3K18 and H4K12 in a dose-dependent manner. To further test for the presence of lysine propionylation on histones H3 and H4, we treated CRC cells with ^13^C-labeled propionate and searched for heavy peptides containing modified sites of interest by liquid chromatography-tandem mass spectrometry (LC-MS/MS).

Our results clearly indicate a direct relationship between propionate supplementation and lysine propionylation on H3 and H4, showing a steady increase in a dose-dependent manner. For example, at 10 mM supplementation, the amount of propionylation on H3K18 and H4K12 increased 1.84-fold (*P* = 0.0043) and 1.75-fold (*P* = 0.0017), respectively, compared to the control (Supplementary Fig. 1c, d). These data indicate the uptake of propionate and its deposition onto histones H3 and H4 as propionyl lysine marks. As stated earlier, major families of HATs can catalyze histone acylation such as Kpr and Kbu with similar efficiencies^12–14^. Our results demonstrate that histone acetyltransferase (HAT1), known to acetylate lysine K5 and K12 sites also mediates the incorporation of propionate into chromatin on the histone H4 K12 site. Short hairpin RNA (shRNA) induced depletion of HAT1 diminished the incorporation of propionate into H4 K12 site, compared to the control shRNA (Supplementary Fig. 1e). These findings point to a direct relationship between SCFA supplementation and histone propionylation and butyrylation at 1 and 10 mM concentrations, and more specifically, histone propionylation on H4K12 by HAT1 at 10 mM propionate supplementation.

### Acyl-CoA levels increase with SCFA supplementation

To further investigate the relationship between SCFA supplementation and the generation of the corresponding active acyl-CoA forms, we performed a quantitative metabolomics experiment using LC-MS/MS and measured the concentrations of acyl-CoA species as function of propionate and butyrate supplementation. Our results clearly indicate a dose-dependent rise in acyl-CoA levels with increasing SCFA concentrations. For example, the amount of propionyl-CoA increased from 0.08 ng/1×10^6^ cells (*P* < 0.0061) to 0.35 ng/1×10^6^ cells (*P* < 0.0001), and 0.59 ng/1×10^6^ cells (*P* < 0.0001) at 0.1 mM, 1 mM and 10 mM propionate supplementations, respectively, compared to the control (Supplementary Fig. 2a). Similarly, the amounts of butyryl-CoA increased from 0.04 ng/1×10^6^ cells (*P* = 0.0220) to 0.08 ng/1×10^6^ cells (*P* = 0.0007), at 1 mM and 10 mM butyrate supplementation, respectively (Supplementary Fig. 2b).

We then examined the levels of acetyl-CoA as a function of propionate and butyrate supplementation. With respect to propionate supplementation, our results do not show any significant increase in acetyl-CoA levels (Supplementary Fig. 2c). By contrast, butyrate supplementation led to a 2.25-fold decrease in acetyl-CoA levels at 1 mM (*P* < 0.0014), and a 3.81- fold decrease at 10 mM (*P* < 0.0002) concentrations (Supplementary Fig. 2d). While acetyl-CoA remain the dominant species in terms of absolute amounts, supplementation with propionate and butyrate results in dose-dependent increases of their cognate acyl-CoA forms, and in case of butyrate, nearly four-fold reduction of acetyl-CoA species. This points to a direct conversion of SCFA substrates to their corresponding active CoA forms, and in case of butyrate (but not propionate) a shift towards reduction of acetyl-CoA at 1 and 10 mM concentrations in SW480 CRC cells.

The effect of butyrate as an HDAC inhibitor is well established, so we decided to investigate the effect of propionate supplementation on histone acetylation levels in CRC cells. To that end, we examined the relative levels of acetylated vs unmodified states on several histone lysine sites as a function of propionate supplementation (0 – 10 mM). Our results do not show significant changes in the levels of acetylation on H3K18 or H4K12 (Supplementary Fig. 3a, b). The most abundant states on H3 and H4 were the unmodified ones, with a ratio of the unmodified to acetylated states approximately 10:1.

To test whether SCFA supplementation resembled HDAC inhibition we compared treatment of CRC cells with increasing levels of propionate, butyrate and trichostatin A (TSA), a potent HDAC inhibitor. TSA treatment led to loss of cell viability at 0.1 μM and near complete cell death at 1 μM treatment (*P* < 0.0001) (Supplementary Fig. 3c). Butyrate, with its much weaker HDAC activity compared to TSA, resulted in reduced cell viability only at 10 mM supplementation (*P* = 0.0002). Propionate on the other hand, did not affect cell viability in the 0 – 10 mM range but only at 100 mM (*P* < 0.0001). Both propionate and butyrate supplementations resulted in near complete loss of cell viability at 100 mM concentrations. Because of the toxicity apparent at high levels, our experiments were performed with 10 mM propionate and 1 mM butyrate to avoid cytotoxicity artifacts.

### Genomic localization of H3K18pr and H4K12pr in CRC cells

We next focused on the genome-wide distribution of H3K18pr and H4K12pr. Given the close coupling of acetylation, propionylation and butyrylation, we have compared differential Kpr binding to the corresponding acetyl marks. Results showed that out of 19,167 sites identified as differentially bound after propionate supplementation, 17,299 (90%) were associated with H3K18pr versus 1,868 associated with H3K18ac (FDR < 0.05) (Fig. 1a, b, g). Gene Ontology (GO) analysis of Kpr-bound regions and distal *cis*-regulatory elements^33^ pointed to enrichment in epidermal growth factor stimulus and cadherin binding, as well as endoplasmic reticulum (ER) unfolded protein response (Fig. 1c). KEGG pathway analysis also showed enrichment focal adhesion (110), actin cytoskeleton (115) and cancer regulatory pathways (Supplementary Fig. 4a, b). We focused on enrichment of CRC-relevant motifs such as SMAD2/3 of TGF-β pathway, as well as AP-1, FOSL2 and JUNB. All four motifs showed enrichment in K18pr over K18ac and input, with JUNB, FOS2L and AP-1 exhibiting greater than a six, seven and five-fold enrichment over background, respectively (Fig. 1e). Differential binding of key Wnt/β-catenin pathway genes showed ∼3-fold enrichment in *CTNNB1*, *TCF20*, *LEF1* (Supplementary Fig. 4c). We also observed a 2-3-fold enrichment in *FOS* and *JUN*. Distribution of reads over all differentially bound sites showed Kpr as having a higher mean read concentration compared to Kac (Fig. 1d). Hierarchical clustering of Kpr annotated genes showed several clusters including cell projection organization and localization, while chromosomal positions and distribution by gene type showed enriched regions on chromosomes 3, 9, 17, 19, as well as the presence of lncRNAs and processed pseudogenes (Supplementary Fig. 4d, e).

**Figure 1:**
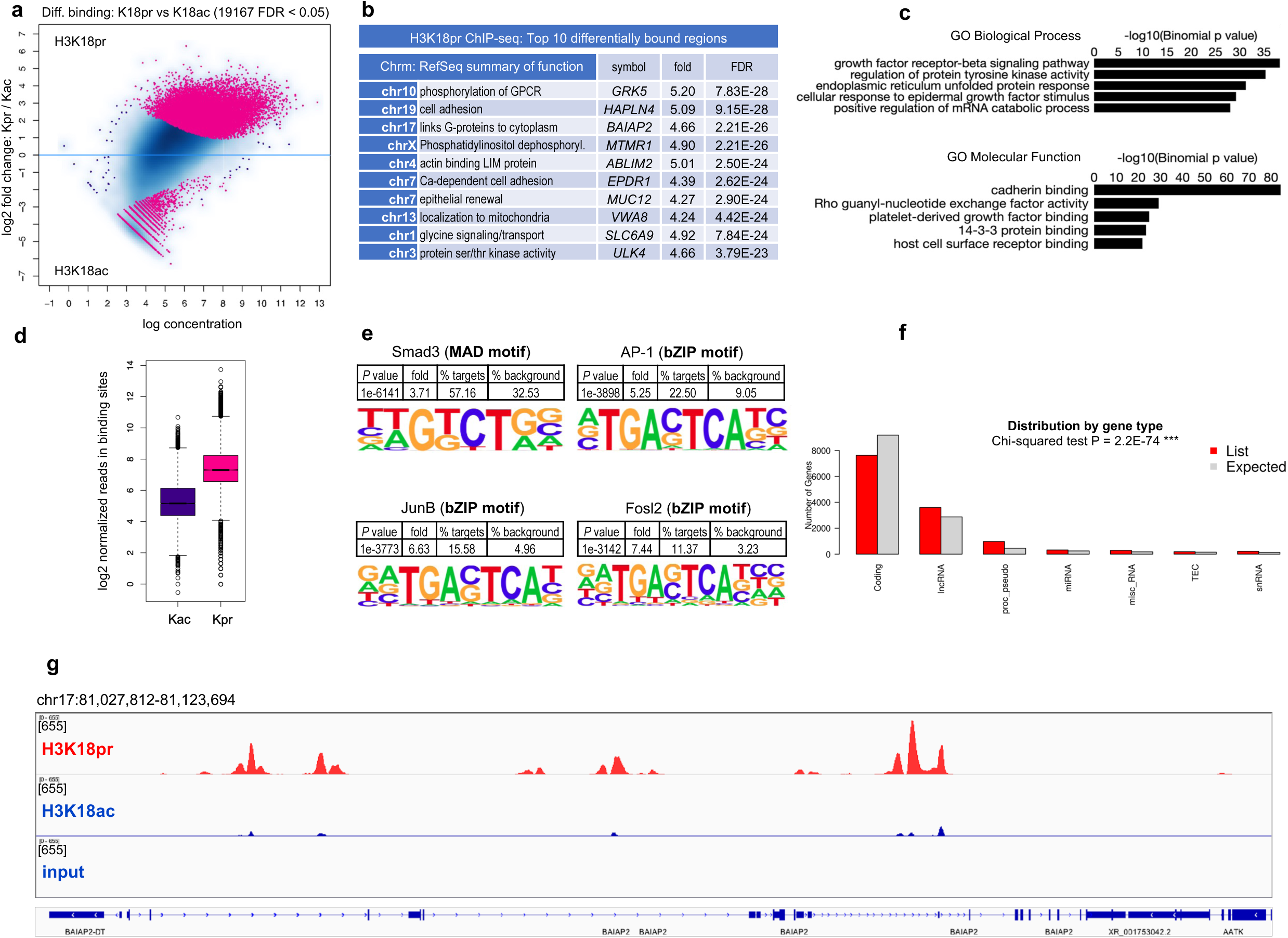
Genome-wide H3K18pr distribution. **a.** H3K18pr vs H3K18ac differential binding at 10 mM propionate treatment. Sites identified as significantly differentially bound are shown in red. n = 3 technical replicates for each condition. Differential binding was performed by DiffBind package with DESeq2 using a two-sided test for both increased and decreased binding affinity between conditions followed by multiple hypothesis testing and FDR correction. **b.** Top ten differentially bound regions associated with H4K12pr, annotated to within 1 Kb of TSS, sorted by false-discovery rate adjusted *P* value (FDR < 0.05). **c.** Top GO Biological Process and Molecular Function terms of H3K18pr-associated *cis*-regulatory elements (5+ 1 Kb) determined by GREAT against a whole genome background using a binomial test over genomic regions, followed by multiple hypothesis testing using FDR corrected *P* values (FDR < 0.05). **d.** Normalized reads in H4K12pr vs H4K12ac-associated binding sites at 10 mM propionate treatment. Box plots display: The minimum, first quartile (Q1, 25^th^ percentile), median, third quartile (Q3, 75^th^ percentile), and maximum. The bottom of the box is Q1 and the top of the box is Q3. The line within the box represents the median (50^th^ percentile) value. The whiskers extend to the most extreme data points within 1.5 times the IQR (interquartile range). **e.** Differential motif analysis of H3K18pr vs H3K18ac peaks was analyzed by HOMER, using a one-sided hypergeometric test for overrepresentation (enrichment) of motifs in the target sequences compared to the background, followed by multiple hypothesis testing and FDR correction. **f.** Distribution of H3K18pr peaks by gene type was measured by two-sided Chi-squared test to assess whether the observed distribution significantly differs from the control distribution, without specifying a direction (enrichment or depletion) followed by multiple hypothesis testing and FDR correction. **g.** Signal tracks for regions representing *BAIAP2*. Signal intensity of peaks in 95 Kb-spanning *BAIAP2* region showing H3K18pr vs H3K18ac binding at 10 mM propionate treatment with input as background.

H4K12pr ChIP-seq identified 28,465 sites as differentially bound between Kpr and Kac, with 27,175 (95%) sites associated with Kpr (FDR < 0.05) (Fig. 2a, b, g). Genomic regions annotation pointed to enrichment in genes controlling Rho-guanyl-nucleotide exchange activity and differentiation, cadherin and ER protein binding, as well as calcium channel signaling (Fig. 2b). As with H3K18pr, GO and KEGG analysis pointed to pathways controlling platelet-derived growth factor and cadherin binding, ER unfolded protein response, and Ca^2+^ channel activity (Fig. 2c, Supplementary Fig. 5a, b). Motif analysis also showed enrichment in the SMAD family and Krüppel-like factors KLF5, KLF14 known for their regulatory role as recruiters of other TFs (Fig. 2e). Hierarchical clustering showed several clusters such as cellular catabolic processes, protein transport and cellular localization, while chromosomal positions and distribution by gene type showed enrichment along chromosomes 7, 9, 11, 17, as well as the presence of long non-coding RNAs (lncRNAs) and processed pseudogenes (Supplementary Fig. 5d, e, Fig. 2f). H4K12pr peak distributions, regulated genes, GO enrichments analysis showed generally similar results with H3K18pr analyses, including prioritization of Wnt/β-catenin, TGF-β and FOS, JUN, MYC TFs (Figs. 2a, c, e, Supplementary Fig. 5c).

**Figure 2:**
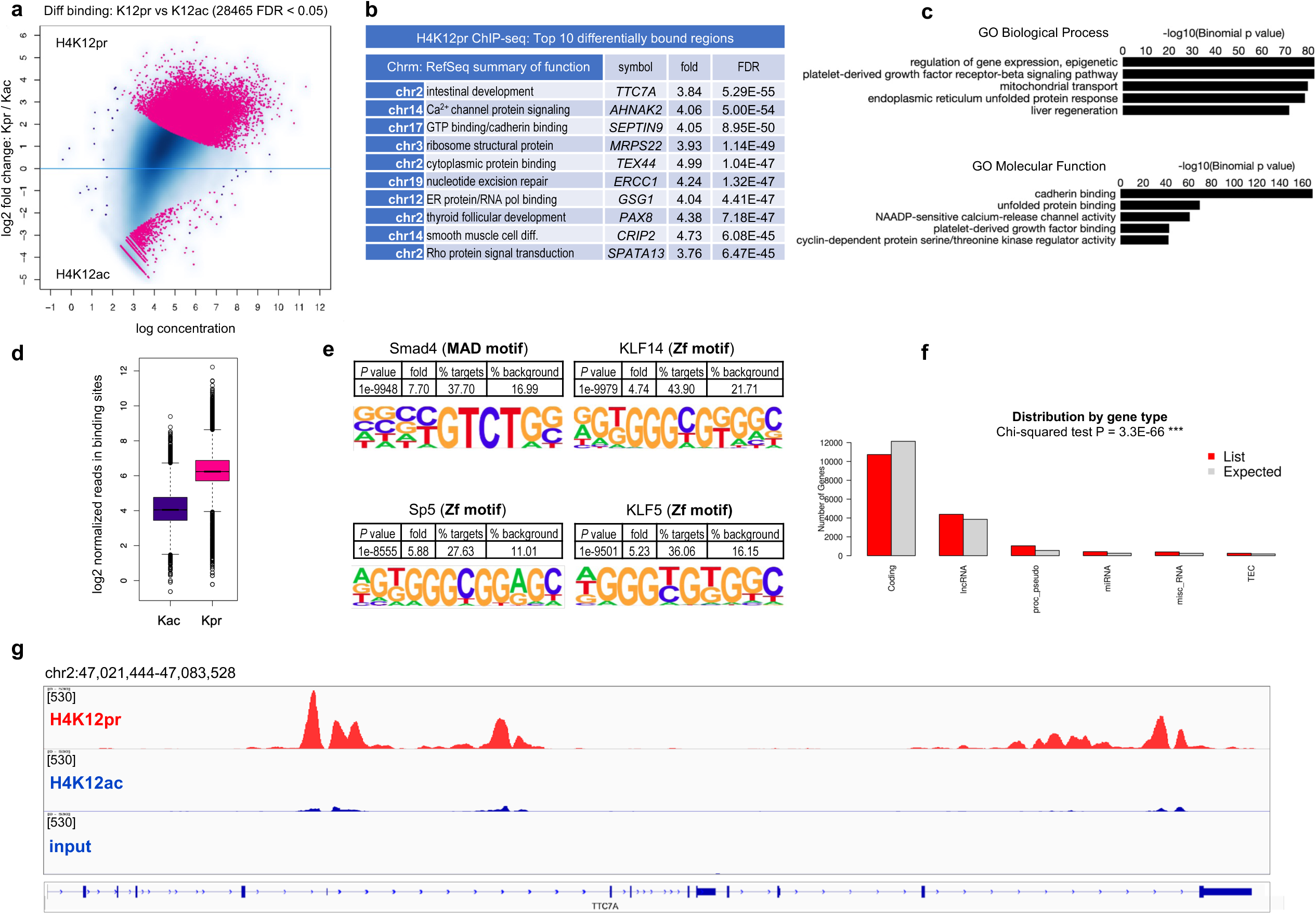
Genome-wide H4K12pr distribution. **a.** H4K12pr vs H4K12ac differential binding at 10 mM propionate treatment. Sites identified as significantly differentially bound are shown in red. n = 3 technical replicates for each condition. Differential binding was performed by DiffBind package with DESeq2, using a two-sided test for both increased and decreased binding affinity between conditions, followed by multiple hypothesis testing and FDR correction. **b.** Top ten differentially bound regions associated with H4K12pr, annotated to within 1 Kb of TSS, sorted by false-discovery rate adjusted *P* value (FDR < 0.05). **c.** Top GO Biological Process and Molecular Function terms of H4K12pr-associated *cis*-regulatory elements (5+ 1 Kb) determined by GREAT against a whole genome background using a binomial test over genomic regions, followed by multiple hypothesis testing using FDR corrected *P* values (FDR < 0.05). **d.** Normalized reads in H4K12pr vs H4K12ac-associated binding sites at 10 mM propionate treatment. Box plots display: The minimum, first quartile (Q1, 25^th^ percentile), median, third quartile (Q3, 75^th^ percentile), and maximum. The bottom of the box is Q1 and the top of the box is Q3. The line within the box represents the median (50^th^ percentile) value. The whiskers extend to the most extreme data points within 1.5 times the IQR (interquartile range). **e.** Differential motif analysis of H4K12pr vs H4K12ac peaks was analyzed by HOMER, using a one-sided hypergeometric test for overrepresentation (enrichment) of motifs in the target sequences compared to the background, followed by multiple hypothesis testing and FDR correction. **f.** Distribution of H4K12pr peaks by gene type was measured by two-sided Chi-squared test to assess whether the observed distribution significantly differs from the control distribution, without specifying a direction (enrichment or depletion) followed by multiple hypothesis testing and FDR correction. **g.** Signal tracks for regions representing *TTC7A*. Signal intensity of peaks in 95 Kb-spanning *TTC7A* region showing H4K12pr vs H4K12ac binding at 10 mM propionate treatment with input as background.

We then integrated our Kpr ChIP-seq results with assay for transposase-accessible chromatin followed by sequencing (ATAC-seq) and RNA-seq results in CRC cells to determine intersecting genomic coordinates and overlapping annotated genes. Genomic coordinates present in both Kpr ChIP-seq and propionyl ATAC-seq data sets (n = 4391*, P* = 4.75e-4 and n = 4038, *P* = 2.31e-38, respectively) showed enrichment in β-catenin-TCF complex assembly, negative regulation of MAPK cascade, and actin filament organization (Supplementary Fig. 6a, b, e, f). There was also enrichment in Rho guanyl-nucleotide exchange factor activity and proteins involved in cell adhesion. Genes relevant to the identified pathways, and CRC in particular, also showed increased accessibility by ATAC-seq, indicating more open chromatin structure following propionate supplementation (Supplementary Fig. 6c, d, g, h). Integration with propionyl RNA-seq showed overlap between both H3K18pr and H4K12pr targets and upregulation (n = 1528, *P* = 7.10e-154 and n = 1585, *P* = 1.69e-147), as well as downregulation (n = 1015, *P* = 3.29e-12 and n = 1317, *P* = 1.35e-167) of gene expression (Supplementary Fig. 7a, c, e, g). GO analysis of Kpr targets and upregulated genes identified pathways in differentiation, localization as well as actin filament-based processing, while downregulated genes were involved in mitotic cell cycle, RNA metabolism and processing (Supplementary Fig. 7b, d, f, h). This points to a mechanism through which propionate affects the CRC epigenetic regulatory landscape to promote growth, differentiation, and localization over regulation of cell cycle and RNA processing.

### Genomic localization of H3K18bu and H4K12bu in CRC cells

We next examined Kbu marks on the same sites to gain functional insight and observe butyrate supplementation (1 mM) effects on chromatin structure and accessibility. Butyrate has been shown to activate the TGF-β pathway^34^ and the Wnt signaling pathway genes in CRC in particular^35^. ChIP-seq experiments on Kbu on H4K5 and H4K8 in the context of sperm cell differentiation show histone butyrylation as a direct stimulator of transcription, while also competing with acetylation in chromatin reorganization^4,36^. H3K14bu ChIP-seq in mouse livers was recently shown to be associated with transcriptionally active chromatin and specifically with carboxylic acid and lipid metabolism^37^.

Compared to Kpr, fewer differentially bound regions were associated with H3K18bu and H4K12bu: 2305 and 793, respectively (Supplementary Fig. 8a, 9a). Interestingly, top scoring sites associated with both Kbu marks were lncRNAs *PRNCR1* and *PCAT1*, enhancers in the same chromosomal region as *MYC*, shown to play a pivotal oncogenic role in CRC^38,39^. *PRNCR1* showed a 3.4-fold (FDR = 2.25e-09) and 2.9-fold (FDR = 3.14e-77) increase in H3K18bu and H4K12bu binding, respectively (Supplementary Figs. 8b, f, 9b), whereas *PCAT1* showed a 4.19-fold increase in H3K18bu (FDR = 6.79e-16) and a 2.8-fold increase in H4K12bu binding (FDR = 1.57e-12). GO enrichment, differential motif analysis and integration with butyryl ATAC-seq results identified pathways similar to Kpr and propionyl ATAC-seq (Supplementary Figs. 8c-e, 9c-f, 10d-g).

Next, we visualized the read coverage over genomic regions in 1 kB upstream and downstream of TSS for all our acyl lysine histone marks to evaluate global enrichment across all TSS (Fig. 3). In each case, we observed considerable enrichment of our marks proximal to the TSS and beyond, with increases in read density over input proportional to the length of the acyl lysine chain. Propionyl and butyryl marks showed significantly higher density distributions in the positive, downstream of TSS direction compared to their acetyl counterparts. All three types of acyl marks showed consistency in the distribution profiles (Fig. 3, upper panels) and heatmap densities (Fig. 3, bottom panels). These results indicate that Kpr and Kbu marks increased chromatin accessibility, compared to Kac. Feature distribution of differentially bound genes (+/- 3 kB of TSS) also showed increased chromatin accessibility and consistency in feature distributions among the three types of acyl marks, with a shift towards distal intergenic regions in Kpr and Kbu (Extended Fig. 1). These results indicate a greater role played by *cis*-regulatory elements and a more open chromatin structure.

**Figure 3:**
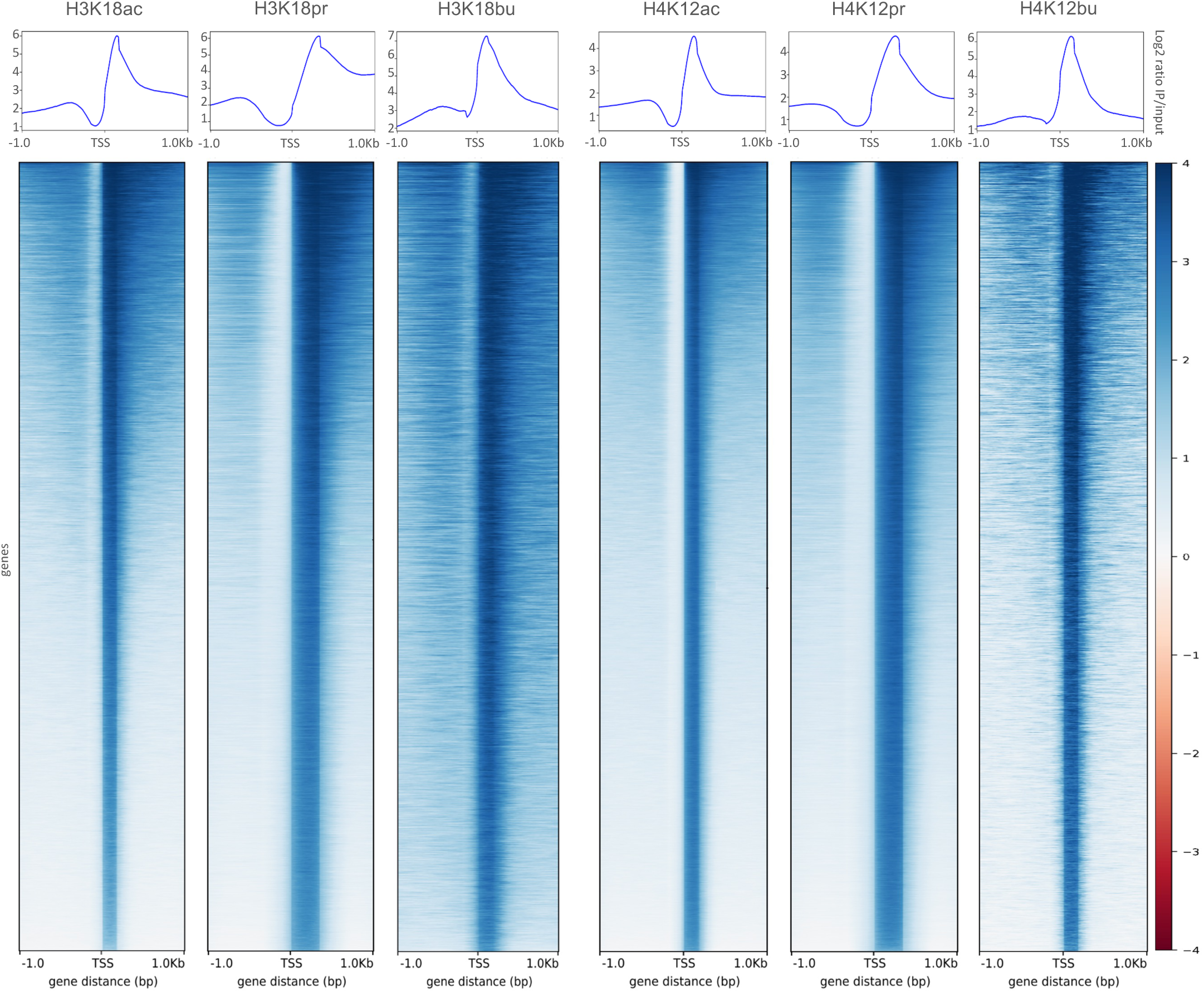
TSS distribution profiles of H3K18ac/pr/bu and H4K12ac/pr/bu associated ChIP-seq peaks as function of read coverage. *Upper panels*: Aggregate read density profile plots of genomic region distributions within +/- 1 Kb of TSS as a function of log2 IP/input ratio. *Lower panels*: Read density heatmaps of gene distributions with maximum (z = 4) and minimum (z = - 4) values of heatmap intensities. Plots generated by deepTools.

Lastly, we examined functional differences between the three types of acyl marks by examining annotated genes that were unique to each (Extended Fig. 2). Terms unique to Kac marks pointed to regulation of cell cycle and chromosome organization (Extended Fig. 2a, d). By contrast, terms unique to Kpr pointed to regulation of cation transport, organ morphogenesis and locomotion (Extended Fig. 2b, e). Elements unique to Kbu were associated almost exclusively with regulation of cell motility, migration and locomotion (Extended Fig. 2c). The results indicate that while there are some functional similarities between Kpr and Kbu marks, they are distinct from Kac. Comparison of shared annotated features between Kpr/Kbu marks and ATAC/RNA-seq results revealed many of the same pathways, highlighting their functional similarities as well as their differences (Extended Fig. 3).

### SCFA increase chromatin accessibility in CRC cells

To address the effect of propionate and butyrate on chromatin accessibility at a more global level, we performed differential ATAC-seq. We hypothesized that propionate and butyrate supplementation would result in greater chromatin accessibility. Out of a total of 22,238 regions identified as differentially accessible, 18,404 (83%) sites showed positive fold-change in the propionate-treated group compared to 3,834 in the untreated group (FDR < 0.05) (Supplementary Fig. 10a, top). Differentially accessible genes and GO pathways associated with propionate treatment were consistent with Kpr ChIP-seq results, particularly with respect to genes controlling cell-substrate adhesion (*COL26A1*, 2.82-fold, FDR = 8.78e-37) as well as epithelial development (*KLF2*, 3.39-fold, FDR = 6.09e-37) and β-catenin-TCF complex assembly (Supplementary Fig. 10b, top). Sites that showed a reduction in accessibility were involved in cell cycle G1 arrest (*CDKN1A*, −1.78-fold, FDR = 9.04e-14), as well regulation of cell cycle and cell proliferation (*ZNF703,* −1.74-fold, FDR = 1.41e-11) (Supplementary Fig. 10a, top). Among differentially bound sites overall, the propionate-treated group had a higher mean read concentration, indicating increased binding affinity under more open chromatin (Supplementary Fig. 10c, top).

By contrast, over three times as many sites were identified with butyryl ATAC-seq compared to propionyl ATAC-seq. Out of 71,318 sites identified as differentially accessible, 38,089 (53%) showed positive fold-change in the treated group and 33,229 underwent negative fold-change in the untreated group (Supplementary Fig. 10a, bottom). Genomic annotations identified genes involved in muscle contraction (*SSPN,* 2.31-fold, FDR = 5.41e-136), axon guidance and differentiation (*SEMA3D*, 2.38-fold, FDR = 5.16e-131), as well as actin and microtubule binding (*MYO3B*, 2.41-fold, FDR = 8.11e-99) (Supplementary Fig. 10b, bottom). GO pathway analysis showed enrichment in mesenchymal cell proliferation, metanephros development and fibroblast apoptotic process, as seen with Kbu ChIP-seq (Supplementary Fig 10b, bottom, Supplementary Fig. 8c, 9c).

Among sites that lost accessibility were genes encoding a zinc finger TF required for DNA damage-induced p53 activation (*CXXC5*, −2.06-fold, FDR = 1.26e-187) and a G protein subfamily member mediating transmembrane signaling (*GNAZ*, −3.04-fold, FDR = 4.76e-160). Among differentially accessible sites overall, the butyrate-treated group did not have a significantly higher mean read concentration, indicating increased accessibility in the treated group was offset by decreased accessibility in the untreated group (Supplementary Fig. 10c, bottom).

### SCFA affect gene expression in CRC cells

To gain a more complete picture of SCFA induced alterations on the regulatory landscape of CRC, we examined global changes in gene expression by bulk RNA-seq. Differential gene expression analysis following propionate supplementation identified 2,027 upregulated and 1,151 downregulated genes (Fig. 4a). Among the genes that showed significant upregulation, and were also identified as Kpr targets and exhibited increased accessibility by ATAC-seq were *SAT1* (2.99- fold FDR = 2.04e-100), *AHNAK2* (2.98-fold, FDR = 8.71e-71), *DHRS2* (3.80-fold, FDR = 2.44e-67), *KLF2* (3.28-fold, FDR = 4.95e-32) as well as *MYC* and *FOS* (Supplementary Fig. 4c, 5c). Among genes that underwent downregulation were mainly those involved in cell proliferation (*ANP32B*, −2.63-fold, FDR = 2.01e-52) and cell cycle progression (*MKI67*, −1.57-fold, FDR = 5.90e-48). Hierarchical clustering showed upregulation in receptor signaling, development, anatomical structure morphogenesis, and downregulation in cell cycle, cell division, and chromatin organization (Fig. 4b). The heatmap of top 50 most variable genes showed enrichment in *MYC, JUN* and *AHNAK2* (Fig. 4c).

**Figure 4:**
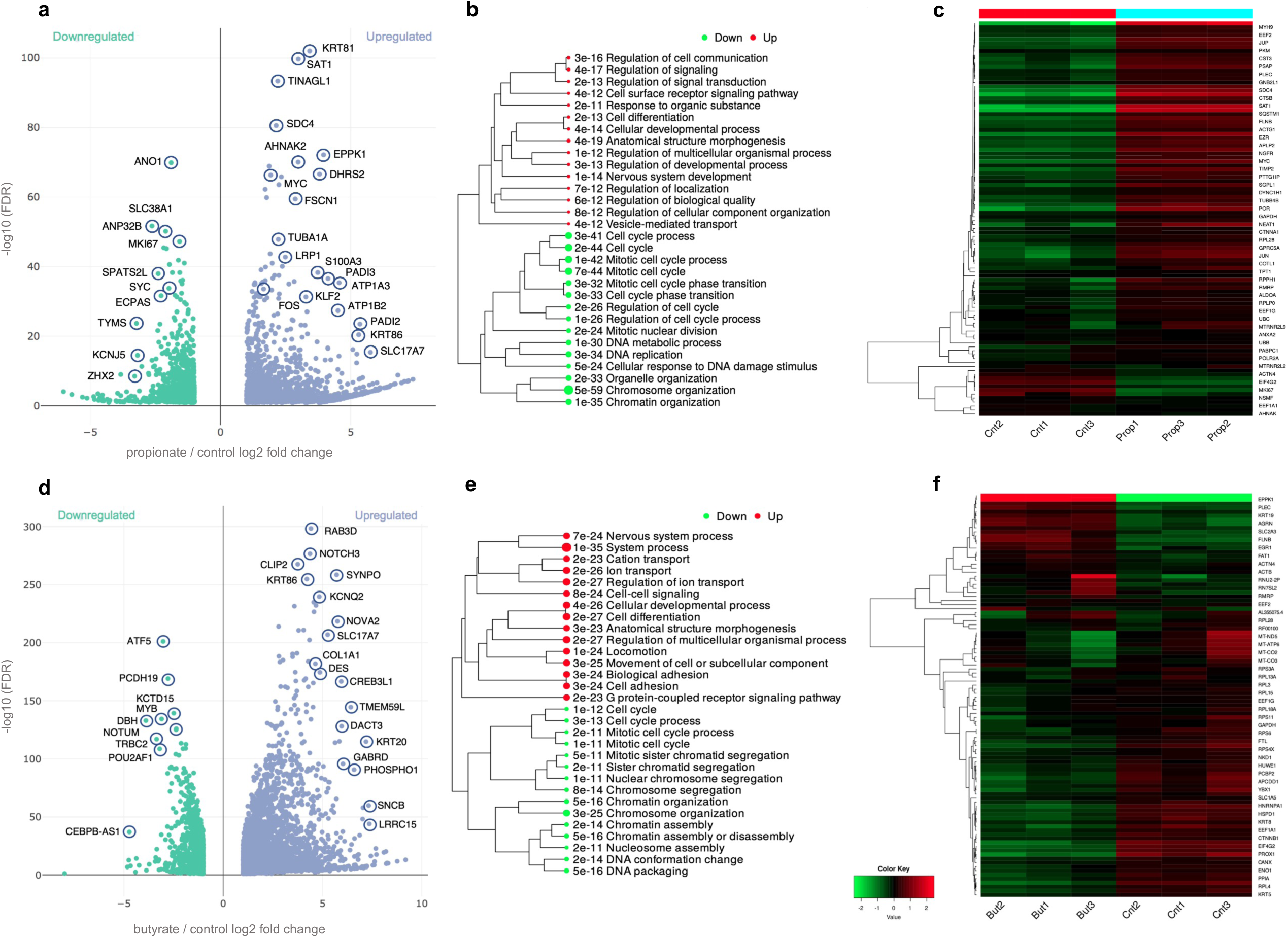
Propionyl and butyryl differential gene expression by RNA-seq. **a.** Volcano plot showing gene upregulation vs downregulation in 10 mM propionate treated vs control groups. n = 3 technical replicates for each condition. Differential expression analysis performed by DESeq2 **b.** Hierarchical cluttering of GO ‘Biological Process’ terms of upregulated vs downregulated pathways in propionate treated vs control groups. Hierarchical clustering of the pathways was performed using ShinyGO. Pathways were clustered together based on shared genes and gene enrichment analysis was performed using two-sided Fisher’s exact test, and FDR correction was applied to adjust for multiple comparisons in the pathway analysis and hierarchical clustering. Size of dots indicates statistically significant FDR adjusted (FDR < 0.05) *P* values. **c.** Heatmaps of 50 most variable genes in propionate treated vs control groups. **d.** Volcano plot showing gene upregulation vs downregulation in 1 mM butyrate treated vs control groups. **e.** Hierarchical cluttering of GO ‘Biological Process’ terms of upregulated vs downregulated pathways in butyrate treated vs control groups. **f.** Heatmaps of 50 most variable genes in butyrate treated vs control groups. Hierarchical clustering was performed using ShinyGO.

Similarly, butyrate supplementation led to upregulation of genes controlling ion transport, anatomical structure morphogenesis and cell adhesion, and downregulation in cell cycle and chromatin assembly (Fig. 4d-f). Taken together, our data point to a mechanism whereby the antiproliferative properties of propionate and butyrate in CRC can be attributed to their dysregulation of key CRC oncogenes such as *MYC*, *FOS* and *JUN*, as well their simultaneous triggering of downregulation of genes controlling cell cycle and cell division.

To examine the differences in expression that were unique to propionate and butyrate we compared differential gene expression under both types of enrichments. Density distributions of counts were consistent across conditions and replicates within each condition (Fig. 5a). Principal component analysis (PCA) showed close clustering of replicates within each condition, with the most variation seen between different conditions forming separate clusters (Fig. 5b). Differential expression following propionate versus butyrate supplementation showed 3,082 upregulated and 2,783 downregulated genes (Fig 5c). The heatmap of top 50 most variable genes and hierarchical clustering of GO ‘Biological Process’ terms showed preferential enrichment of organic and carboxylic acid metabolism associated with butyrate supplementation, while propionate supplementation showed enrichment in cell motility and locomotion, as well as cellular development and differentiation (Fig. 5d, e).

**Figure 5:**
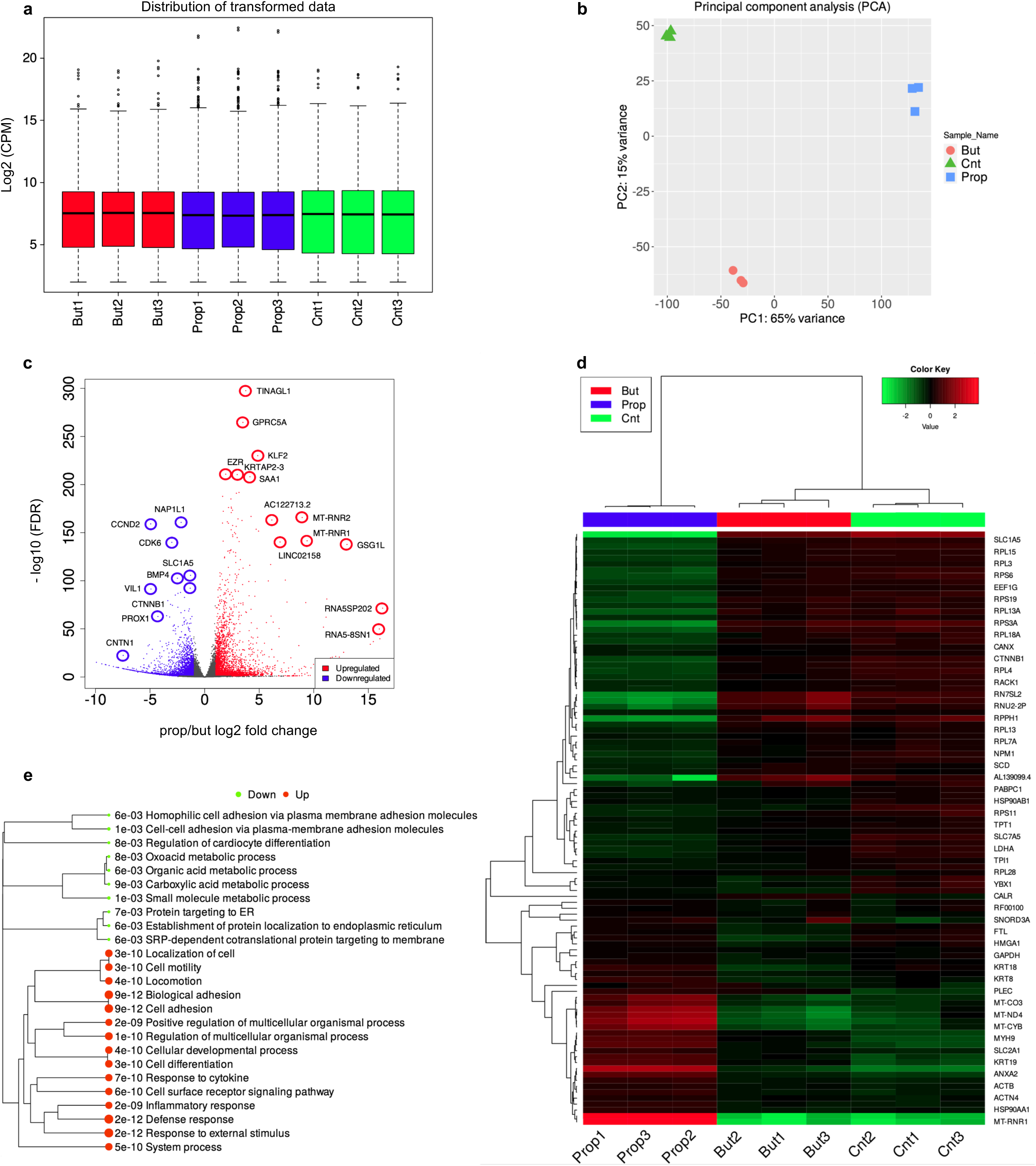
Propionyl/butyryl differential RNA-seq. **a.** Distribution of log2 CPM (counts per million) transformed expression data for all conditions. n = 3 technical replicates for each condition. Normalization of raw counts performed by ‘cpm’ analysis in edgeR. Box plots display: The minimum, first quartile (Q1, 25^th^ percentile), median, third quartile (Q3, 75^th^ percentile), and maximum. The bottom of the box is Q1 and the top of the box is Q3. The line within the box represents the median (50^th^ percentile) value. The whiskers extend to the most extreme data points within 1.5 times the IQR (interquartile range). **b.** Principal component analysis of log2 CPM transformed expression data for all conditions. **c.** Volcano plot of propionyl vs butyryl differential expression. **d.** Heatmaps of 50 most variable genes for all three conditions. **e.** Hierarchical cluttering of GO ‘Biological Process’ terms of differentially expressed pathways in propionyl vs butyryl RNA-seq. Hierarchical clustering of the pathways was performed using ShinyGO. Pathways were clustered together based on shared genes and gene enrichment analysis was performed using two-sided Fisher’s exact test, and FDR correction was applied to adjust for multiple comparisons in the pathway analysis and hierarchical clustering. Size of dots indicates statistically significant FDR adjusted (FDR < 0.05) *P* values.

To address the osmolarity of the sodium counterion and potential off-target effects of sodium resulting from NaPr and NaBu supplementation at 1 mM and 10 mM concentrations used in our differential chromatin accessibility by ATAC-seq and differential expression by RNA-seq data, we performed those experiments in the presence and absence of 1 mM and 10 mM sodium chloride (NaCl), rather than NaPr and NaBu. Differential accessibility by ATAC-seq at 1 mM NaCl supplementation vs control did not generate sufficient contrast between two conditions for differential expression analysis. At 10 mM NaCl vs control, 26 sites were identified (FDR < 0.05, log2 FC > 1) as differentially accessible with only 4 regions showing positive fold change in accessibility as a result of 10 mM NaCl supplementation (Extended Fig. 4a). Three of those regions showed only very modest fold change (log2 1.08 - 1.11, FDR = 0.0012 – 00029) with only one region showing 3.15-fold change in accessibility (FDR = 2.06e-29).

Annotation of differentially bound regions at 10 mM NaCl supplementation identified this region 12q13.13 mapping to *HOX6,* a member of a homeobox family of transcription factors, and an adjacent ncRNA gene *MIR196A2I,* affiliated with the miRNA class (3.15-fold, FDR = 2.06e-29) (Extended Fig. 4b). The regions that underwent negative fold-change (or were preferentially enriched in the control group) upon 10 mM NaCl supplementation beyond the significance threshold were *SLC1A5* (−1.08-fold, FDR = 1.81e-14) and *SLC43A* (−1.04-fold, FDR = 4.13e-10) genes encoding sodium-coupled neutral amino acid transporters; *TBC1D16* (−1.01-fold, FDR = 1.20e-09), a TBC1 domain family member involved in GTPase activator activity and regulation of receptor recycling; and *LCN9* (−1.1-fold, FDR = 3.44e-06), a member of the lipocalin family involved in binding and transport of small hydrophobic ligands. Overall, at 10 mM NaCl supplementation, the number of differentially accessible regions by ATAC-seq that showed positive fold change or enrichment over the control beyond the significance threshold was insufficient to account for any potential off-target effects due to sodium in NaPr and NaBu supplementation.

Similarly, differential expression analysis by RNA-seq in the presence and absence of 1 mM and 10 mM NaCl did not result in any significant contrast between NaCl and control conditions (Extended Fig. 4c, d). No genes were identified beyond the significance threshold at 1 mM vs control (Extended Fig. 4c, d, insert), and only that two that showed slight preferential enrichment in the 10 mM NaCl condition; *DCBLD2* (0.31-fold, FDR = 0.00016), a gene involved in negative regulation of cell growth and wound healing; and *TGM2* (0.22-fold, FDR = 0.01147), a gene involved in GTP binding and protein-glutamine gamma-glutamyltransferase activity. Genes showing negative fold-change in expression were *JUNB* (−0.45-fold, FDR = 0.00016), and *HR* (- 0.54-fold, FDR = 0.000838); genes involved in DNA-binding transcription factor activity and transcription corepressor activity.

Since all immunoprecipitation-based experiments were carried out under the same NaPr and NaBu enrichment conditions as the controls, and only acyl-lysine antibodies were compared, the effects of the sodium counterion were thus normalized across all samples and conditions in our Kpr/Kbu ChIP-seq and CUT&Tag experiments. Therefore, only ATAC-seq and RNA-seq experiments that were carried out at different treatment conditions were further investigated for potential off-target effects due to the sodium counterion. Based on the significance threshold (FDR < 0.05, log2 FC > 1) for differential accessibility and expression, we did not see evidence of off-target effects due to sodium that would affect the results in our ATAC-seq and RNA-seq experiments. Furthermore, the RNA-seq correlation heatmap of 1 mM and 10 mM NaCl groups vs the control showed clustering of replicates under different conditions (Extended Fig. 4e). Thus, there is not a significant statistical difference in expression between 1 mM, 10 mM NaCl supplemented groups and the control in SW480 cells.

### Genomic localization of Kbu marks in normal vs cancer cells

We next examined the differences in genome-wide distribution of Kbu marks in CRC (SW480) versus normal (CCD841) cells following 1 mM butyrate supplementation, particularly in relation to CRC-relevant genes. Out of 89,340 targets associated with H3K18bu differential binding in the two cell lines, 81,999 (92%) sites had higher binding affinity in normal cells, compared with 7,341 sites in cancer cells (FDR < 0.05) (Extended Fig. 5a). GO ‘Biological Process’ analysis in the normal cell line showed enrichment in cell-substrate and adherens junction assembly, as well as ion transport and protein processing (Extended Fig. 5b, top). By contrast, the same mark in the cancer cell line showed enrichment in mesenchymal cell proliferation, β-catenin-TCF complex assembly, and fibroblast apoptotic process (Supplementary Fig. 15b, bottom). Similarly, out of 64,886 targets associated with H4K12bu, 61,978 (96%) sites had higher binding affinity in normal cells, compared to 2,908 sites in cancer cells (Extended Fig. 5c). GO ‘Biological Process’ analysis in normal cells also showed enrichment in adherens junction assembly and transport but also regulation of chromatin silencing and H3K9 demethylation (Extended Fig. 5d, top). By contrast, GO analysis in the cancer cell line showed enrichment in mesenchymal cell proliferation, Wnt signaling pathway, and ER stress-induced apoptotic signaling (Extended Fig. 5d, bottom).

While both marks were associated with adherens and cell junction assembly, in SW480 cancer cells they were associated with mesenchymal cell proliferation, Wnt/β-catenin signaling, as well as apoptotic processes, while in normal cells they showed association with transmembrane ion and endosome to lysosome transport, as well as protein processing. Moreover, following butyrate supplementation, many of the CRC-relevant genes monitored throughout the study, such as *MYC* and *FOSL1* showed a 3 to 7-fold reduction in Kbu binding affinity in the normal cell line compared to the CRC cell line (Extended Fig. 5e-h).

### Genomic localization of Kbu marks in mouse intestines

To further investigate the link between dietary fiber metabolism, chromatin accessibility and histone butyrylation, we performed ATAC-seq in CT26 mouse colorectal cells and cleavage under targets and tagmentation (CUT&Tag) on large intestine tissues from mice fed chow containing the dietary fiber arabinoxylan (AX, 5% w/w) (see Methods). First, we examined differential chromatin accessibility following 1 mM butyrate supplementation in CT26 murine CRC cells. Out of 54,171 sites identified as differentially accessible, 39,956 (74%) gained accessibility following butyrate supplementation, compared to 14,215 sites in the untreated group (FDR < 0.05) (Fig. 6a). Peak distributions, regulated genes and GO enrichments analysis showed similar results to those in SW480 and CCD841 cell lines (Fig. 6a-e). Differential Kbu binding analysis of mouse intestines on a 5% arabinoxylan diet by CUT&Tag identified 21,665 sites associated with H3K18bu versus Kac, and 26,103 sites associated with H4K12bu versus Kac (FDR < 0.05) (Extended Fig. 6a, b, d, e). GO analysis and feature distributions were in agreement with Kbu ChIP-seq results in SW480 cells (Extended Figs. 6c, f, g, h, 11). Annotation of top H3K18bu-associated regions identified genes involved in cell proliferation and differentiation (*Fgf14*, 4.60-fold, *P* = 1.61e-05) as well as ion channel regulatory activity (*Akap9,* 5.72-fold, *P* = 3.55e-05) and actin filament polymerization (*Cyria,* 5.86-fold, *P* = 3.92e-05) (Extended Fig. 6b). Similarly, top H4K12bu-associated regions were involved in fibroblast growth factor receptor activity (*Fgfrl1,* 6.97-fold, *P* = 5.33e-06) as well as tight junction cell adhesion activity (*Cldn3*, 5.72-fold, *P* = 5.33e-06) (Extended Fig. 6e). H3K18bu GO analysis identified pathways controlling cell-substrate junction assembly, cytoskeletal organization as well as autophagosome assembly (Fig. 6g). H4K12bu GO analysis identified pathways controlling protein processing, membrane localization, as well as TGF-β production (Fig. 6h). Comparing mouse butyryl ATAC-seq in cells following butyrate supplementation to H3K18bu targets *in vivo* revealed 9,221 (72%) overlapping elements out of 13,867 (*P* = 6.41e-05) (Fig. 6i). Comparing mouse butyryl ATAC-seq results following butyrate supplementation to H4K12bu targets *in vivo* revealed 10,243 (64%) overlapping elements out of 16,046 (*P* = 5.81e-55) (Fig. 6j).

**Figure 6:**
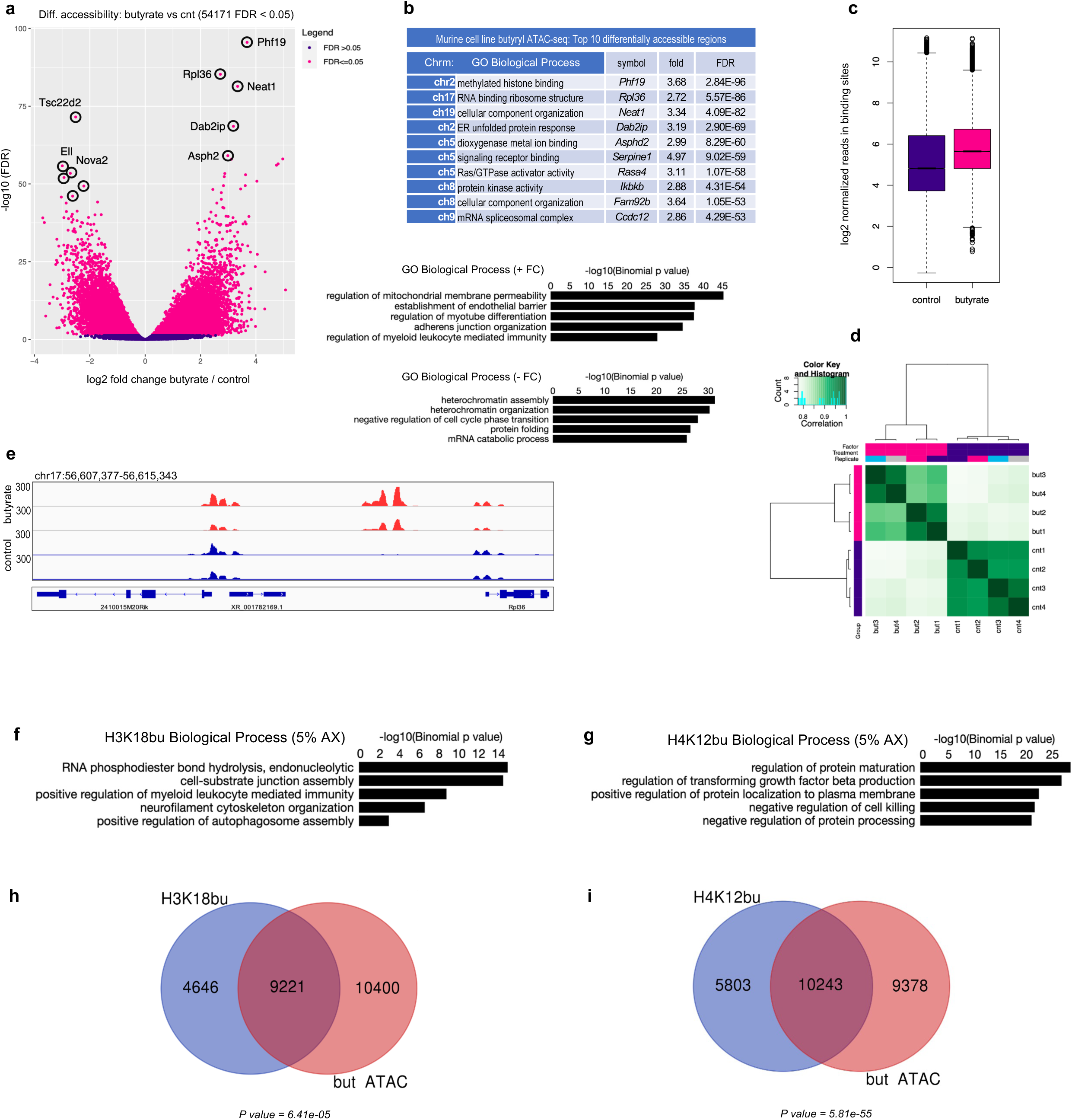
Murine cell line butyryl ATAC-seq and Kbu associated targets in mouse intestines. **a.** Differential accessibility at 1 mM butyrate supplementation. Sites identified as significantly differentially accessible are shown in red. n = 4 technical replicates for each condition. Differential accessibility was performed by DiffBind package with DESeq2 using a two-sided test for both increased and decreased binding affinity between conditions followed by by multiple hypothesis testing and FDR correction. **b.** Top ten differentially accessible regions associated with butyrate supplementation sorted by false-discovery rate adjusted *P* value (FDR < 0.05), and top GO ‘Biological Process’ terms associated with positive vs negative fold change determined by GREAT against a whole genome background using a binomial test over genomic regions, followed by multiple hypothesis testing with FDR corrected *P* values (FDR < 0.05). **c.** Normalized reads in binding sites at butyrate supplementation. Box plots display: The minimum, first quartile (Q1, 25^th^ percentile), median, third quartile (Q3, 75^th^ percentile), and maximum. The bottom of the box is Q1 and the top of the box is Q3. The line within the box represents the median (50^th^ percentile) value. The whiskers extend to the most extreme data points within 1.5 times the IQR (interquartile range). **d.** Correlation heatmap showing clustering of replicates from butyrate supplemented vs untreated group. **e.** Signal tracks representing differential accessibility between butyrate supplemented and untreated group. **f.** Top GO ‘Biological Process’ terms associated with H3K18bu in mouse intestines (5% AX). **g.** Top GO ‘Biological Process’ terms associated with H4K12bu in mouse intestines (5% AX). **h.** Annotated peak overlap between mouse intestinal H3K18bu (5% AX) targets and butyryl ATAC- seq in murine cells. **i.** Annotated peak overlap between mouse intestinal H4K12bu (5% AX) targets and butyryl ATAC- seq in murine cells.

## DISCUSSION

The SCFAs propionate and butyrate are produced by the microbiome and have broad biological effects. To gain insights into how they may directly affect gene regulation and expression we have combined histone PTM profiling by tandem LC-MS/MS with ChIP-seq, CUT&Tag, ATAC-seq and RNA-seq to understand the epigenetic regulatory function of these SCFA metabolites in CRC versus normal cells and *in vivo* (Extended Fig. 7). We identified the potential functional role of several propionyl- and butyryl-lysine modifications on H3 and H4 by identifying their genomic regions of interaction as well as their biological pathways. We have also shown the effect of these marks in promoting chromatin accessibility and changes in gene expression.

Our results link SCFA supplementation to epigenetic regulation of gene expression via non-canonical histone modifications and increased chromatin accessibility. In the context of CRC, SCFA supplementation led to a homeostatic dysregulation via hyperactivation of Wnt/β-catenin as well as TGF-β signaling pathways and activation of *MYC, FOS* and *JUN* oncogenes. Our results point to a 3-4-fold decrease in binding affinity of both Kbu marks to these oncogenic targets in normal versus cancer cells. To our knowledge, this is the first instance supporting Kbu direct targeting of genomic regions controlling growth and differentiation, rather than inhibition of deacetylation. It is also, to our knowledge, the first time that lncRNAs like *PRNCR1*, *PCAT1* and *CRAT37* were reported to be associated with Kbu binding in colorectal cells. It is generally thought that in cancer cells under aerobic metabolism butyrate (and to some extent propionate) accumulates and acts as an HDAC inhibitor leading to apoptosis^40^. Here we have expanded upon SCFA role as unique regulatory elements and have shown the genome-wide localization of H3K18pr/bu and H4K12pr/bu in CRC cells, and in case of Kbu in mouse intestines as well.

Our data point to a mechanism involving dysregulation of a broad range of oncogenes such as *MYC*, and tumor suppressing genes such as *TGF-βR2* and *SMAD2/3*. Our integrated multiomics data show the unique role of propionate and butyrate as regulators of histone acyl lysine levels leading to increased chromatin accessibility. In cancer cells, this results in overexpression of already high levels of proto-oncogenes controlling growth and differentiation, which may ultimately lead to cell death, especially in instances of elevated butyrate levels. This model extends the existing repertoire of SCFAs as epigenetic regulators beyond inhibitors of deacetylation. Regarding HDAC activity, especially in case of propionate, we have not observed significant increases in acetylation in our CRC cell line. In addition, for each Kpr and Kbu ChIP-seq experiment, we used the corresponding Kac marks as controls, to distinguish each histone mark’s unique targets under enrichment.

These data support a model that colonic SCFAs produced by microbial metabolism of fiber increase epithelial homeostatic gene expression pathways and impair carcinogenesis by direct histone modification. Given the rapid increase in colorectal cancers, especially in younger age populations recently, our results imply that dietary factors should be optimized to improve human health and diminish cancer onset. For example, the results raise the possibility of modulating histone post-translational modifications through dietary adjuvants, or through the creation of synthetic acyl chains to more precisely tune colonic epithelial gene expression^41^.

In addition, modulation of microbial populations or microbial metabolism may lead to improved epithelial homeostasis via epigenetic remodeling from microbial-derived acyl intermediates. Finally, given the high concentrations of SCFAs in the colonic environment, our results raise the possibility that chromatin-embedded acylations could serve as a storage depot of acylations that may be removed and further metabolized or recycled when colonic SCFA levels drop due to fiber limitation, or antibiotic usage that destroys microbial populations^42,43,44,45^. Taken together, our results highlight the crucial importance of understanding mechanisms of SCFA utilization and point towards ways of utilizing SCFA epigenetic modifications to improve human health.

## METHODS

### Cell lines

Adherent SW480, CCD841 and CT26 cells were obtained from ATCC (SW480 # CLL - 228, CCD841 CoN, # CRL - 1790, CT26.WT # CRL-2638) and cultured in commercially grown EMEM media containing 10 % fetal bovine serum (FBS)/1 % Pen/Strep (PS), or EMEM -glucose + 1 % PS (Sciencecell) and passaged every 3 - 5 days at 70 - 75 % confluence. CT26 cells were cultured in RPMI media containing 10 % FBS/1 % PS. Cells were treated with 0 - 10 mM NaPr and NaBu for 12 h. followed by cell counting and harvesting. All cell lines were maintained at 37° C (33° C in case of CCD841) in a humidified atmosphere containing 5 % CO_2_. All cell lines were tested for mycoplasma contamination by vendor and authenticated using STR profiling.

### Immunoblots

Histones were first acid-extracted as described below. Protein extracts were made in RIPA buffer and quantitated by BCA assay and diluted to equal concentrations and mixed with 4x LDS sample buffer and sample reducing agent (Invitrogen). Polyacrylamide gel electrophoresis was performed on NuPAGE Novex gradient gels (Thermo Fisher) followed by wet transfer to nitrocellulose membranes. Blocking was briefly performed with 5 % non-fat milk and primary antibody was incubated overnight at 4° C in 5 % milk, followed by washing in PBST, then with HRP-conjugated secondary antibody (Cell Signaling) at room temperature for 1 hour followed by washing, then developed with ECL pico or femto kits (Thermo Fisher) and imaged using ChemiDoc Imaging system (Bio-Rad).

### RNAi and cDNAs

HAT1 shRNAs were purchased from OriGene with the following sequences: A) GATGGCACTACTTTCTAGTATTTGAGAAG; B) AAGGATGGAGCTACGCTCTTTGCGACCGT; C) TCCTACAGTTCTTGATATTACAGCGGAAG.

The HAT1 cDNA was purchased from OriGene (catalog# RC209571L1) and the 5’ end was extended by a Gibson assembly reaction to add the additional 255 nucleotides to obtain a full-length clone. Mutagenesis of this cDNA was performed with the QuikChange II Site-Directed Mutagenesis kit with the following primers to make the D276Q acetylation-dead form: CAGTTCTTGATATTACAGCGCAAGATCCATCCAAAAGCTAT, ATAGCTTTTGGATGGATCTTGCGCTGTAATATCAAGAACTG.

To mutate the shRNA-C target site mutagenesis was performed with the following primers: GGTCGCAAAGAGCGTAGCTCCGTCTTTGTTGTATTTCTCAAATACTAGAAAGTA GTGCCATCTTTCATC, GATGAAAGATGGCACTACTTTCTAGTATTTGAGAAATACAACAAAGACGGAGC TACGCTCTTTGCGACC.

### Cell viability assays

Cell viability in the presence of increasing levels of NaPr and NaBu was performed using the CellTiter-Blue® Cell Viability Assay (Promega). TSA was used as a negative control. The assay measures the ability of living cells to convert a redox dye (resazurin) into a fluorescent end product (resorufin). Nonviable cells do not generate a fluorescent signal. 100 uL (or ∼ 5,000 adherent cells) were plated in 96-well plates in EMEM media containing 10 % fetal bovine serum (FBS)/1 % Pen/Strep (PS). All cell lines were maintained at 37° C in a humidified atmosphere containing 5 % CO_2_. The media was aspirated after 24 h and replaced with media containing increasing levels of NaPr and NaBu (0 - 100 mM) and TSA (0 - 100 µM) in ten-fold increments with three replicates per condition. After 72 h, 40 uL of CellTiter-Blue® reagent was added to each well and cells were incubated for 4 h. Following incubation, fluorescence at 560/590 nm was measured using Infinite M1000 Tecan i-control (v1.10.4.0) plate reader.

### Histone acid extraction

SW480 and CCD841 colorectal cells were grown in EMEM media containing 10 % fetal bovine serum (FBS)/1 % Pen/Strep (PS). All cell lines were maintained at 37° C. Prior to reaching confluence, cells were treated with increasing levels (0, 0.1, 1, 10 mM) of ^13^C3- Sodium Propionate (Cambridge Biosciences CLM-1865) for 12 h. Cells were harvested and pelleted at 218 x g (RCF) for 5 min at 4° C. Cells were resuspended in TEB buffer (PBS, supplemented with: 0.5 % Triton X-100, 2 mM PMSF, 0.02 % NaN_3_) in 1 mL per 10^7^ cells. Cells were incubated on ice for 10 min with gentle stirring then centrifuged at 1,966 x g (RCF) for 5 min at 4° C. Pellets were resuspended in 200 uL of extraction buffer per 10^7^ cells. Cells were then incubated on ice for 30 min, centrifuged at 3,209 x g (RCF) for 5 min 4° C. The supernatant was acetone precipitated in 600 uL of acetone per 10^7^ cells and incubated overnight at – 20° C. Histones were then acid extracted and acetone precipitated. Pellets were air dried and saved for downstream MS analysis or resuspended in 100 uL of H_2_O and diluted to 1 µg or 10 µg levels for immunoblot analysis. Protein concentration was determined using Pierce™ BCA Protein Assay Kits (Thermo Scientific cat # 23225) with BSA as a standard.

### Mass spectrometric identification of histone propionylation

Histone extracts were purified and derivatized according to Garcia et al.,^46^ and analyzed by nano-capillary liquid chromatography triple quadrupole mass spectrometry (nLC-QqQ MS). The digested and derivatized histone peptides were diluted in 0.1% TFA and injected to nLC-QqQ MS (Dionex nanoLC and a ThermoFisher Scientific TSQ Quantum). Peptides were first loaded to a trapping column (2 cm × 150 μm) and then separated with an analytical capillary column (10 cm × 75 μm). Both were packed with Magic C18 resin (Michrom). The chromatograph gradient was achieved by increasing percentage of buffer B from 2 – 35 % at a flow rate of 0.35 μL/min (A: 0.1 % formic acid in water, B: 0.1 % formic acid in acetonitrile) over 40 min. The peptides were then introduced into QqQ MS by electrospray from an emitter with 10 μm tip (New Objective) as they were eluted from capillary column. The QqQ settings were as follows: collision gas pressure of 1.5 mTorr; Q1 peak width of 0.7 (FWHM); cycle time of 3.5 s; skimmer offset of 10 V; electrospray voltage of 2.6 kV. Targeted histone PTM analysis was carried out using Multiple Reaction Monitoring (MRM), on a Thermo Scientific™ TSQ (Triple-Stage Quadrupole) instrument. Only specific precursor peptides were fragmented and specific product ion intensities were measured. Chromatographic separation and MS-measured intensities of different forms of peptides were used to distinguish modifications of interest. Exogenously introduced lysine propionylation was reported as ^13^C/^12^C heavy/light ratio.

### Determination of Acyl-CoA levels by LC-MS/MS and data analysis

Acetyl-CoA (≥ 93 % purity by HPLC), Propionyl-CoA (≥ 85 % purity by HPLC), and Butyryl-CoA (≥ 90 % purity by HPLC) standards were purchased from Sigma-Aldrich.^13^C_2_ Acetyl-CoA (97 % purity by HPLC) used as an internal standard (IS) was purchased from Millipore Sigma. Individual analytes and IS primary stock solutions (1 mg/mL) were prepared separately in 10 mM ammonium acetate/water. Intermediate stock solution was prepared by mixing individual stock solutions of each analyte followed by dilution to yield a 200 µg/mL working solution. This working solution was serially diluted with 10 mM ammonium acetate water to obtain a series of standard spiking solutions, which were used to generate the calibration curve. Calibration curves were prepared by spiking 10 µL of each standard working solution into 50 µL of homogenization buffer (methanol/water 1:1 v/v) followed by addition of 10 µL internal standard solution (1000 ng/mL ^13^C_2_ Acetyl-CoA). A calibration curve was prepared fresh with each set of samples. The calibration curve range for all analytes was 0.5 ng/mL – 1000 ng/mL using 50 µL aliquot. All cell lines were maintained at 37° C in EMEM media containing 10 % fetal bovine serum (FBS)/1 % Pen/Strep (PS) as described previously. The media was aspirated after 24 h and replaced with media containing increasing levels of NaPr and NaBu (0 - 100 mM) in ten-fold increments with three replicates per condition. Adherent cells were detached using 0.25 % (w/v) Trypin-0.53 mM EDTA solution placed at 37° C for 5 min. Cells were quenched in media and counted using the Automated Cell counter. The cells were resuspended in 2 mL then transferred to the 2 mL tubes you gave me, spun down again, aspirated and the pellets were stored in −80C. A 200 uL cold methanol/water (1:1 v/v) solution was added to each cell pellet. Samples were vortexed and sonicated for 2 minutes x 2 times; followed by centrifugation for 10 minutes at 2229 x g (RCF) at 4 °C. 50 µL supernatant was transferred to a clean eppendorf tube (1.7 mL). 10 µL internal standard solution was added to 50 µL supernatant aliquot followed by vortexing. 100 µL ice cold solution of methanol was added to the sample, vortexed, and then samples were placed at −20 °C for an hour to facilitate protein precipitation. After centrifugation, supernatant was transferred to a glass test tube and evaporated to dryness under nitrogen at 60 °C, reconstituted in 10 mM ammonium acetate buffer, sonicated, vortexed, transferred to an injection vial and analyzed by LC-MS/MS. All analyses were carried out by positive electrospray LC-MS/MS using a Waters Acquity I-class UPLC system with Waters Xevo TQ-XS triple quadrupole mass spectrometer (RRID:SCR_018510). Waters Atlantis T3 2.1×100 mm 3 um particle size column (Waters Corp., P/N1860203718) was operated at 40 °C at a flow rate 0.2 mL/min. Mobile phases consisted of A: 10 mM ammonium acetate in water and B: acetonitrile. Elution profile: linear gradient of 0 % - 25 % B from 0 to 5 minutes, followed by a linear gradient of 25 % - 95 % B in 1 minute, hold at 95 % for 2 minutes, equilibrate back to 0 % B; total run time was 10 minutes. Injection volume was 10 µL. Selected reaction monitoring (SRM) was used for quantification. Analytes mass transitions were as follows: Acetyl-CoA: m/z 810.0 → m/z 303.1 (quantifier) and m/z 810.0 → m/z 135.9 (qualifier); Propionyl-CoA: m/z 824.1 → m/z 317.1 (quantifier) and m/z 824.1 → m/z 135.9 (qualifier); Butyryl-CoA: m/z 838.1 → m/z 331.1 (quantifier) and m/z 838.1 → m/z 427.9 (qualifier); For internal standard: ^13^C_2_ Acetyl-CoA m/z 812.0→ m/z 305.0 (quantifier) and m/z 812.0 → m/z 135.9 (qualifier). Quantitative analysis was done with TargetLynx quantification software (Waters Corp.) using an internal standard approach. Calibration curves were linear (R > 0.99) over the concentration range using a weighting factor of 1/X^2^ where X is the concentration. The back-calculated standard concentrations were ± 15 % from nominal values, and ± 20 % at the lower limit of quantitation (LLOQ = 0.5 ng/mL).

### ChIP-seq and data analysis

For H3K18ac/pr/bu and H4K12ac/pr/bu ChIP-seq, cells were trypsinized and cross-linked with 1 % formaldehyde (EMD Millipore) for 10 min at RT. To quench the formaldehyde, 2 M glycine (Thermo Fisher Scientific) was added and incubated for 5 min at room temperature. Cells were washed with ice cold PBS twice, snap frozen and stored at − 80 °C. For ChIP-DNA preparation, cells were thawed by adding PBS and incubated at 4° C with rotation. Cells were treated with hypotonic buffer (20 mM HEPES pH 7.9, 10 mM KCl 1 mM EDTA pH 8.0, 10% glycerol) for 10 min on ice in the presence of protease inhibitors (G6521, Promega). Nuclear pellets were resuspended in RIPA buffer (Millipore) and incubated for 30 min on ice. Chromatin corresponding to 10 million cells for histone modifications was sheared with SFX250 Sonifier (Branson) 3 sets of 3 x 30 second sonications, set to intensity (output control) of 3.5. The lysates were transferred to Diagenode tubes, sonicate 16 rounds of 30 seconds on 30 seconds off, vortexing every 4^th^ round. Protein G beads (80 uL) and lysates were washed with RIPA buffer and immunoprecipitated with antibodies targeting H3K18ac (Abcam #ab40888), H3K18pr (PTM Bio #PTM-213), H3K18bu (Abcam #ab241458), H4K12ac (Abcam #ab46983), H4K12pr (PTM Bio #PTM-209), H4K12bu (# ab241120), 5 μg for each condition, at 4° C overnight on a nutator. For the input sample, 200 μl of sheared nuclear lysate was removed and stored overnight at 4° C. On the second day, supernatants containing ChIP-DNA and input was reverse crosslinked by incubating overnight at 65° C in 1% SDS, 1X TE buffer. On the third day, ChIP-DNA was treated with 250 μL 1X TE containing 100 μg RNase A (Qiagen) and 5.0 μL of 20 mg/mL Proteinase K (ThermoFisher Scientific) and then purified using Qiagen QIAquick purification columns. The ChIP-DNA samples were end-repaired using End-It DNA End Repair Kit (Lucigen) and A-tailed using Klenow Fragment and dATP (New England Biolabs). Illumina TruSeq adapters (Illumina) were ligated and libraries size-selected (200 − 400 bp) by gel extraction before PCR amplification. The purified libraries were subjected to paired-end sequencing on the Illumina HiSeq 4000/Novaseq 6000 SP to obtain an average of approximately 30 − 35 million uniquely mapped reads for each sample (Stanford Center for Genomics and Personalized Medicine, supported by NIH grants S10OD025212 and 1S10OD021763).

The resulting data were processed using the Kundaje Lab ChIP-seq ENCODE processing pipeline v2.2.2 (https://github.com/ENCODE-DCC/chip-seq-pipeline2). Briefly, this pipeline takes FASTQ files for ChIP samples, and input as controls and outputs peak calls (bound regions; BR). Alignments to BR (GRCh38.p13) were performed using Bowtie2 (https://doi.org/10.1186/gb-2009-10-3-r25) and peak calling was performed using MACS2 (https://doi.org/10.1186/gb-2008-9-9-r137). Peak flies were analyzed for differential binding using DiffBind R package v.2.4.8 to produce a count matrix with an FDR-adjusted *P* value cutoff of < 0.05. Data visualization was performed in R as well as using the Integrative Genomics Viewer (http://www.broadinstitute.org/igv/). Differential motif enrichment analysis in HOMER (http://homer.ucsd.edu/homer/) was performed using function findMotifsGenome with default parameters to search for motif enrichment in the full accessible regions, and control IP and/or input used as background. Annotation was performed using ChIPpeakAnno and ChIPseeker R packages. TSS distribution heatmaps and profiles were determined using computeMatrix, plotHeatmap and plotProfile functions within deepTools v.3.1.0 (https://deeptools.readthedocs.io/en/develop/index.html). Genomic coordinate overlaps were determined using the intersect ‘function’ in bedtools (v.2.31.0), whereby the original entry in one set was reported once if any overlaps were found in the second set, effectively reporting that at least one overlap was found (https://bedtools.readthedocs.io/en/latest/content/tools/intersect.html). Gene Ontology analysis was performed using GREAT (http://great.stanford.edu/public/html/index.php) and ShinyGO (DOI: 10.1093/bioinformatics/btz931). Images in KEGG pathway analysis were produced by PATHVIEW(https://bioconductor.org/packages/release/bioc/html/pathview.html).

### ATAC-seq and data analysis

ATAC-seq was performed using Omni-ATAC as described in Buenrostro et al.,^47^. Briefly, 50,000 viable cells were pelleted at 500 RCF for 5 min at 4° C. Cells were resuspended in 50 μl of ATAC-Resuspension Buffer (RSB) containing 0.1 % NP40, 0.1 % Tween-20, and 0.01 % Digitonin and mixed by pipetting. Following incubation on ice for 3 min, cells were washed with 1 mL cold RSB containing 0.1 % Tween-20 but no NP40 or Digitonin. Nuclei were pelleted at 500 RCF for 10 min at 4° C. Cells were resuspended in 50 μL of transposition mix containing 25 µL 2X TD buffer, 2.5 µL transposase (Illumina Tagmented DNA enzyme and buffer kit, Ref 20034210) (100 nM final), 16.5 µL PBS, 0.5 µL of 1 % Digitonin, 0.5 µL 1 % Digitonin, 0.5 µL of 10 % Tween-20, and 5 µL H_2_O). The final reaction was incubated at 37° C for 30 min in a thermomixer with 1000 RCF mixing. Pre-amplified transposed fragments were cleaned using the Zymo DNA Clean and Concentrator-5 Kit (cat # D4013). DNA was eluted in 21 µL of elution buffer. Samples then underwent 5 cycles of PCR using NEBNext 2x MasterMix. Each reaction contained 2.5 µL of 25 µM i5 primer, 2.5 µL of 25 µM i7 primer, 25 uL of 2x NEBNext master mix, and 20 µL of transposed/cleaned-up sample. Reactions were PCR amplified (72° C for 5 min, 98° C for 30 sec, followed by 5 cycles of (98° C for 10 min, 63° C for 30 sec, 72° C for 1 min) then held at 4° C. Using 5 µL of pre-amplified mixture, 15 μL qPCR (Applied Biosystems, Quantstudio 6 Flex) was then performed to determine the additional number of cycles. The conditions for qPCR were: 3.76 μL sterile H_2_O, 0.5 μL 25 µM i5 primer, 25 µM i7 primer, 0.24 μL 25x SYBR gold (in DMSO), 5 μL 2x NEBNext master mix and 5 μL of pre-amplified sample. Cycling conditions for qPCR were 98° C for 30 sec, followed by 20 cycles of (98° C for 10 sec, 63° C for 30 sec, 72° C for 1 min). qPCR amplification profiles were then manually assessed to determine the required number of additional cycles. Using the remainder of the pre-amplified DNA, 2 - 3 additional cycles were performed. Final PCR reactions were purified using Zymo DNA Clean and Concentrator-5 Kit (cat # D4013) and eluted in 20 μL of H_2_O. Amplified DNA library concentration was determined using Qubit 4 Fluorometer (Invitrogen). Library quality and size was assessed on an Agilent Bioanalyzer 2100 system using a high-sensitivity DNA kit. Multiplexed libraries were paired-end sequenced on the Illumina HiSeq 4000/ Novaseq 6000 SP to obtain an average of approximately 50 million uniquely mapped reads per sample (Stanford Center for Genomics and Personalized Medicine, supported by NIH grants S10OD025212 and 1S10OD021763). The resulting data were processed using the Kundaje Lab ENCODE ATAC-seq processing pipeline v2.2.3 (https://github.com/ENCODE-DCC/atac-seq-pipeline). Briefly, this pipeline takes FASTQ files as input and outputs peak calls (accessible regions, AR). All annotations have been carried out against GENCODE annotation database for human (GRCh38.p13) and mouse (GRCm38.p4) genomes. Alignments to AR were performed using Bowtie2 (https://doi.org/10.1186/gb-2009-10-3-r25) and peak calling was performed using MACS2 (https://doi.org/10.1186/gb-2008-9-9-r137). Peak flies were analyzed for differential accessibility using DiffBind v.2.4.8 R package to produce a counts matrix with an FDR-adjusted *P* value cutoff of < 0.05. Venn diagrams generated by https://bioinformatics.psb.ugent.be/webtools/Venn/.

### CUT&Tag and data analysis

CUT&Tag analysis of NaBu treated CCD841 cells and mouse intestinal samples were performed using CUT&Tag-IT^TM^ assay kits (Active Motif # 53160, 53170). Approximately 30 mg of tissue was homogenized per reaction by first chopping frozen tissue into smaller pieces in a petri dish with a razor blade on dry ice. 1 mL of CUT&Tag-IT Lysis Buffer (50 mM Tris pH 8.0, 10 mM EDTA, 0.4 % w/v SDS, 0.1 % protease inhibitor cocktail) was added per 10 mg of tissue and samples were further cut and minced in bulk. Samples were then Dounced 20 times with a loose pestle and 10 times with a tight pestle. Samples were then filtered with a 40 µM strainer in 15 mL conical tubes. Samples were centrifuged at 500 x g for 5 min at 4 °C. Samples of nuclei suspension were counted and normalized to ∼ 500,000 nuclei per reaction. CCD841 cells (∼ 500,000 / reaction) treated with NaBu for 12 h were harvested by centrifugation at 600 x g at RT. Both CCD841 cells and mouse tissue extracts were washed with 1X Wash Buffer (1 M HEPES pH 7.5, 1.5 mL 5 M NaCl, 12.5 μL 2 M spermidine), containing protease inhibitor cocktail (10 μL per 1 mL of Wash Buffer), and resuspended in Concanavalin A Beads slurry in a 1X Binding Buffer (1 M HEPES pH 7.5, 100 μL 1 M KCl, 10 μL 1 M CaCl_2_ and 10 μL 1M MnCl_2_). Cells and beads were incubated on an end-over-end rotator for 10 min. Cells were resuspended in ice-cold Antibody Buffer containing 2 mL Dig-Wash buffer (5 % digitonin with 40 mL 1X Wash buffer with PIC) mixed with 8 μL 0.5 M EDTA and 6.7 µL 30 % BSA not to exceed 500,000 cells per 50 μL reaction volume. 5 μg of undiluted primary antibody was added to each sample for an overnight incubation at 4° C with orbital mixing. 100 μL of Guinea Pig Anti-Rabbit secondary antibody diluted 1:100 in Dig-Wash buffer was then added to each reaction and the samples were incubated on an orbital rotator for 60 min at RT. 100 μL of 1:100 diluted CUT&Tag-IT^TM^ Assembled pA-Tn5 Transposomes in Dig-300 Buffer (1 M HEPES pH 7.5, 3 mL 5 M NaCl and 12.5 μL 2 M spermidine with 5 % digitonin and 0.01 % protease inhibitor cocktail) was added to each sample and reactions were incubated on an orbital rotator for 60 min at RT. Following three washes with Dig-300 buffer, 125 μL of Tagmentation Buffer (5 mL Dig-300 buffer and 50 µL 1 M MgCl2) was added to each sample and reactions were incubated at 37° C for 1 h. To stop the tagmentation and solubilize DNA fragments, each sample received 4.2 μL of 0.5 M EDTA, 1.25 μL of 10 % SDS, 1.1 μL of Proteinase K (10 mg/mL). Samples were mixed and incubated at 55° C for 60 min. Following washing, samples were centrifuged at 17,000 x g for 2 min. PCR amplification was performed by adding 30 μL of Tagmented DNA from each reaction to 1 uL of dNTPs (10 mM), 0.5 μL of NEBNext Q5 High-Fidelity DNA Polymerase, 10 μL 5X Q5 Reaction Buffer, 3.5 μL of nuclease-free H2O and 2.5 μL of unique combinations of Nextera i5 and i7 indexing primers to a total of 50 μL reaction. PCR was performed using the following program on a thermal cycler with a heated lid: Cycle 1: 72° C for 5 min (gap filling) Cycle 2: 98° C for 30 sec Cycle 3: 98° C for 10 sec Cycle 4: 63° C for 10 sec. Repeated Cycles 3 - 4 14 times. Held at 72° C for 1 min and held at 10° C. PCR reactions were purified using Zymo DNA Clean and Concentrator-5 Kit (cat # D4013) and eluted in 20 μL of nuclease-free H_2_O. Amplified DNA library concentration was determined using Qubit 4 Fluorometer (Invitrogen). Library quality and size was assessed on an Agilent Bioanalyzer 2100 system using a high-sensitivity DNA kit. Multiplexed libraries were paired-end sequenced on the Illumina HiSeq 4000 / Novaseq 6000 SP to obtain an average of approximately 50 million uniquely mapped reads per sample (Stanford Center for Genomics and Personalized Medicine, supported by NIH grants S10OD025212 and 1S10OD021763). The resulting data were processed using the Kundaje Lab ENCODE ATAC-seq processing pipeline v2.2.3 (https://github.com/ENCODE-DCC/atac-seq-pipeline). Briefly, this pipeline takes FASTQ files as input and outputs peak calls (accessible regions, AR). All annotations have been carried out against GENCODE annotation database for human (GRCh38.p13) and mouse (GRCm38.p4) genomes. Alignments to AR were performed using Bowtie2 (https://doi.org/10.1186/gb-2009-10-3-r25) and peak calling was performed using MACS2 (https://doi.org/10.1186/gb-2008-9-9-r137). Peak flies were analyzed for differential accessibility using DiffBind v.2.4.8 R package to produce a counts matrix with an FDR-adjusted *P* value cutoff of < 0.05.

### RNA-seq and data analysis

RNA was extracted using the RNeasy Mini Kit (#74136, Qiagen). Libraries were prepared using NEBNext Ultra II RNA library prep kit for Illumina (# E7770) and rRNA depletion kit (# E6310). Paired-end sequencing on the Illumina Novaseq 6000 SP yielded an average of approximately 20 million uniquely mapped reads per sample for mRNA-seq (Stanford Center for Genomics and Personalized Medicine, supported by NIH grants S10OD025212 and 1S10OD021763). The resulting data were aligned to the human genome (GRCh38.p13) by STAR v.2.5.4b (https://github.com/alexdobin/STAR). The aligned transcripts were quantitated based on features in the GENCODE annotation database (GRCh38.p13) by RSEM v.1.3.1 (http://deweylab.biostat.wisc.edu/rsem/). Differentially expressed genes were detected using DESeq2 v.1.20.0 R package with a FDR-adjusted *P* value cutoff of < 0.05 (https://bioconductor.org/packages/release/bioc/html/DESeq2.html). Venn diagrams generated by https://bioinformatics.psb.ugent.be/webtools/Venn/.

### Animal handling and preparation of intestinal tissue samples

All experiments were approved by and conducted in strict accordance with Stanford University’s Administrative Panel on Laboratory Animal Care (APLAC 34017). Male CETP-ApoB-100 transgenic mice were procured from Taconic Biosciences (New York, USA). All animals were housed in a temperature-controlled, specific pathogen-free environment, with a 12:12 light-dark cycle, temperature 24 ± 1° C and humidity ranging between 40 – 60 %. Food and water were available *ad libitum*. Mice were assigned randomly to two dietary groups: 1) low fat control diet (10% fat, TD. 08485, Envigo Teklad, Madison, WI, USA), or a 2) high fat, high sucrose diet (42% fat, TD 88137, Envigo Teklad, Madison, WI, USA). The high fat, high sucrose (HFS) diet was selected to mimic the *western diet* aiming to characterize and augment atherosclerosis development in the ApoE-deficient transgenic mouse. Over a span of 4 weeks, both groups were subjected to their respective diets. Following the initial 4-week duration, the control group continued on their diet for an additional 4 weeks. In contrast, the HFS group received a supplementation of 5 % arabinoxylan (5 % w/w of total HFS, blended in-house with their powdered HFS diet) for an additional 4 weeks. At the conclusion of this second 4-week period, all mice underwent a 6-hour fast and were then anesthetized with isoflurane and subsequently sacrificed via cervical dislocation. The large intestine was dissected, weighed, flash frozen in liquid nitrogen and subsequently stored in – 80° C until analysis.

There was no randomization in the organization of the experimental conditions or stimulus presentation. Data collection and analysis were not performed blind to the conditions of the experiments. No animals or data points were excluded from the analyses. No statistical methods were used to pre-determine sample sizes, but our sample sizes are similar to those reported in a previous publication^48^. The data met the assumptions of the statistical tests used, including normality and equal variances were formally tested by running diagnostics and visualizations such as residual, homoscedasticity and QQ plots using Brown-Forsythe and Shapiro-Wilk tests.

## Supporting information

Nshanian_et_al_Supplemental_Figures

Nshanian_et_al_Extended_Figures

## DATA AVAILABILITY

ChIP-seq, ATAC-seq and CUT&Tag raw data and differential peak call files have been uploaded to GEO with accession numbers GSE252649, GSE252652 and GSE252754, respectively. RNA-seq data are deposited in GEO with accession number GSE252753. GSE252653 is the reference for the Series of four data sets.

## ACKNOWLEDGEMENTS

We wish to thank members of Snyder laboratory for useful discussions and suggestions about this project. We also thank Stanford Center for Genomics and Personalized Medicine Sequencing Center (supported by NIH grants S10OD025212 and 1S10OD021763), Jaison Arivalagan, Young Ah Goo and Juliette Andria Morris of the Northwestern Proteomics Center of Excellence Core Facility. J.J.G. was supported by CPRIT grant RR200090 and NIH/NCI grant K08CA245024. M.N. was supported by NIH grant 3UM1HG009442-04S1 to M.P.S. for the Production Center of Mapping Regulatory Regions of the Human Genome. This work utilized the Xevo TQ-XS mass spectrometer system (RRID:SCR_018510) that was purchased with funding from National Institutes of Health Shared Instrumentation Grant S10OD026962.

## AUTHOR CONTRIBUTIONS STATEMENT

Conceptualization, M.N., J.J.G., and M.P.S.; Investigation, M.N., J.J.G., B.S.G., F.C., S.L., J.M.C., L.A.; Methodology, M.N., J.J.G., B.S.G., and M.P.S.; Validation, M.N., B.S.G., S.M.W., and M.P.S.; Formal analysis, M.N.; Data curation, M.N.; Visualization, M.N.; Writing - original draft, M.N.; Writing - review & editing, M.N., J.J.G., S.L., S.M.W., and M.P.S.; Resources, M.P.S., J.J.G., N.L.K., Y.Z.; Funding acquisition, M.P.S.

---

**Supplementary Fig. 1.**
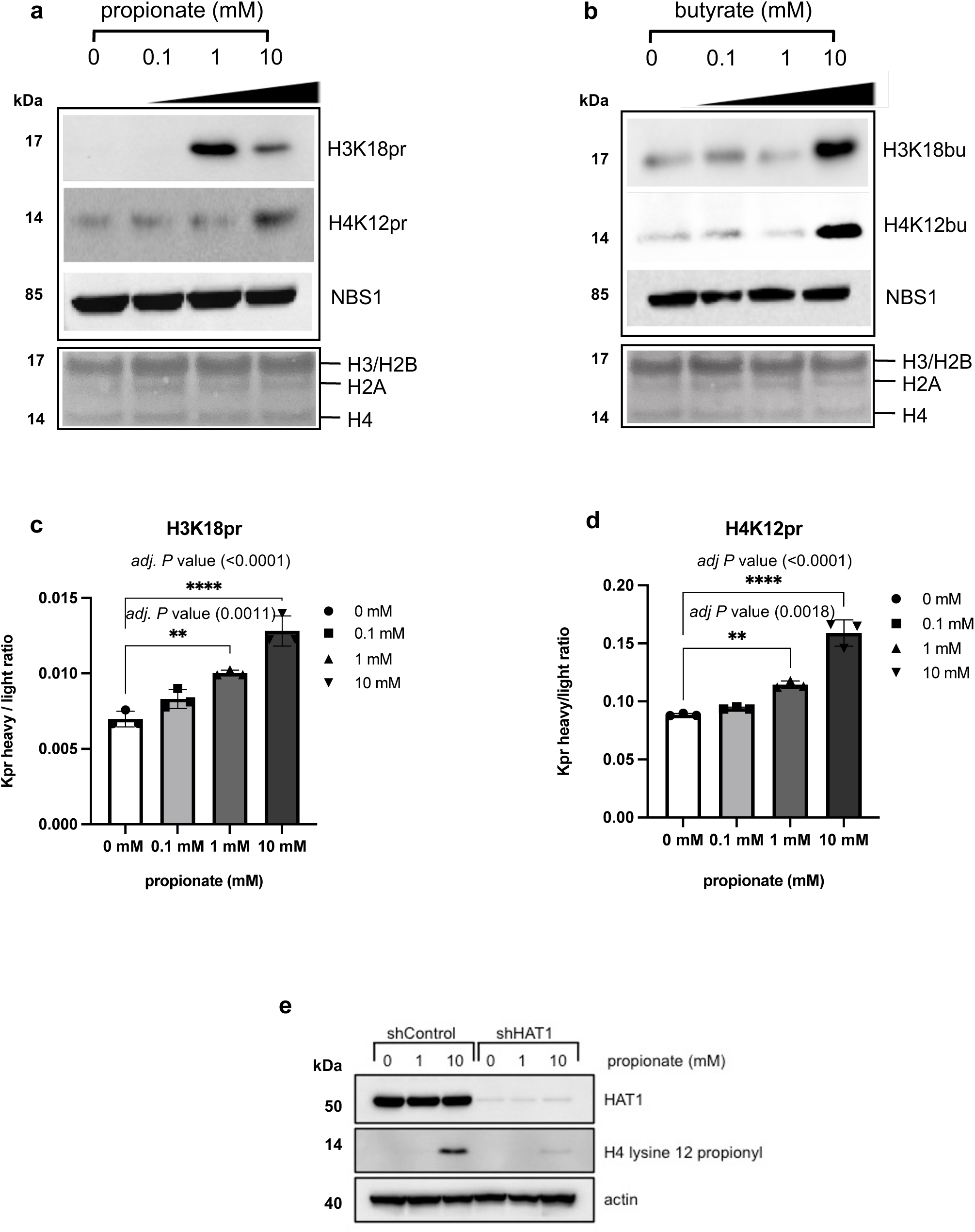
Immunoblots of acid-extracted **a** H3K18/H4K12pr and **b** H3K18/H4K12bu histone marks at 0 ⎼ 10 mM sodium propionate/butyrate treatments with anti-NBS1 as control (top). SDS-PAGE of histones H2A/B, H3/H4 (bottom). Dose-dependent ^13^C-propionate incorporation into H3 and H4 as measured by increases in the heavy/light propionyl lysine containing peptides. Heavy/light propionyl lysine containing peptides representing **c** H3K18pr and **d** H4K12pr levels at 0, 0.1, 1, and 10 mM ^13^C-propionate supplementation (mean ± SD, n = 3). Multiple comparisons by ordinary, one-way ANOVA followed by hypothesis testing using the Bonferroni correction method with 0.05 *P* value cutoff. \*\**P* < 0.01, **** *P* < 0.0001 **e** Depletion of HAT1 diminishes incorporation of propionate into the H4 lysine 12 site.

**Supplementary Fig. 2.**
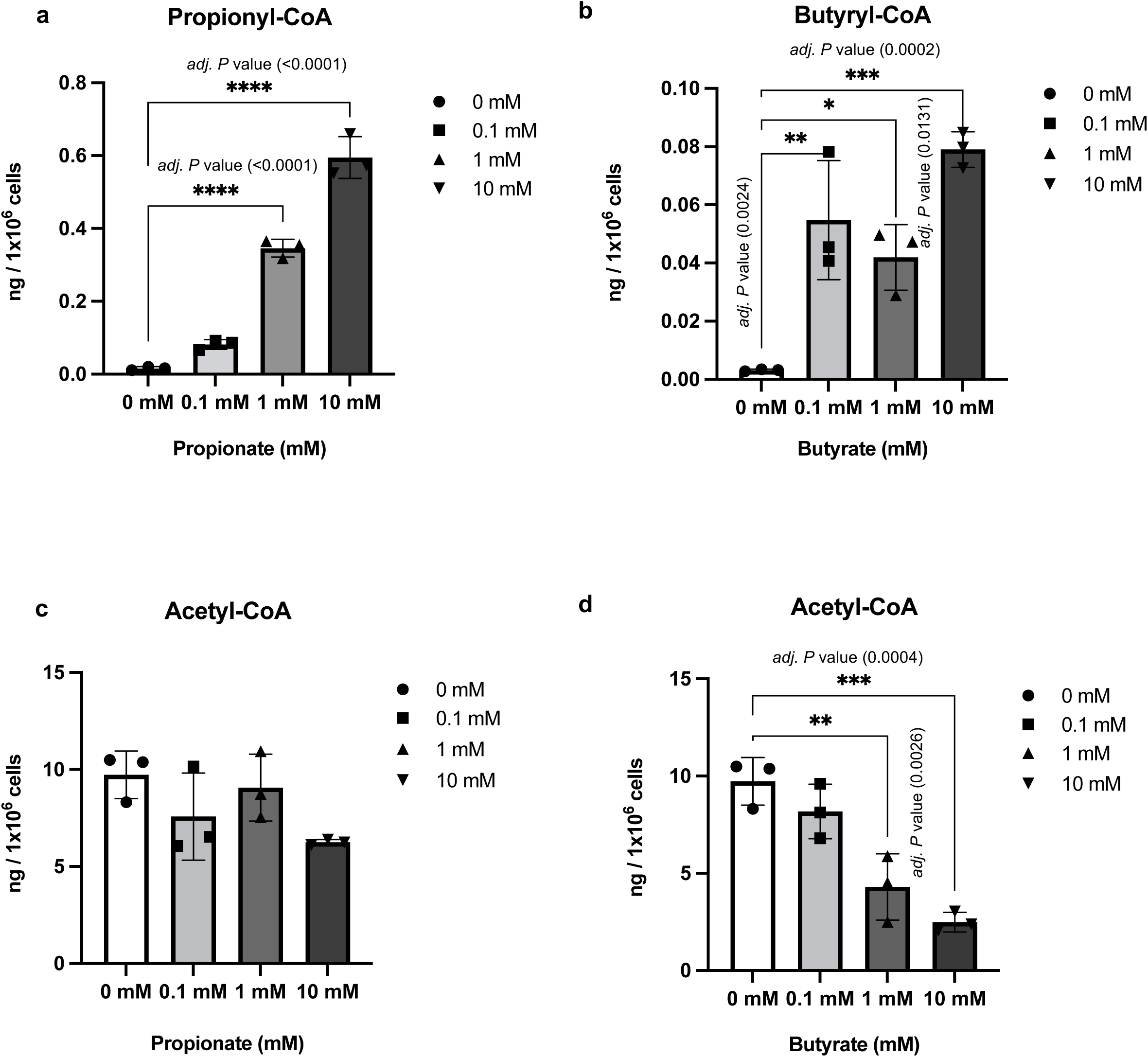
Quantitative analysis of acyl-CoA levels by LC-MS/MS following SCFA supplementation. **a** Dose-dependent increases in **a** Propionyl-CoA and **b** Butyryl-CoA levels upon increasing propionate and butyrate supplementation at 0, 0.1, 1, and 10 mM. Acetyl-CoA levels upon increasing **c** propionate and **d** butyrate supplementation. Quantitative analysis was done by multiple reaction monitoring (MRM) using an internal standard approach. Calculated Acyl-Co concentrations in each sample were normalized to the number of cells (mean ± SD, n = 3). Multiple comparisons by ordinary, one-way ANOVA using hypothesis testing followed by Bonferroni correction method with 0.05 *P* value cutoff. \**P* < 0.05, \*\**P* < 0.01, \*\*\**P* < 0.001, \*\*\*\**P* < 0.0001.

**Supplementary Fig. 3.**
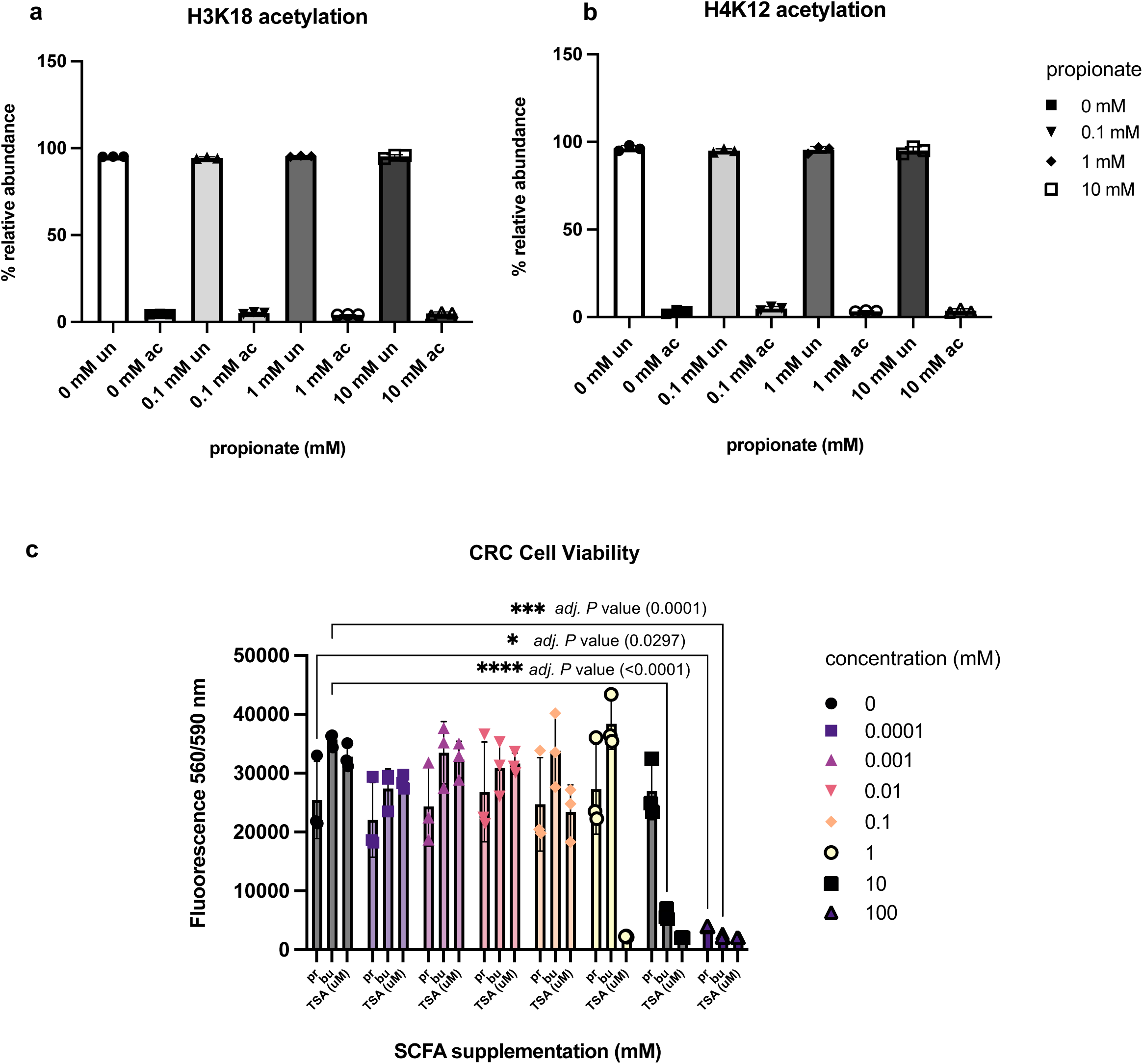
H3 and H4 acetylated vs unmodified states as a function of propionate supplementation. Relative abundances of acetylated vs unmodified states on **a** H3K18 **b** H4K12 **c** H3K9 and **d** H3K23 following 0, 0.1, 1, and 10 mM propionate supplementation (mean ± SD, n = 3). **e** CRC cell viability as a function of NaPr and NaBu supplementation over 72 hrs. as measured by CellTiter-Blue® fluorescence assay. TSA, a known HDAC inhibitor with IC50 ∼ 2 nM was used as a negative control on a μM level. (mean ± SD, n = 3). Multiple comparisons by two-way ANOVA using statistical hypothesis testing followed by Bonferroni correction method with 0.05 *P* value cutoff. \**P* < 0.05, ***p < 0.001, ****p < 0.0001.

**Supplementary Fig. 4.**
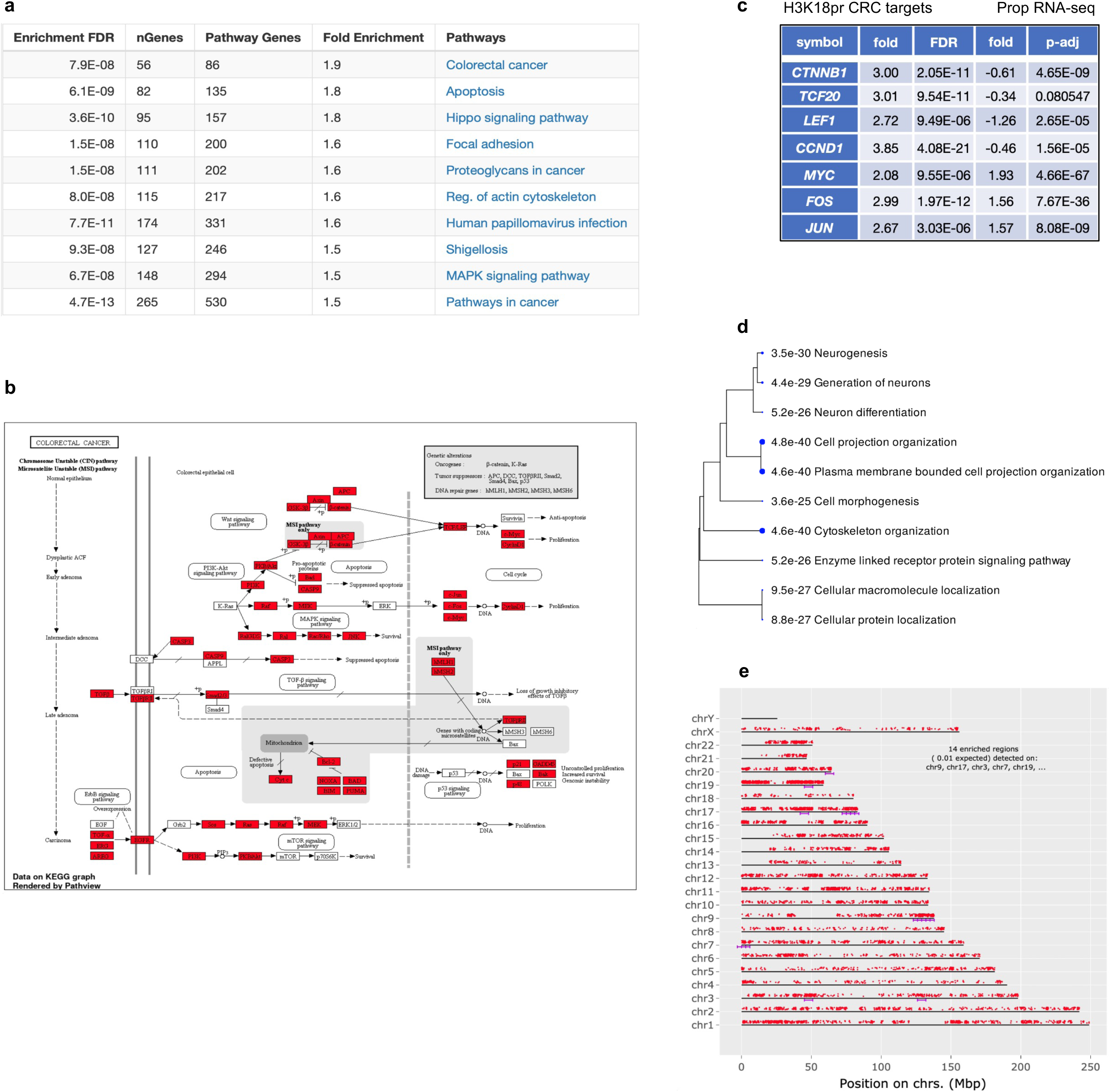
KEGG pathway analysis of H3K18pr-associated genes. **a** Top ten pathways with their number of genes and log2 fold enrichment. FDR is calculated from a nominal *P* value obtained from a hypergeometric test (FDR < 0.05). Fold enrichment is calculated as percentage of H3K18pr differentially bound genes associated with a pathway divided by the corresponding percentage in input. **b** CRC KEGG pathway enrichment by Pathview. Genes that are overrepresented compared to input are in red. **c** H3K18pr-associated differential binding of key CRC genes and log2 fold changes in their expression levels, as determined by differential RNA-seq of 10 mM treated vs. untreated conditions. n = 3 experimental replicates per condition (FDR < 0.05). **d** Hierarchical clustering tree summary of correlations among significant pathways in H3K18pr-associated annotated genes. Hierarchical clustering of the pathways was performed using ShinyGO. Pathways were clustered together based on shared genes and gene enrichment analysis was performed using two-sided Fisher’s exact test, and FDR correction was applied to adjust for multiple comparisons in the pathway analysis and hierarchical clustering. Size of dots indicates statistically significant FDR adjusted (FDR < 0.05) *P* values. **e** Chromosomal position of H3K18pr-associated regions represented by red dots. Purple lines represent statistically significant enrichment compared to input. The genome was scanned with a sliding window (size 6 Mb) further subdivided into 2 equal-sized steps for sliding. Within each window a hypergeometric test was used to test for enrichment over input. FDR-adjusted *P* value cutoff for window was 1E-05. Chromosomes may be partly shown due to scaling to last genes location. Gene chromosomal mapping was performed using ShinyGO.

**Supplementary Fig. 5.**
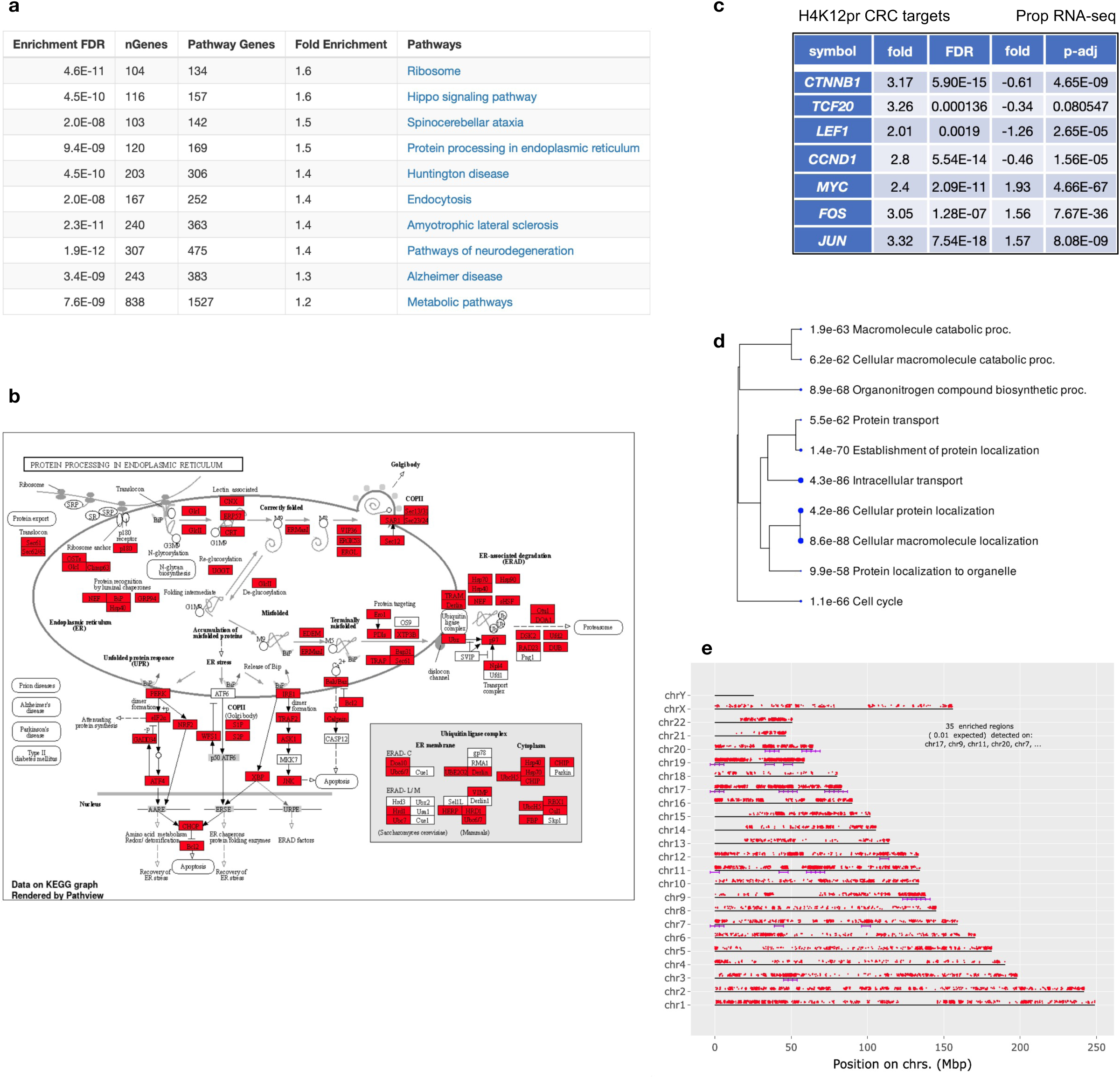
KEGG pathway analysis of H4K12pr associated genes. **a** Top ten pathways with their number of genes and log2 fold enrichment. FDR is calculated from a nominal *P* value obtained from a hypergeometric test (FDR < 0.05). Fold enrichment is calculated as percentage of H4K12pr differentially-bound genes associated with a pathway divided by the corresponding percentage in input. **b** Protein Processing in ER KEGG pathway enrichment by Pathview. Genes that are overrepresented compared to input are in red. **c** H4K12pr-associated differential binding of key CRC genes and log2 fold changes in their expression levels, as determined by differential RNA-seq of 10 mM treated vs. untreated conditions. n = 3 experimental replicates per condition (FDR < 0.05). **d** Hierarchical clustering tree summary of correlations among significant pathways in H3K18pr-associated annotated genes. Hierarchical clustering of the pathways was performed using ShinyGO. Pathways were clustered together based on shared genes and gene enrichment analysis was performed using two-sided Fisher’s exact test, and FDR correction was applied to adjust for multiple comparisons in the pathway analysis and hierarchical clustering. Size of dots indicates statistically significant FDR adjusted (FDR < 0.05) *P* values. **e** Chromosomal position of H3K18pr-associated regions represented by red dots. Purple lines represent statistically significant enrichment compared to input. The genome was scanned with a sliding window (size 6 Mb) further subdivided into 2 equal-sized steps for sliding. Within each window a hypergeometric test was used to test for enrichment over input. FDR-adjusted *P* value cutoff for window was 1E-05. Chromosomes may be partly shown due to scaling to last genes location. Gene chromosomal mapping was performed using ShinyGO.

**Supplementary Fig. 6.**
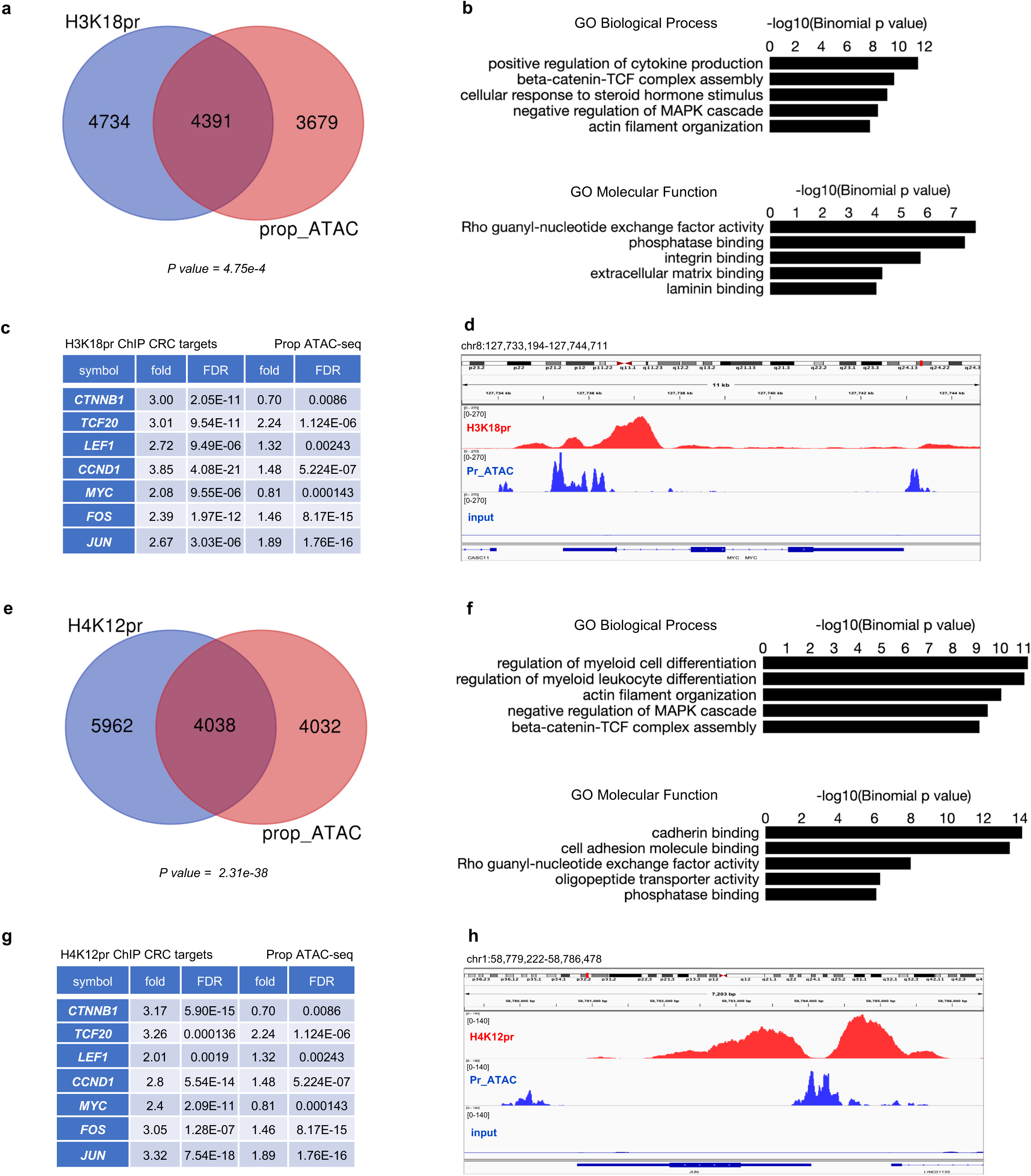
H3K18/H4K12pr ChIP-seq and propionyl ATAC-seq integration. **a, e** Overlap between TSS-proximal regions (+/- 1 Kb) for H3K18pr and H4K12pr differentially bound genes by ChIP-seq and differentially accessible genes following 10 mM propionate supplementation. Significance of overlap determined by hypergeometric test-generated *P* value. **b, f** GO ‘Biological Process’ and ‘Molecular Function’ pathway terms associated with H3K18pr and H4K12pr differentially bound genomic coordinates that are also present in ATAC-seq sorted by binomial *P* value. **c, g** Log2 fold change and FDR-adjusted *P* value in CRC-relevant gene targets associated with H3K18/H4K12pr ChIP-seq and propionyl ATAC-seq. **d, h** Signal tracks for *MYC* and *JUN* regions showing ChIP-seq and ATAC-seq profiles with input as background.

**Supplementary Fig. 7.**
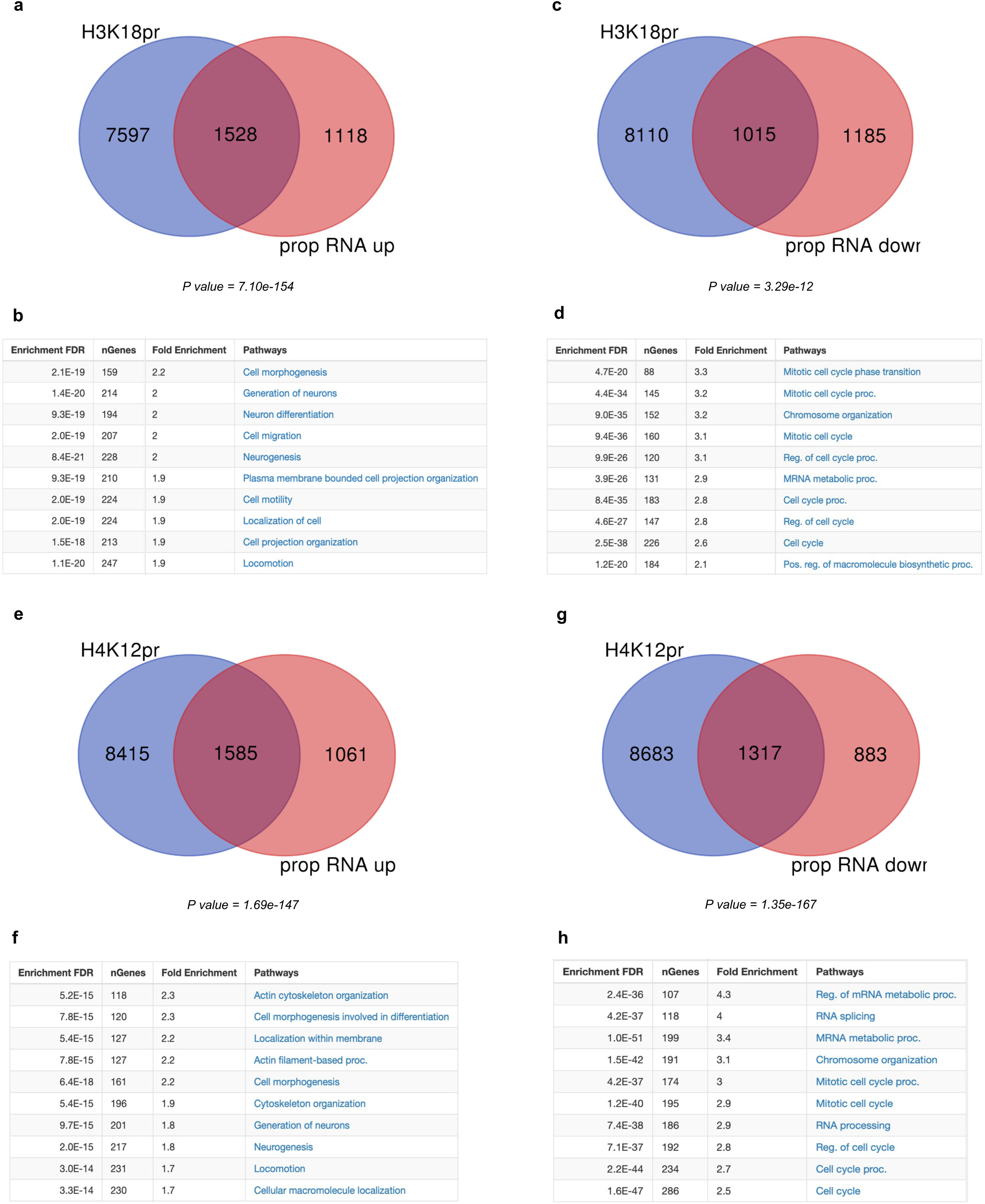
H3K18/H4K12pr ChIP-seq and propionyl RNA-seq integration. Overlap between TSS-proximal regions (+/- 1 Kb) for H3K18pr and H4K12pr associated genes by ChIP-seq, and upregulation of gene expression in 10 mM NaPr treated group **a**, **e** vs downregulation in the untreated group **c**, **g** by RNA-seq. Significance of overlap determined by hypergeometric test-generated *P* value. GO ‘Biological Process’ pathway terms associated with overlapping Kpr targets and upregulated genes **b**, **f** and downregulated genes **d**, **h** sorted by log2 fold enrichment.

**Supplementary Fig. 8.**
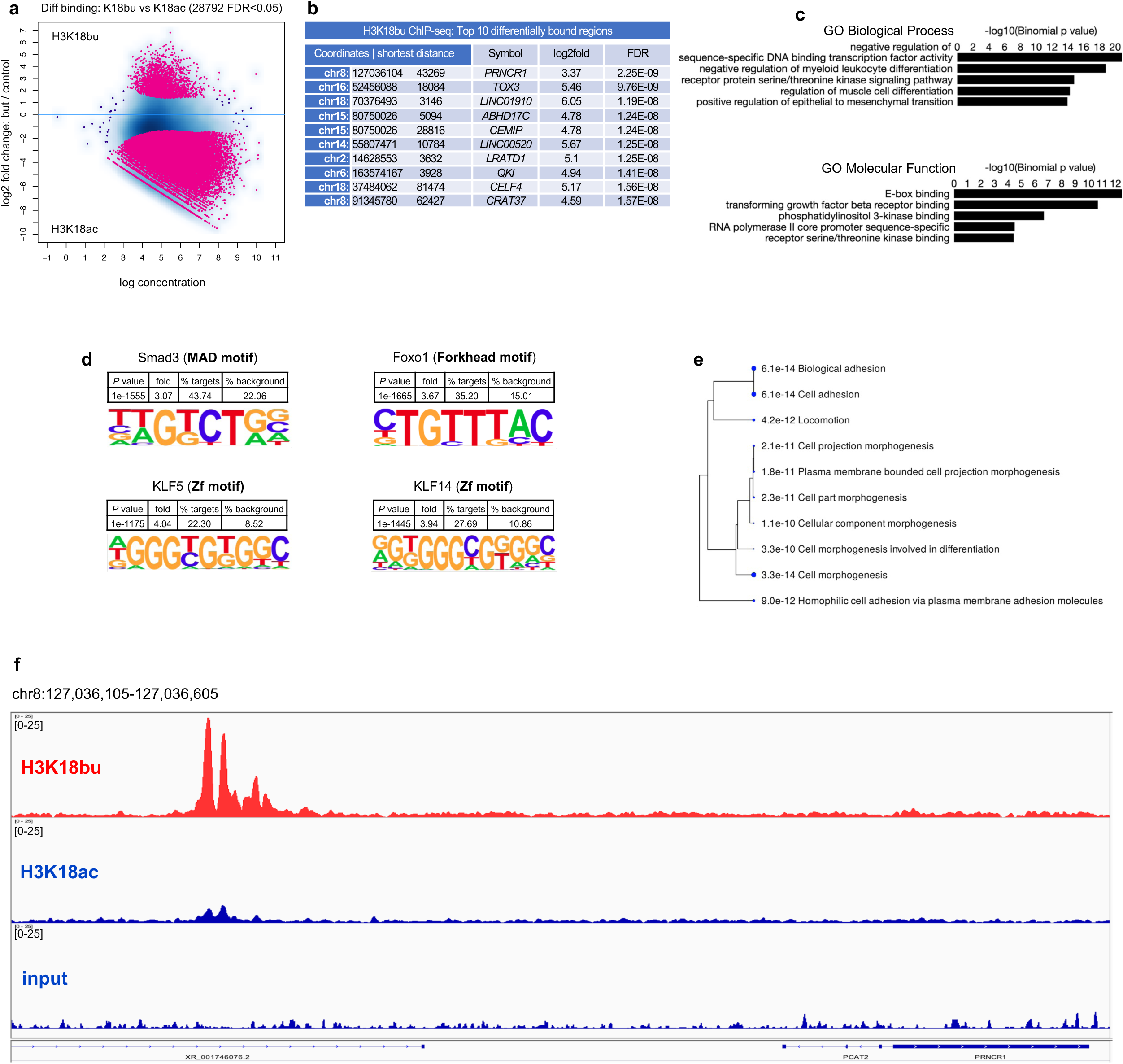
Genome-wide H3K18bu distribution. **a** H3K18bu vs H3K18ac differential binding following 1 mM butyrate supplementation. Sites identified as significantly differentially bound are shown in red. Differential binding was performed by DiffBind package with DESeq2 using a two-sided tests for both increased and decreased binding affinity between conditions followed by by multiple hypothesis testing and FDR correction. **b** Top ten differentially bound regions associated with H3K18bu sorted by false-discovery rate adjusted *P* value (FDR < 0.05). **c** Top GO ‘Biological Process’ and ‘Molecular Function’ terms associated with H3K18bu-bound *cis*-regulatory elements determined by GREAT against a whole genome background using a binomial test over genomic regions, followed by multiple hypothesis testing using FDR corrected *P* values (FDR < 0.05). **d** Differential motif analysis of H3K18bu vs H3K18ac peaks was analyzed by HOMER, using a one-sided hypergeometric test for overrepresentation (enrichment) of motifs in the target sequences compared to the background, followed by multiple hypothesis testing and FDR correction. **e** Hierarchical clustering tree summary of correlations among significant pathways in H3K18pr-associated annotated genes. Hierarchical clustering of the pathways was performed using ShinyGO. Pathways were clustered together based on shared genes and gene enrichment analysis was performed using two-sided Fisher’s exact test, and FDR correction was applied to adjust for multiple comparisons in the pathway analysis and hierarchical clustering. Size of dots indicates statistically significant FDR adjusted (FDR < 0.05) *P* values. **f** Signal tracks of 71 Kb-spanning *PRNCR1* region showing H3K18bu vs H3K18ac binding with input as background.

**Supplementary Fig. 9.**
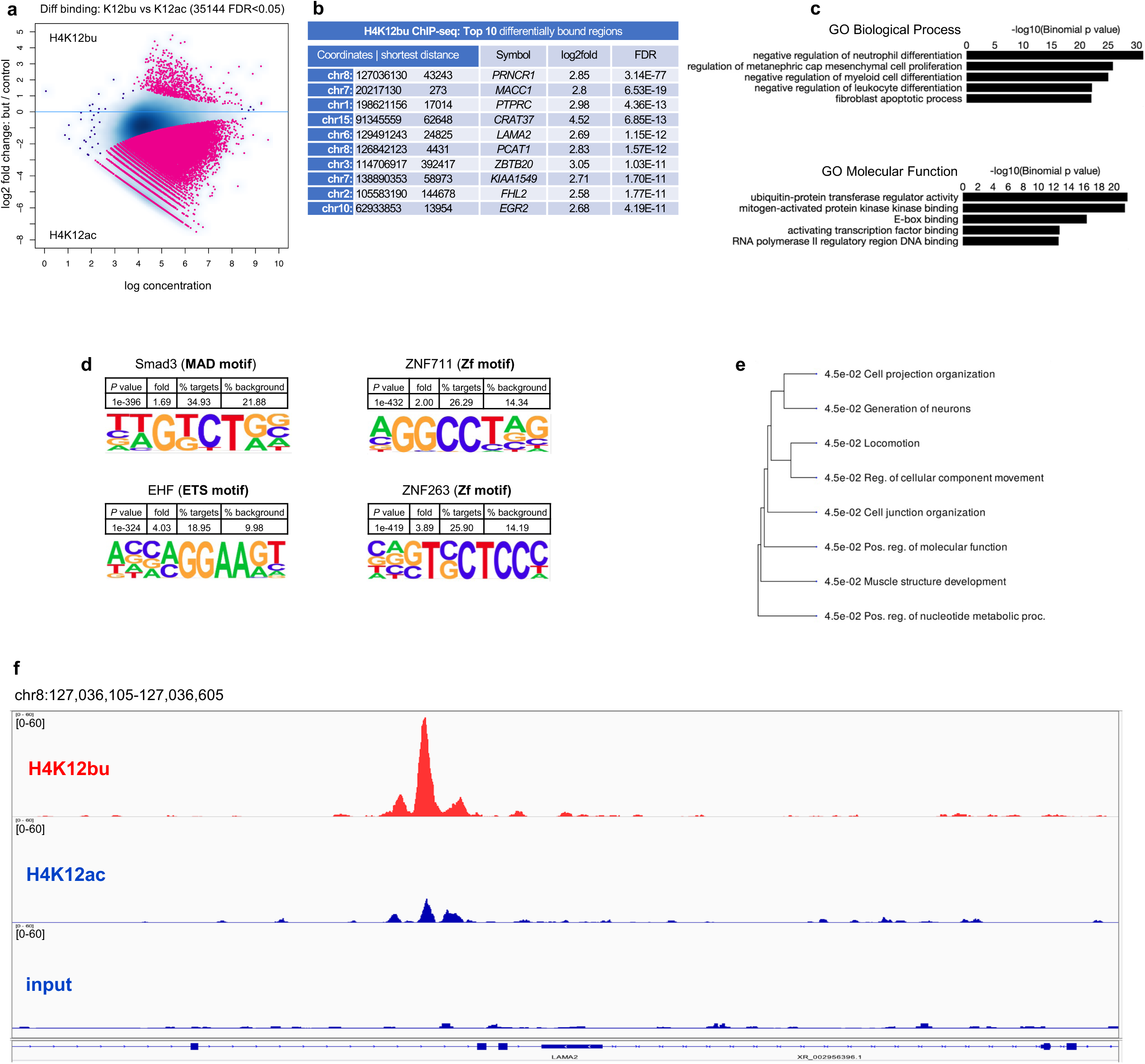
Genome-wide H4K12bu distribution. **a** H4K12bu vs H4K12ac differential binding following 1 mM butyrate supplementation. Sites identified as significantly differentially bound are shown in red. Differential binding was performed by DiffBind package with DESeq2 using a two-sided tests for both increased and decreased binding affinity between conditions followed by by multiple hypothesis testing and FDR correction. **b** Top ten differentially bound regions associated with H4K12bu sorted by false-discovery rate adjusted *P* value (FDR < 0.05). **c** Top GO ‘Biological Process’ and ‘Molecular Function’ terms associated with H3K18bu-bound *cis*-regulatory elements determined by GREAT against a whole genome background using a binomial test over genomic regions, followed by multiple hypothesis testing using FDR corrected *P* values (FDR < 0.05). **d** Differential motif analysis of H4K12bu vs H4K12ac peaks was analyzed by HOMER, using a one-sided hypergeometric test for overrepresentation (enrichment) of motifs in the target sequences compared to the background, followed by multiple hypothesis testing and FDR correction. **e** Hierarchical clustering tree summary of correlations among significant pathways in H4K12bu-associated annotated genes. Hierarchical clustering of the pathways was performed using ShinyGO. Pathways were clustered together based on shared genes and gene enrichment analysis was performed using two-sided Fisher’s exact test, and FDR correction was applied to adjust for multiple comparisons in the pathway analysis and hierarchical clustering. Size of dots indicates statistically significant FDR adjusted (FDR < 0.05) *P* values. **f** Signal tracks of 20 Kb-spanning *LAMA* region showing H4K12bu vs H4K12ac binding with input as background.

**Supplementary Fig. 10.**
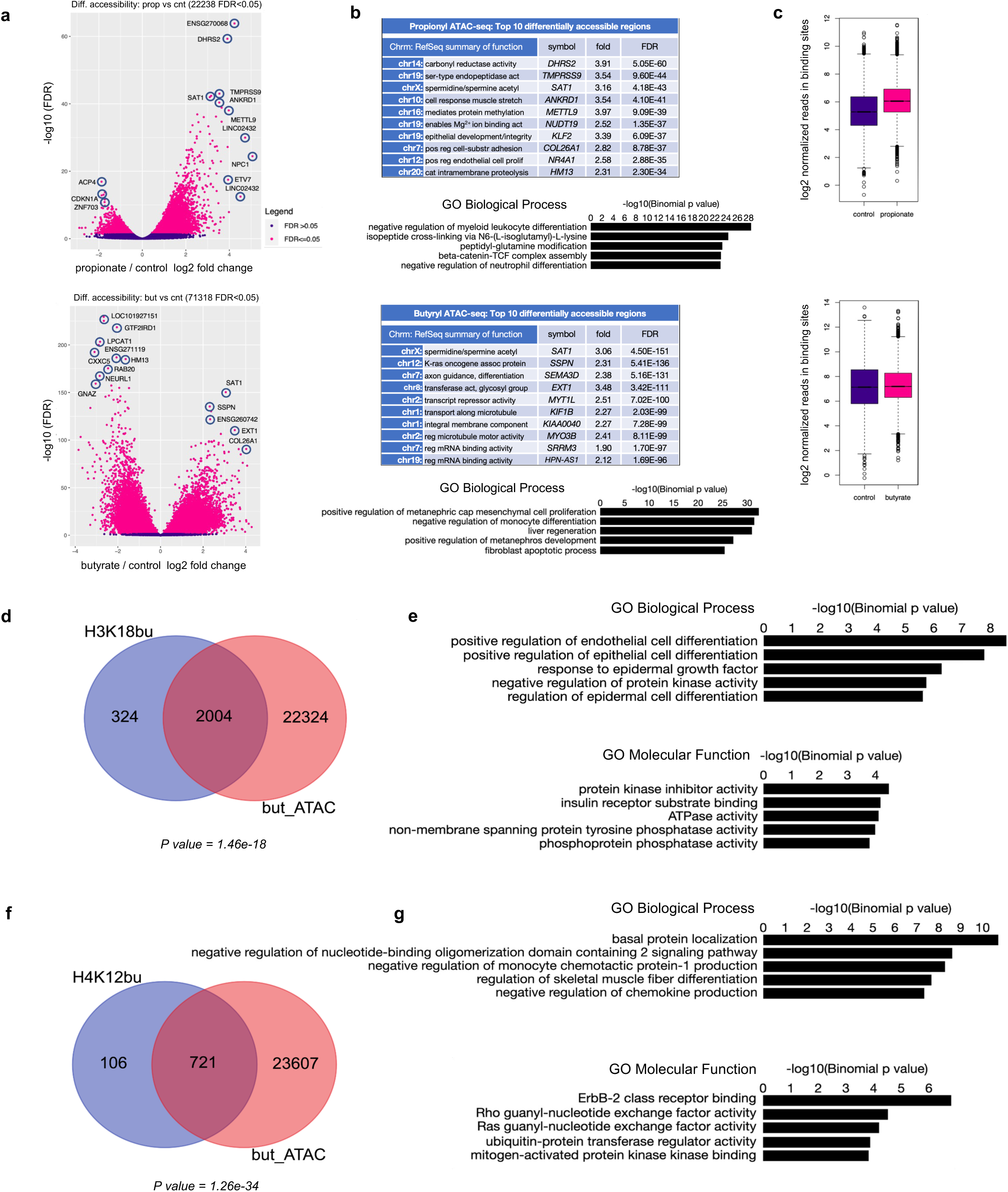
Propionyl and butyryl ATAC-seq and Kbu ChIP-seq and ATAC-seq integration. **a** Differential accessibility following propionate and butyrate supplementation. Sites identified as significantly differentially accessible are shown in red. n = 3 technical replicates for each condition. Differential accessibility was performed by DiffBind package with DESeq2 using a two-sided tests for both increased and decreased binding affinity between conditions followed by by multiple hypothesis testing and FDR correction. **b** Top ten differentially bound regions associated with propionate and butyrate treatment sorted by false-discovery rate adjusted *P* value (FDR < 0.05) and top GO ‘Biological Process’ terms determined by GREAT against a whole genome background using a binomial test over genomic regions, followed by multiple hypothesis testing using FDR corrected *P* values (FDR < 0.05). **c** Normalized reads in accessible sites following propionate and butyrate treatment. Box plots display: The minimum, first quartile (Q1, 25^th^ percentile), median, third quartile (Q3, 75^th^ percentile), and maximum. The bottom of the box is Q1 and the top of the box is Q3. The line within the box represents the median (50^th^ percentile) value. The whiskers extend to the most extreme data points within 1.5 times the IQR (interquartile range). **d, f** Overlap between Kbu bound genes and differentially accessible regions following 1 mM butyrate treatment. Significance of overlap determined by hypergeometric test-generated *P v*alue. **e, g** GO ‘Biological Process’ and ‘Molecular Function’ pathway terms associated with Kbu bound genomic coordinates that are also present in ATAC-seq data set sorted by binomial *P* value.

**Extended Fig. 1.**
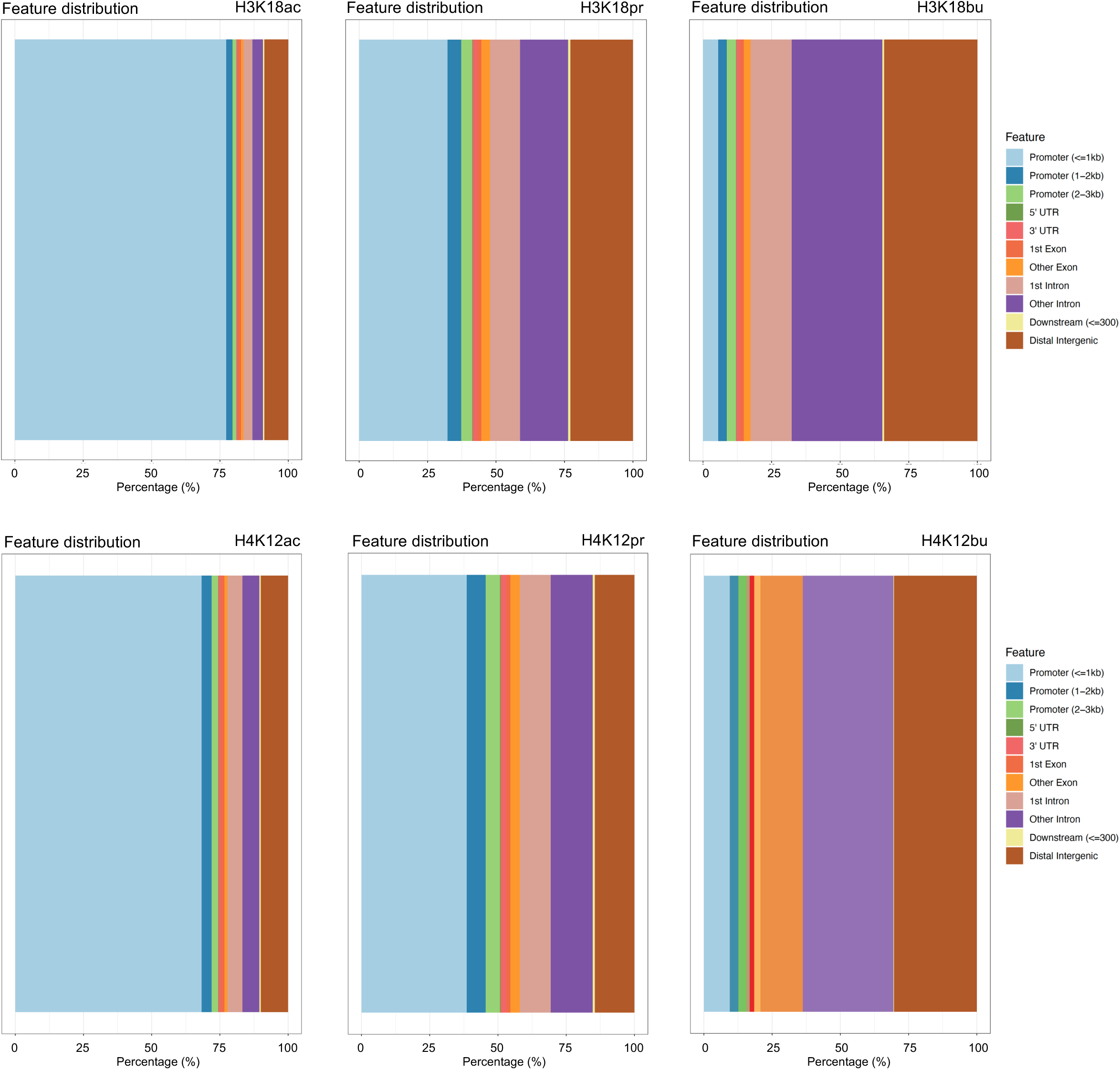
Feature distribution of H3K18ac/pr/bu (top panel) and H4K12ac/pr/bu (bottom panel) associated regions (+/- 3 Kb of TSS). H3K18ac/H4K12ac annotations were taken from ChIP-seq data that was generated without any treatment. H3K18pr/H4K12pr and H3K18bu/H4K12bu ChIP-seq experiments were performed following 10 mM NaPr, and 1 mM NaBu treatments, respectively. The x-axis provides the percentage of sites while colored regions represent distance from TSS. Results obtained using ChIPSeeker R package.

**Extended Fig. 2.**
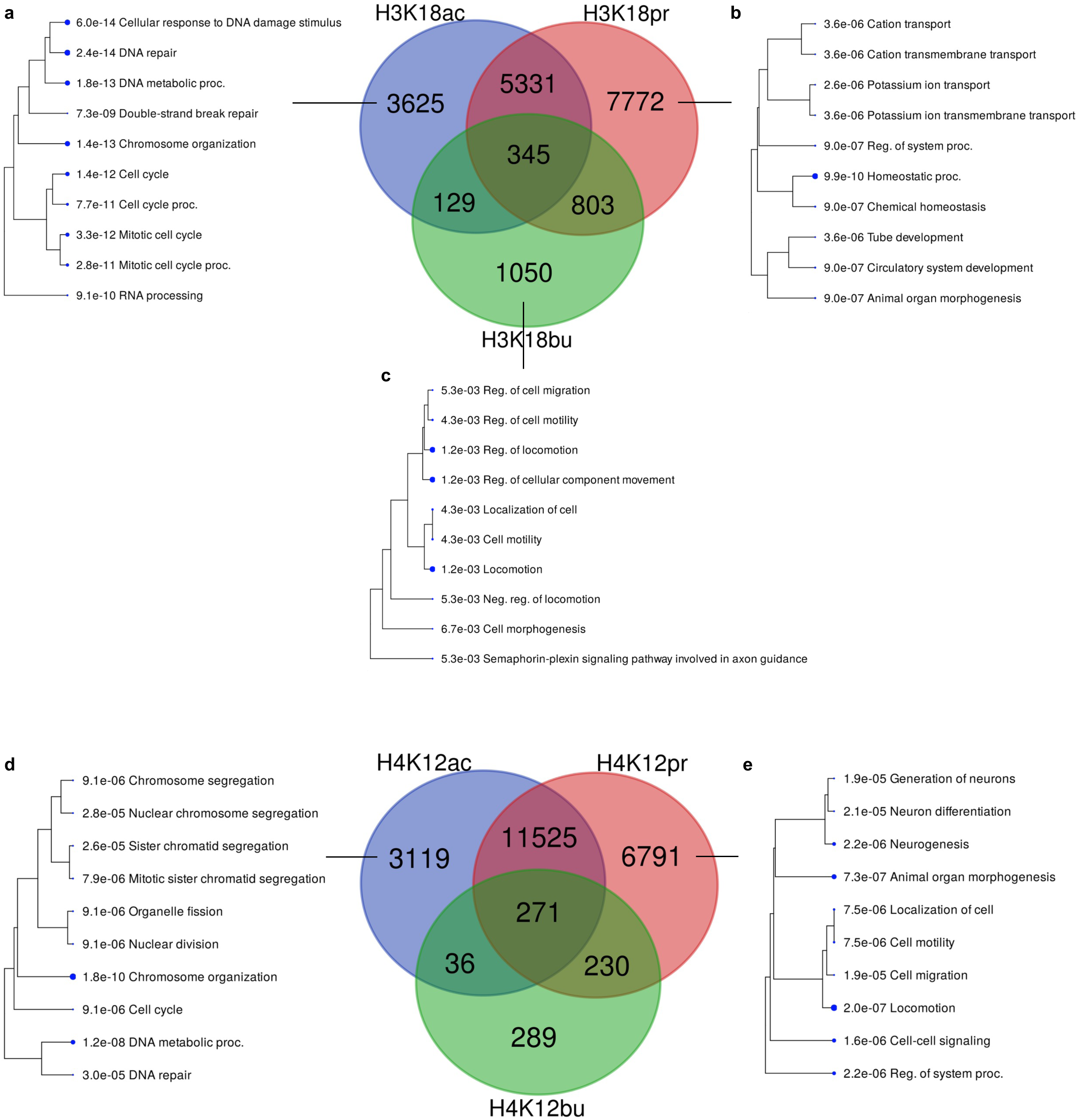
Kac/pr/bu annotated features overlap. **a, d** Hierarchical clustering of GO ‘Biological Process’ terms associated with Kac but not Kpr or Kbu. **b, e** Hierarchical clustering of GO ‘Biological Process’ terms associated with Kpr but not Kac or Kbu. **c** Hierarchical clustering of GO ‘Biological Process’ terms associated with Kbu but not Kac or Kpr. Hierarchical clustering of the pathways was performed using ShinyGO. Pathways were clustered together based on shared genes and gene enrichment analysis was performed using two-sided Fisher’s exact test, and FDR correction was applied to adjust for multiple comparisons in the pathway analysis and hierarchical clustering. Size of dots indicates statistically significant FDR adjusted (FDR < 0.05) *P* values.

**Extended Fig. 3.**
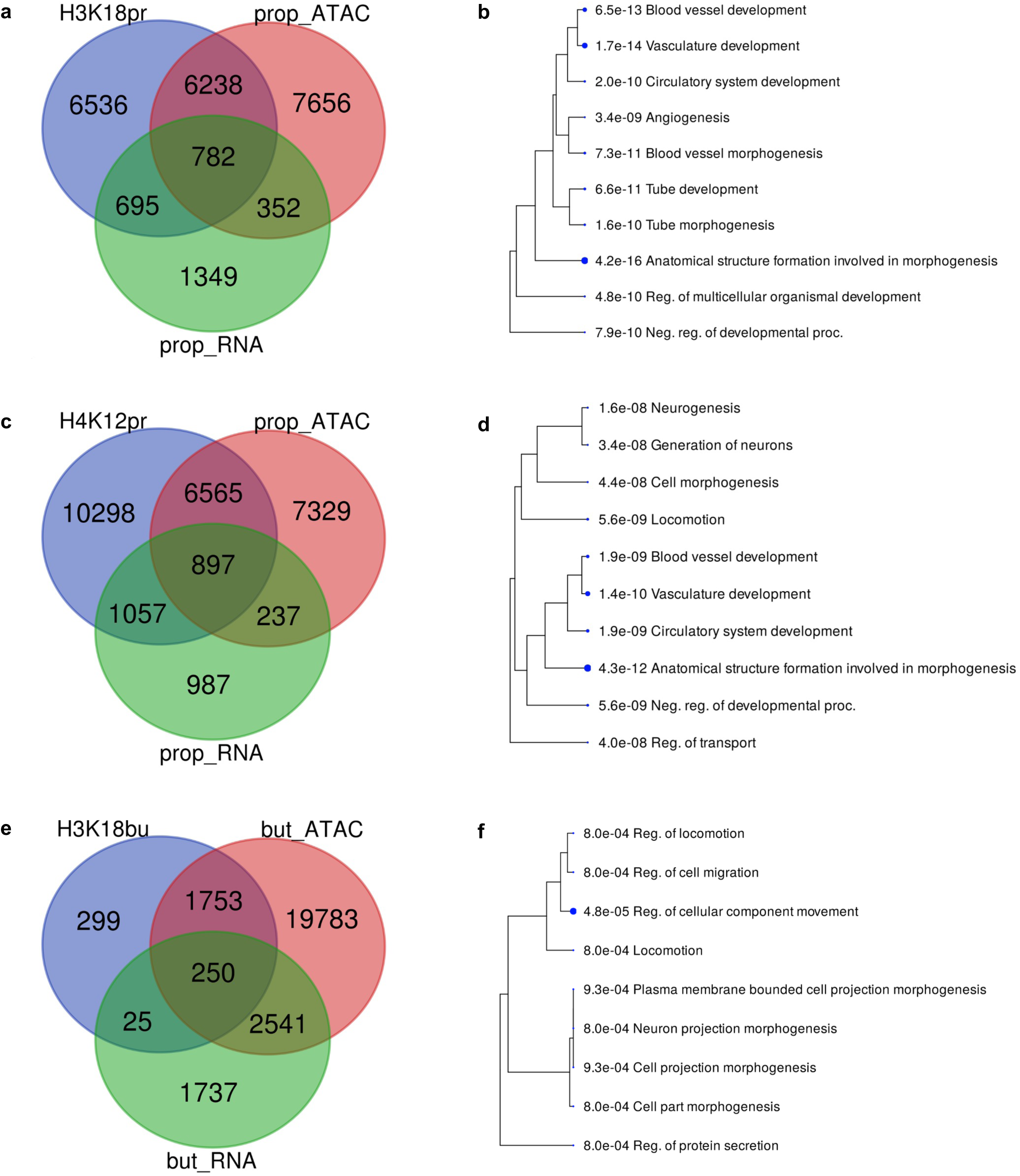
Kpr/bu and ATAC/RNA-seq shared annotated features. **a** H3K18pr, propionyl ATAC/RNA-seq shared annotated features. **b** Hierarchical cluttering of GO ‘Biological Process’ terms associated 782 shared features. **c** H4K12pr, propionyl ATAC/RNA-seq shared features. **d** Hierarchical clustering of GO ‘Biological Process’ terms associated with 897 shared features. Hierarchical clustering of the pathways was performed using ShinyGO. Pathways were clustered together based on shared genes and gene enrichment analysis was performed using two-sided Fisher’s exact test, and FDR correction was applied to adjust for multiple comparisons in the pathway analysis and hierarchical clustering. Size of dots indicates statistically significant FDR adjusted (FDR < 0.05) *P* values. **e** H3K18bu butyryl ATAC/RNA-seq shared features. **f** Hierarchical cluttering of GO ‘Biological Process’ terms associated with 250 shared features.

**Extended Fig. 4.**
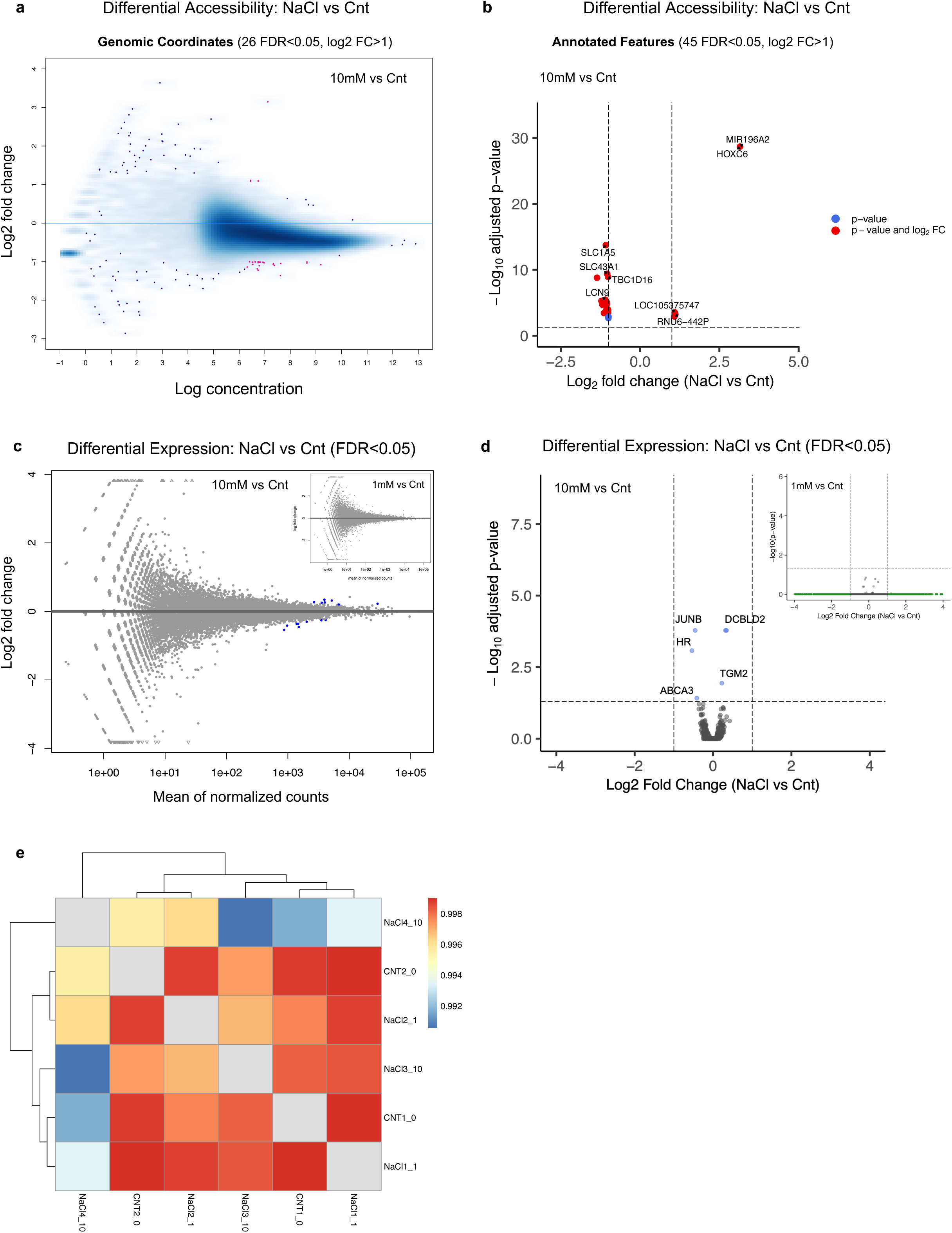
Effect of Na^+^ on differential accessibility and expression. **a** ATAC-seq MA plot of differential accessibility in 10 mM NaCl-treated vs untreated cells (SW480). Differential analysis was performed by DESeq2 using two-sided Wald tests to identify differentially expressed or accessible genes. The results were then adjusted for multiple comparisons using the FDR correction method. Sites identified as significantly differentially accessible (FDR<0.05, log2 FC>1) are shown in red. n = 3 experimental replicates for each condition. **b** Volcano plot of differential accessibility in 10 mM NaCl-treated vs untreated cells. **c** RNA-seq MA plot of differential expression in 10 mM NaCl treated vs untreated cells. Sites identified as significantly differentially expressed (FDR<0.05) are shown blue (*Insert*: 1 mM vs Cnt). **d** Volcano plot of differential expression in 10 mM NaCl-treated vs untreated cells. Sites identified as significantly differentially expressed by FDR only (FDR<0.05) are shown blue (*Insert*: 1 mM vs Cnt). **e** Correlation heatmap of RNA-seq data showing clustering replicates from 1 and 10 mM NaCl-treated vs untreated groups. Normalized measurement of the covariance between replicates is expressed by Pearson’s correlation coefficient.

**Extended Fig. 5.**
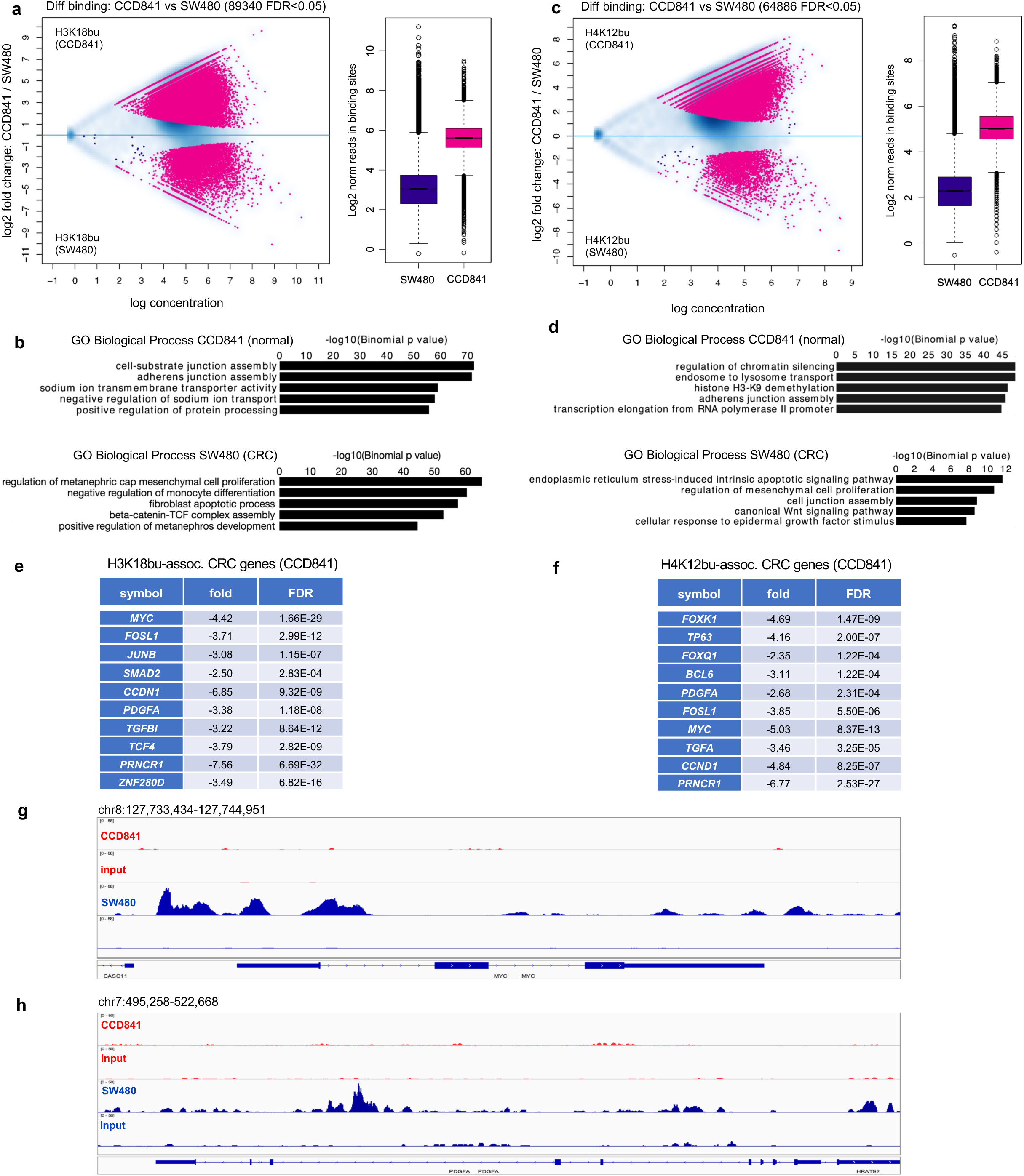
**a** H3K18bu differential binding in cancer (SW480) vs normal (CCD841) cells and normalized reads in H3K18bu-associated binding sites following 1 mM NaBu treatment. Sites identified as significantly differentially bound are shown in red. n = 3 technical replicates for each condition. Differential binding was performed by DiffBind package with DESeq2 using a two-sided tests for both increased and decreased binding affinity between conditions followed by multiple hypothesis testing and FDR correction (FDR < 0.05). Box plots display: The minimum, first quartile (Q1, 25^th^ percentile), median, third quartile (Q3, 75^th^ percentile), and maximum. The bottom of the box is Q1 and the top of the box is Q3. The line within the box represents the median (50^th^ percentile) value. The whiskers extend to the most extreme data points within 1.5 times the IQR (interquartile range). **b** Top GO Biological Process terms for H3K18bu-associated *cis*-regulatory elements in cancer vs normal cells determined by GREAT against a whole genome background using a binomial test over genomic regions, followed by multiple hypothesis testing using FDR corrected *P* values (FDR < 0.05). **c** H4K12bu differential binding in cancer vs normal cells and normalized reads in H4K12bu associated binding sites following 1 mM NaBu treatment. **d** Top GO Biological Process terms for H4K12bu-associated *cis*-regulatory elements in cancer vs normal cells. **e, f** CRC differentially bound genes associated with H3K18bu/H4K12bu. **g, h** Signal tracks showing differential binding in *MYC* and *PDGFA* regions in cancer vs normal cells.

**Extended Fig. 6.**
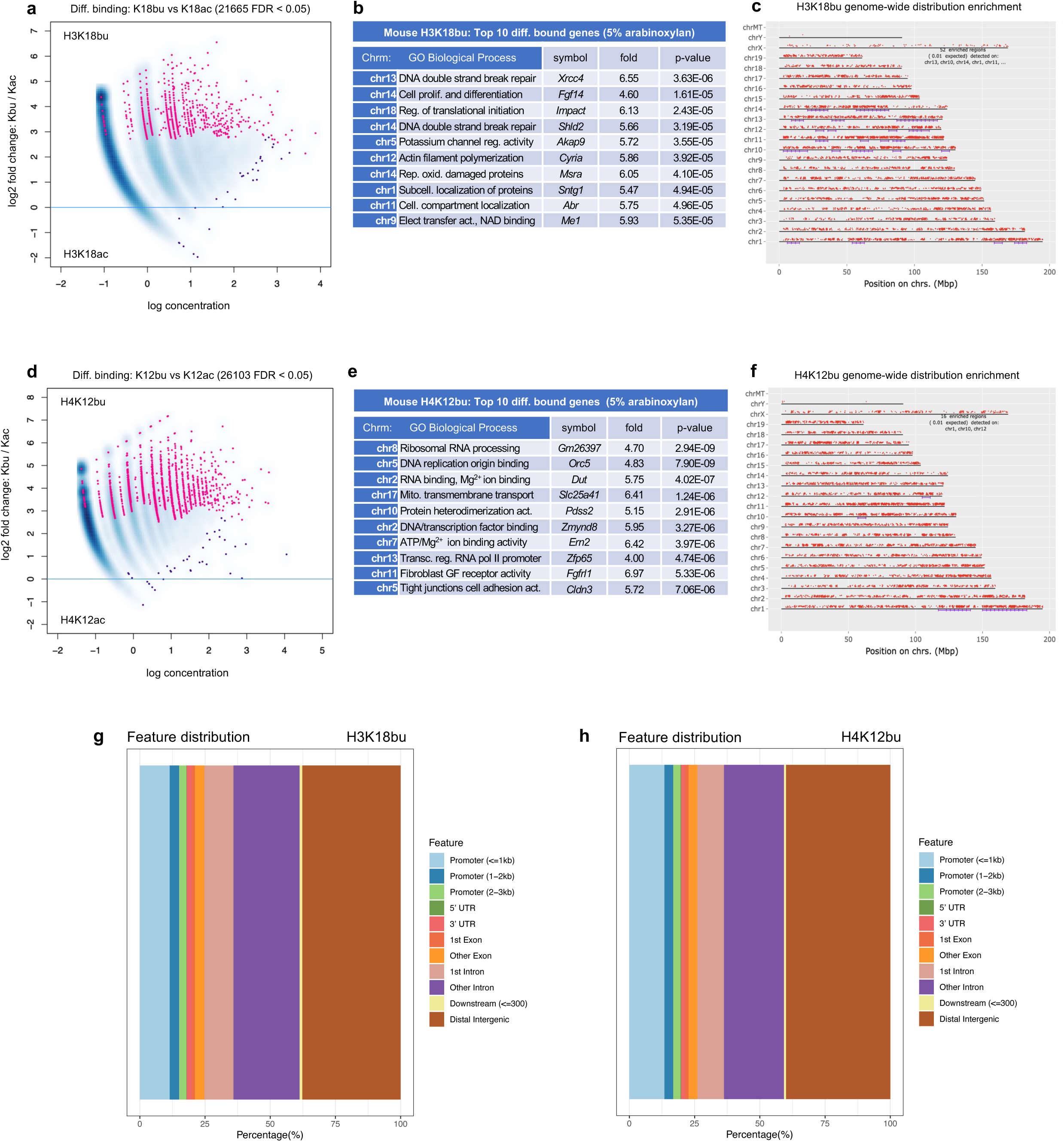
H3K18bu vs ac and H4K12bu vs ac differential binding from mouse intestine on HSF + 5 % arabinoxylan diet. **a, b, d, e** Sites identified as significantly differentially bound are shown in red. n = 3 biological replicates for each condition. Differential binding was performed by DiffBind package with DESeq2 using a two-sided tests for both increased and decreased binding affinity between conditions followed by multiple hypothesis testing and FDR correction. **c, f** Chromosomal positions of H3K18bu and H4K12bu-associated regions represented by red dots. Purple lines represent statistically significant enrichment compared to input. The genome was scanned with a sliding window (size 6 Mb) further subdivided into 2 equal-sized steps for sliding. Within each window a hypergeometric test was used to test for enrichment over all protein-coding genes in the genome. FDR adjusted *P* value cutoff for window was 1E-05. Gene chromosomal mapping performed using ShinyGO. **g, h** Feature distribution of H3K18bu and H4K12bu bound regions (+/- 3 Kb of TSS). The x-axis provides the percentage of sites while colored regions represent distance from TSS. Results obtained using ChIPSeeker R package.

**Extended Fig. 7.**
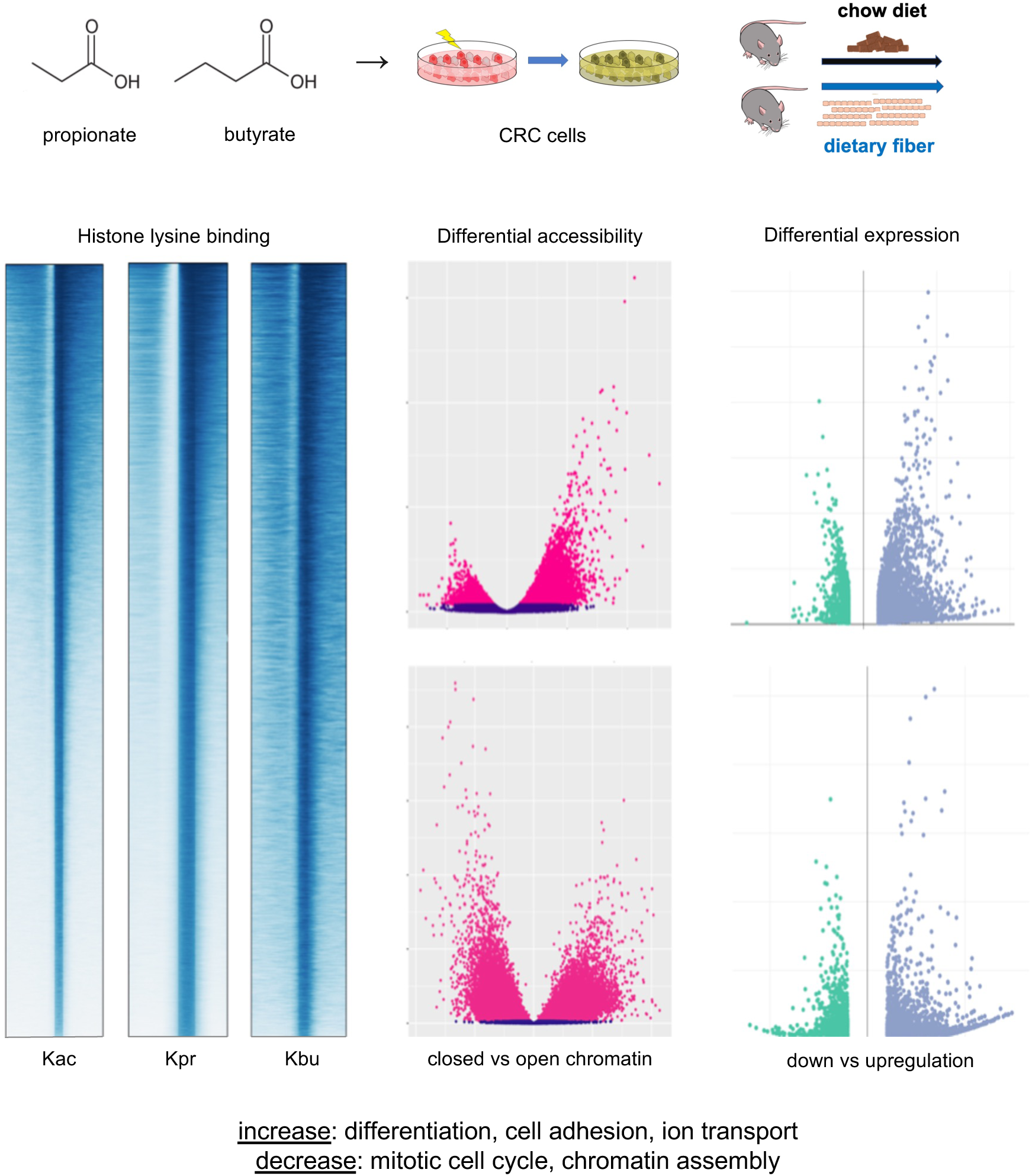
Overview of SCFAs propionate and butyrate as regulatory elements affecting histone binding, chromatin accessibility and gene expression. Upper panels: *In vitro* cellular perturbations and *in vivo* fiber supplementation experimental designs. Bottom panels: Changes in TSS distribution profiles, differential chromatin accessibility and expression following SCFA supplementation. Top and Bottom volcano plots showing differential accessibility and expression following propionate and butyrate supplementation, respectively.

