## Supplementary material for "Short-chain fatty acid metabolites propionate and butyrate are unique epigenetic regulatory elements linking diet, metabolism and gene expression": Nshanian_et_al_Supplemental_Figures

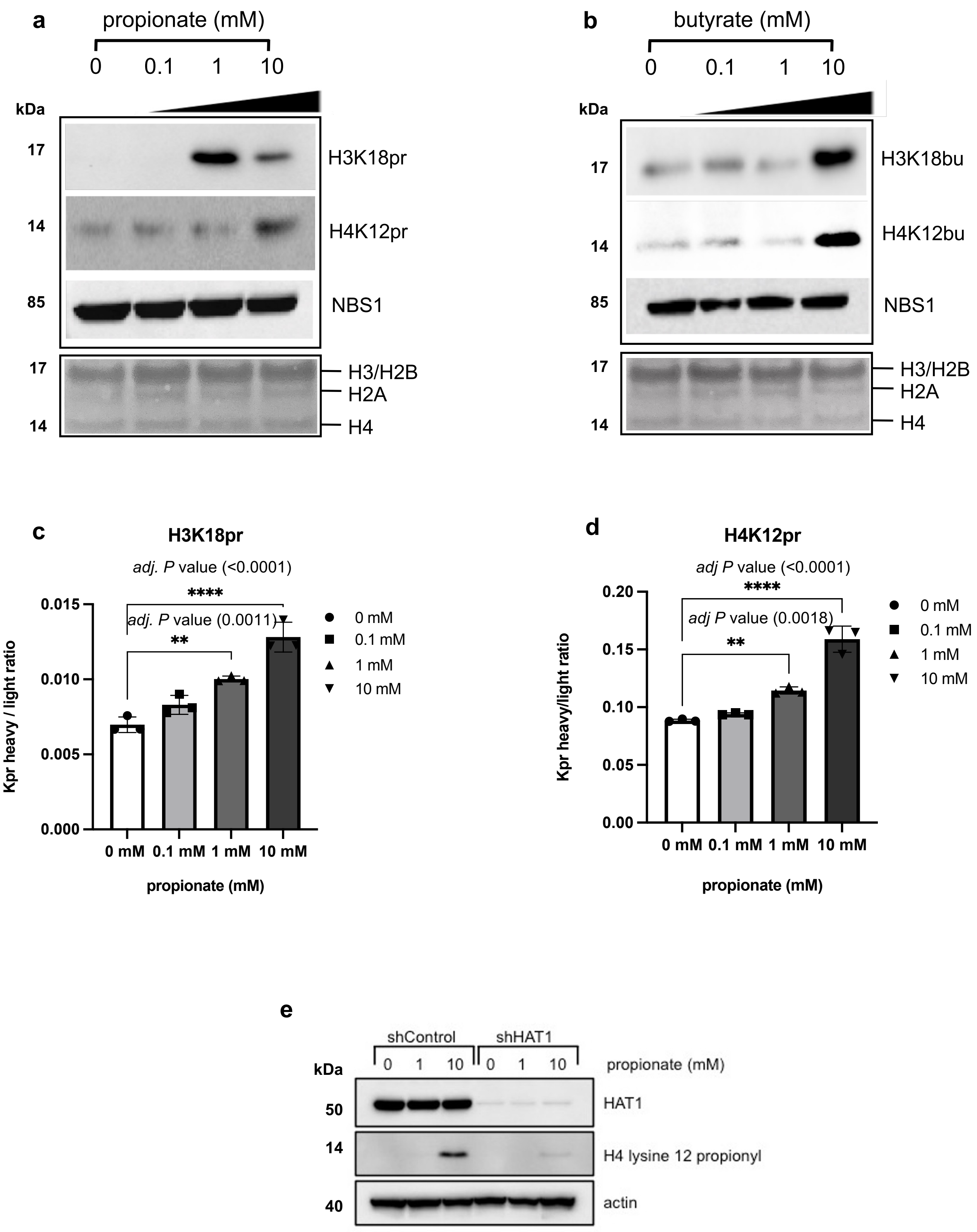


**Supplementary Fig. 1** | Immunoblots of acid-extracted **a** H3K18/H4K12pr and **b** H3K18/H4K12bu histone marks at 0 – 10 mM sodium propionate/butyrate treatments with anti-NBS1 as control (top). SDS-PAGE of histones H2A/B, H3/H4 (bottom). Dose-dependent $^{13}$C-propionate incorporation into H3 and H4 as measured by increases in the heavy/light propionyl lysine containing peptides. Heavy/light propionyl lysine containing peptides representing **c** H3K18pr and **d** H4K12pr levels at 0, 0.1, 1, and 10 mM $^{13}$C-propionate supplementation (mean ± SD, n = 3). Multiple comparisons by ordinary, one-way ANOVA followed by hypothesis testing using the Bonferroni correction method with 0.05 *P* value cutoff. ***P* < 0.01, **** *P* < 0.0001 **e** Depletion of HAT1 diminishes incorporation of propionate into the H4 lysine 12 site.

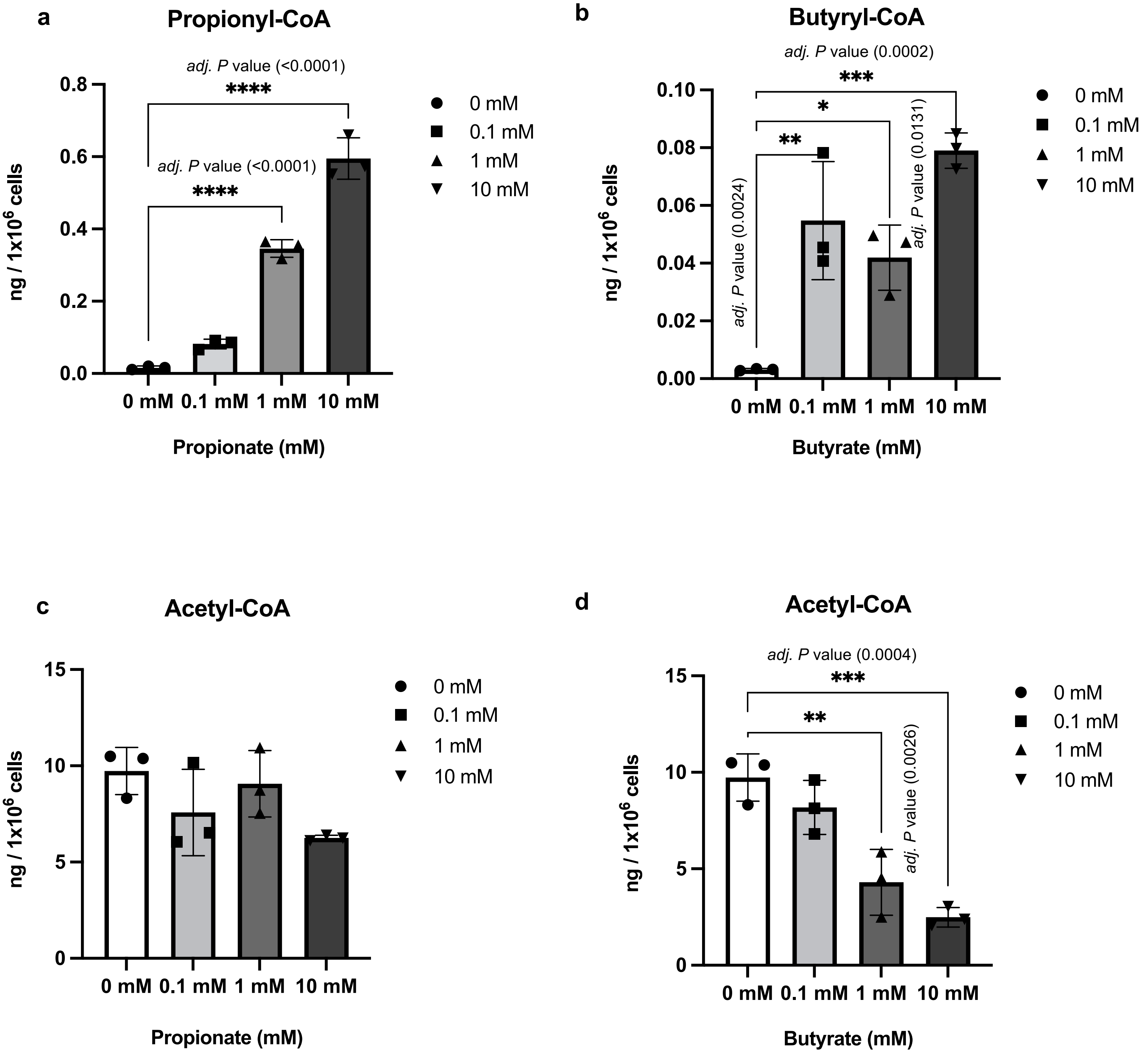


**Supplementary Fig. 2 |** Quantitative analysis of acyl-CoA levels by LC-MS/MS following SCFA supplementation. **a** Dose-dependent increases in **a** Propionyl-CoA and **b** Butyryl-CoA levels upon increasing propionate and butyrate supplementation at 0, 0.1, 1, and 10 mM. Acetyl-CoA levels upon increasing **c** propionate and **d** butyrate supplementation. Quantitative analysis was done by multiple reaction monitoring (MRM) using an internal standard approach. Calculated Acyl-Co concentrations in each sample were normalized to the number of cells (mean ± SD, n = 3). Multiple comparisons by ordinary, one-way ANOVA using hypothesis testing followed by Bonferroni correction method with 0.05 *P* value cutoff. **P* < 0.05, ***P* < 0.01, ****P* < 0.001, *****P* < 0.0001.

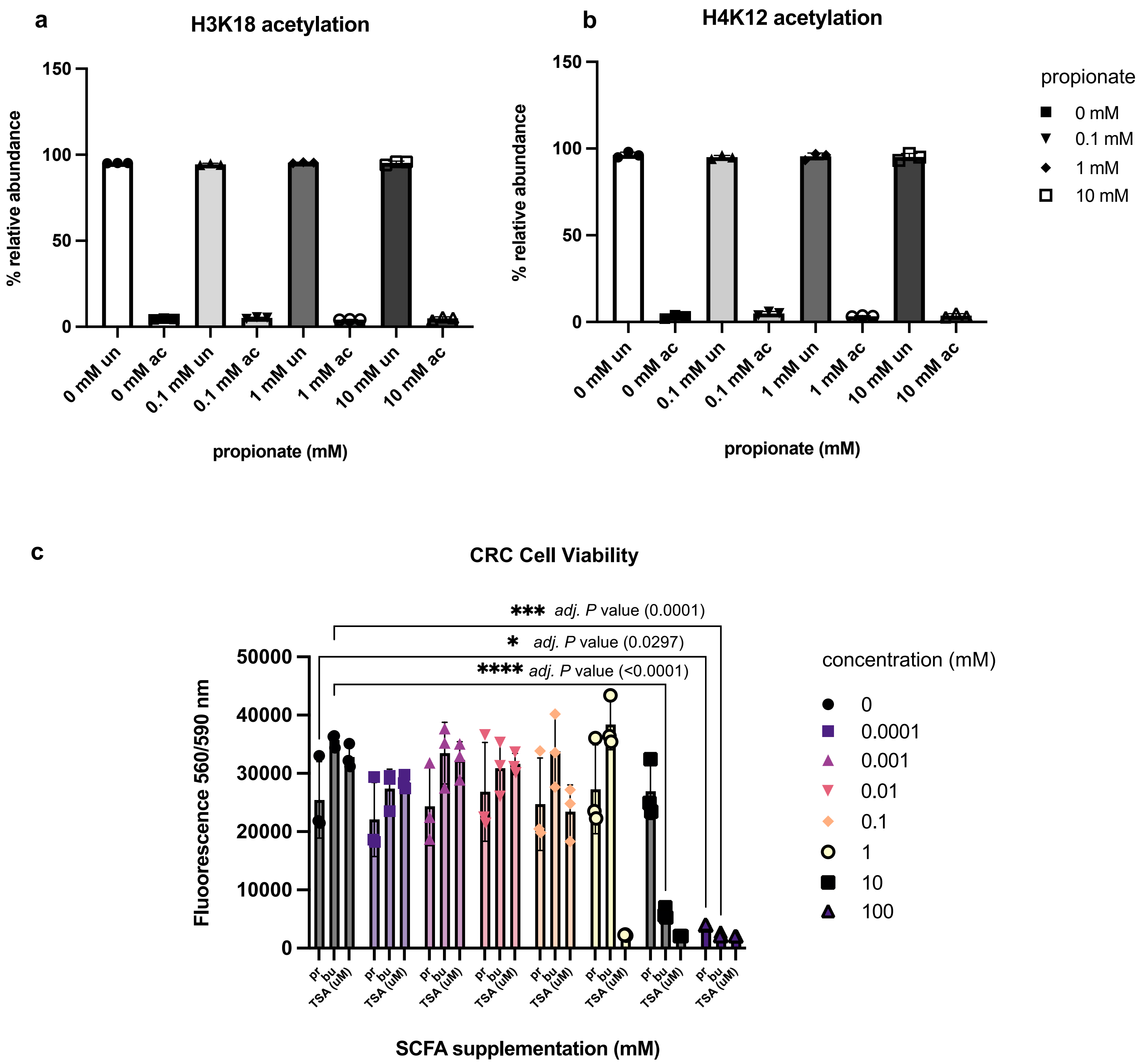


**Supplementary Fig. 3** | H3 and H4 acetylated vs unmodified states as a function of propionate supplementation. Relative abundances of acetylated vs unmodified states on **a** H3K18 **b** H4K12 **c** H3K9 and **d** H3K23 following 0, 0.1, 1, and 10 mM propionate supplementation (mean ± SD, n = 3). **e** CRC cell viability as a function of NaPr and NaBu supplementation over 72 hrs. as measured by CellTiter-Blue® fluorescence assay. TSA, a known HDAC inhibitor with IC50 ~ 2 nM was used as a negative control on a µM level. (mean ± SD, n = 3). Multiple comparisons by two-way ANOVA using statistical hypothesis testing followed by Bonferroni correction method with 0.05 *P* value cutoff. **P* < 0.05, ***p < 0.001, ****p < 0.0001.

**a**

| Enrichment FDR | nGenes | Pathway Genes | Fold Enrichment | Pathways |
|---|---|---|---|---|
| 7.9E-08 | 56 | 86 | 1.9 | Colorectal cancer |
| 6.1E-09 | 82 | 135 | 1.8 | Apoptosis |
| 3.6E-10 | 95 | 157 | 1.8 | Hippo signaling pathway |
| 1.5E-08 | 110 | 200 | 1.6 | Focal adhesion |
| 1.5E-08 | 111 | 202 | 1.6 | Proteoglycans in cancer |
| 8.0E-08 | 115 | 217 | 1.6 | Reg. of actin cytoskeleton |
| 7.7E-11 | 174 | 331 | 1.6 | Human papillomavirus infection |
| 9.3E-08 | 127 | 246 | 1.5 | Shigellosis |
| 6.7E-08 | 148 | 294 | 1.5 | MAPK signaling pathway |
| 4.7E-13 | 265 | 530 | 1.5 | Pathways in cancer |

**c**

| H3K18pr CRC targets | | | Prop RNA-seq | |
|---|---|---|---|---|
| symbol | fold | FDR | fold | p-adj |
| *CTNNB1* | 3.00 | 2.05E-11 | -0.61 | 4.65E-09 |
| *TCF20* | 3.01 | 9.54E-11 | -0.34 | 0.080547 |
| *LEF1* | 2.72 | 9.49E-06 | -1.26 | 2.65E-05 |
| *CCND1* | 3.85 | 4.08E-21 | -0.46 | 1.56E-05 |
| *MYC* | 2.08 | 9.55E-06 | 1.93 | 4.66E-67 |
| *FOS* | 2.99 | 1.97E-12 | 1.56 | 7.67E-36 |
| *JUN* | 2.67 | 3.03E-06 | 1.57 | 8.08E-09 |

**b**

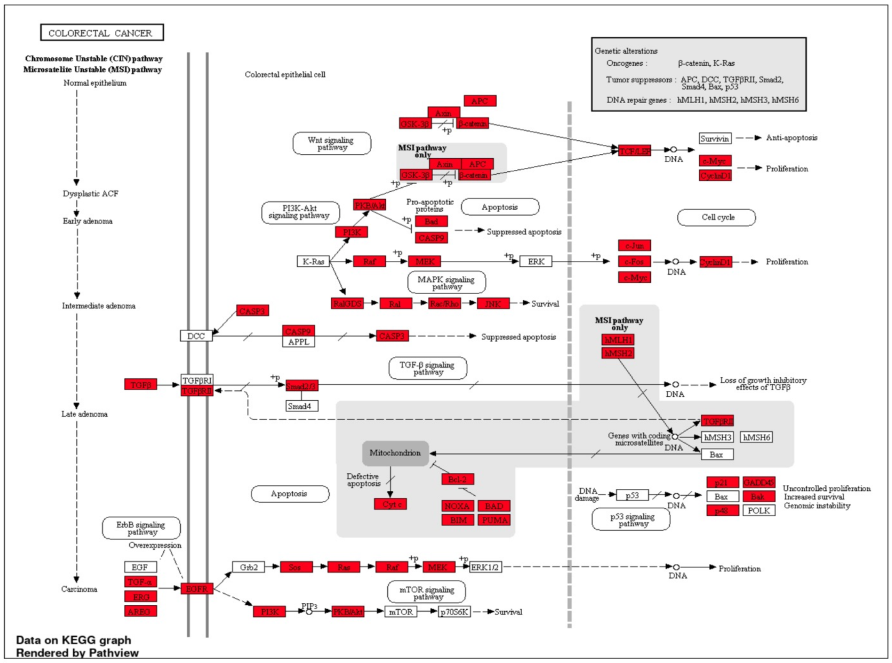


**d**

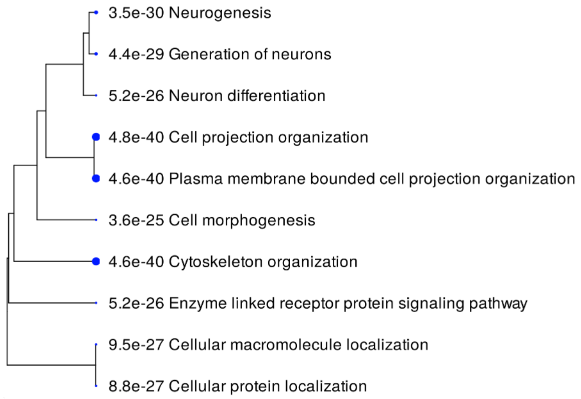


**e**

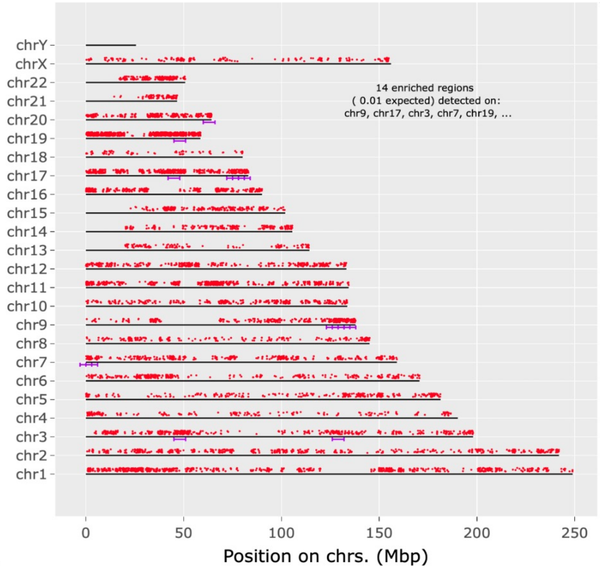


**Supplementary Fig. 4 |** KEGG pathway analysis of H3K18pr-associated genes. **a** Top ten pathways with their number of genes and log2 fold enrichment. FDR is calculated from a nominal *P* value obtained from a hypergeometric test (FDR < 0.05). Fold enrichment is calculated as percentage of H3K18pr differentially bound genes associated with a pathway divided by the corresponding percentage in input. **b** CRC KEGG pathway enrichment by Pathview. Genes that are overrepresented compared to input are in red. **c** H3K18pr-associated differential binding of key CRC genes and log2 fold changes in their expression levels, as determined by differential RNA-seq of 10 mM treated vs. untreated conditions. n = 3 experimental replicates per condition (FDR < 0.05). **d** Hierarchical clustering tree summary of correlations among significant pathways in H3K18pr-associated annotated genes. Hierarchical clustering of the pathways was performed using ShinyGO. Pathways were clustered together based on shared genes and gene enrichment analysis was performed using two-sided Fisher's exact test, and FDR correction was applied to adjust for multiple comparisons in the pathway analysis and hierarchical clustering. Size of dots indicates statistically significant FDR adjusted (FDR < 0.05) *P* values. **e** Chromosomal position of H3K18pr-associated regions represented by red dots. Purple lines represent statistically significant enrichment compared to input. The genome was scanned with a sliding window (size 6 Mb) further subdivided into 2 equal-sized steps for sliding. Within each window a hypergeometric test was used to test for enrichment over input. FDR-adjusted *P* value cutoff for window was 1E-05. Chromosomes may be partly shown due to scaling to last genes location. Gene chromosomal mapping was performed using ShinyGO.

**a**

| Enrichment FDR | nGenes | Pathway Genes | Fold Enrichment | Pathways |
|---|---|---|---|---|
| 4.6E-11 | 104 | 134 | 1.6 | Ribosome |
| 4.5E-10 | 116 | 157 | 1.6 | Hippo signaling pathway |
| 2.0E-08 | 103 | 142 | 1.5 | Spinocerebellar ataxia |
| 9.4E-09 | 120 | 169 | 1.5 | Protein processing in endoplasmic reticulum |
| 4.5E-10 | 203 | 306 | 1.4 | Huntington disease |
| 2.0E-08 | 167 | 252 | 1.4 | Endocytosis |
| 2.3E-11 | 240 | 363 | 1.4 | Amyotrophic lateral sclerosis |
| 1.9E-12 | 307 | 475 | 1.4 | Pathways of neurodegeneration |
| 3.4E-09 | 243 | 383 | 1.3 | Alzheimer disease |
| 7.6E-09 | 838 | 1527 | 1.2 | Metabolic pathways |

**c**

| H4K12pr CRC targets | | | Prop RNA-seq | |
|---|---|---|---|---|
| symbol | fold | FDR | fold | p-adj |
| *CTNNB1* | 3.17 | 5.90E-15 | -0.61 | 4.65E-09 |
| *TCF20* | 3.26 | 0.000136 | -0.34 | 0.080547 |
| *LEF1* | 2.01 | 0.0019 | -1.26 | 2.65E-05 |
| *CCND1* | 2.8 | 5.54E-14 | -0.46 | 1.56E-05 |
| *MYC* | 2.4 | 2.09E-11 | 1.93 | 4.66E-67 |
| *FOS* | 3.05 | 1.28E-07 | 1.56 | 7.67E-36 |
| *JUN* | 3.32 | 7.54E-18 | 1.57 | 8.08E-09 |

**b**

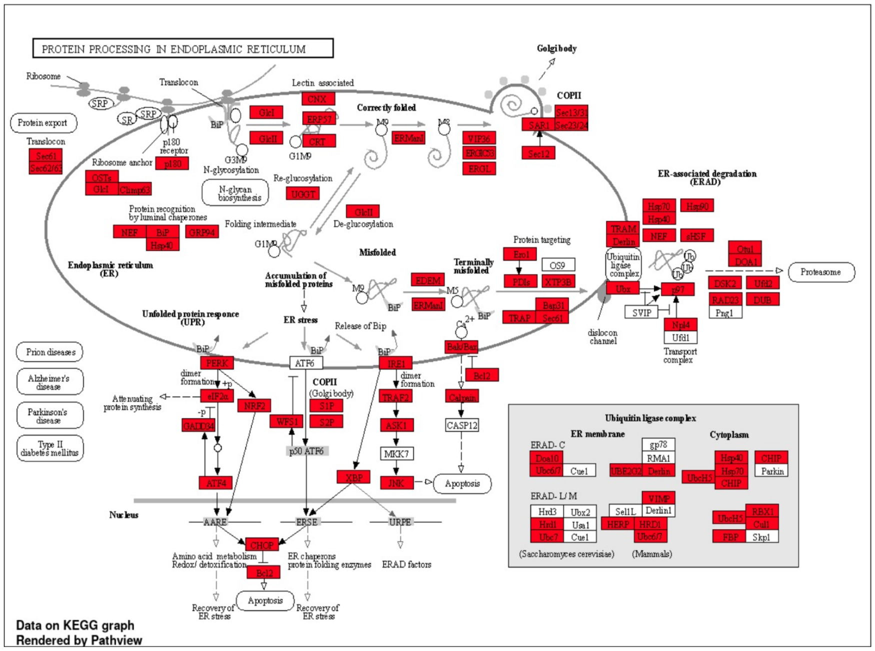


**d**

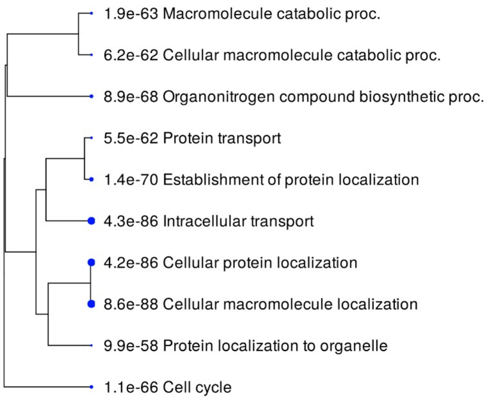


**e**

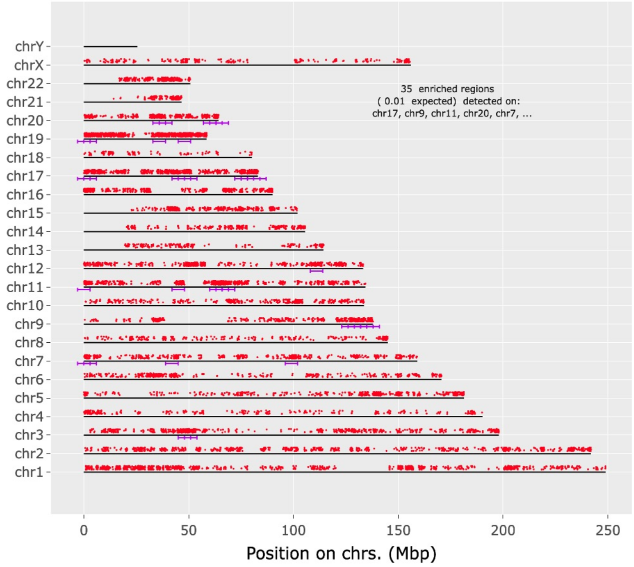


**Supplementary Fig. 5** | KEGG pathway analysis of H4K12pr associated genes. **a** Top ten pathways with their number of genes and log2 fold enrichment. FDR is calculated from a nominal *P* value obtained from a hypergeometric test (FDR < 0.05). Fold enrichment is calculated as percentage of H4K12pr differentially-bound genes associated with a pathway divided by the corresponding percentage in input. **b** Protein Processing in ER KEGG pathway enrichment by Pathview. Genes that are overrepresented compared to input are in red. **c** H4K12pr-associated differential binding of key CRC genes and log2 fold changes in their expression levels, as determined by differential RNA-seq of 10 mM treated vs. untreated conditions. n = 3 experimental replicates per condition (FDR < 0.05). **d** Hierarchical clustering tree summary of correlations among significant pathways in H3K18pr-associated annotated genes. Hierarchical clustering of the pathways was performed using ShinyGO. Pathways were clustered together based on shared genes and gene enrichment analysis was performed using two-sided Fisher's exact test, and FDR correction was applied to adjust for multiple comparisons in the pathway analysis and hierarchical clustering. Size of dots indicates statistically significant FDR adjusted (FDR < 0.05) *P* values. **e** Chromosomal position of H3K18pr-associated regions represented by red dots. Purple lines represent statistically significant enrichment compared to input. The genome was scanned with a sliding window (size 6 Mb) further subdivided into 2 equal-sized steps for sliding. Within each window a hypergeometric test was used to test for enrichment over input. FDR-adjusted *P* value cutoff for window was 1E-05. Chromosomes may be partly shown due to scaling to last genes location. Gene chromosomal mapping was performed using ShinyGO.

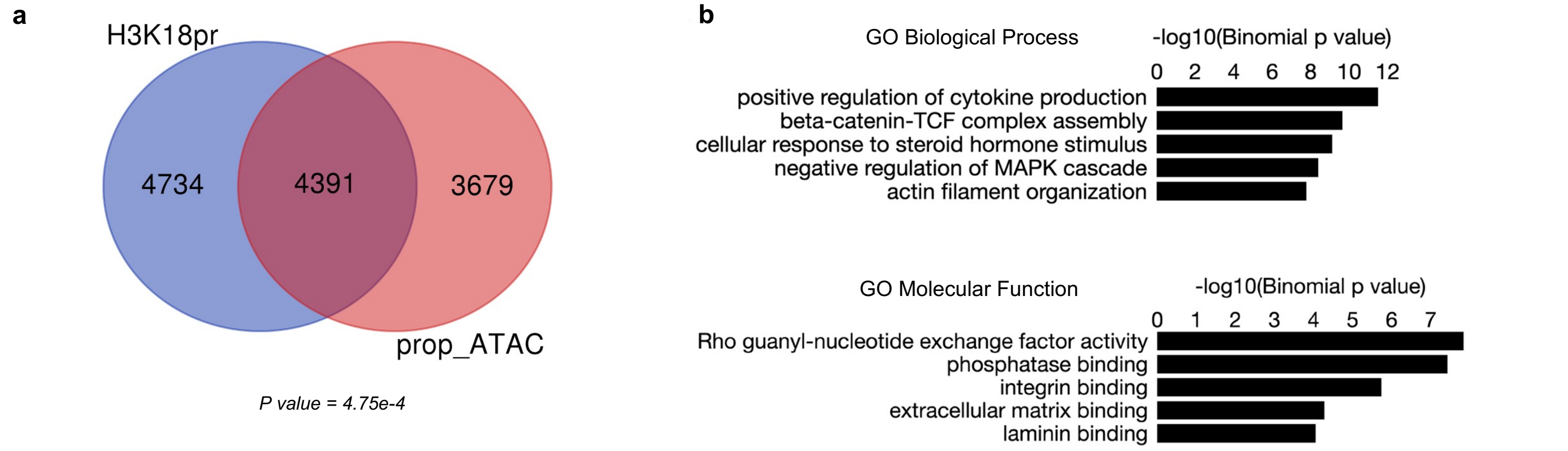


**c**

| H3K18pr ChIP CRC targets | | | Prop ATAC-seq | |
|---|---|---|---|---|
| symbol | fold | FDR | fold | FDR |
| ***CTNNB1*** | 3.00 | 2.05E-11 | 0.70 | 0.0086 |
| ***TCF20*** | 3.01 | 9.54E-11 | 2.24 | 1.124E-06 |
| ***LEF1*** | 2.72 | 9.49E-06 | 1.32 | 0.00243 |
| ***CCND1*** | 3.85 | 4.08E-21 | 1.48 | 5.224E-07 |
| ***MYC*** | 2.08 | 9.55E-06 | 0.81 | 0.000143 |
| ***FOS*** | 2.39 | 1.97E-12 | 1.46 | 8.17E-15 |
| ***JUN*** | 2.67 | 3.03E-06 | 1.89 | 1.76E-16 |

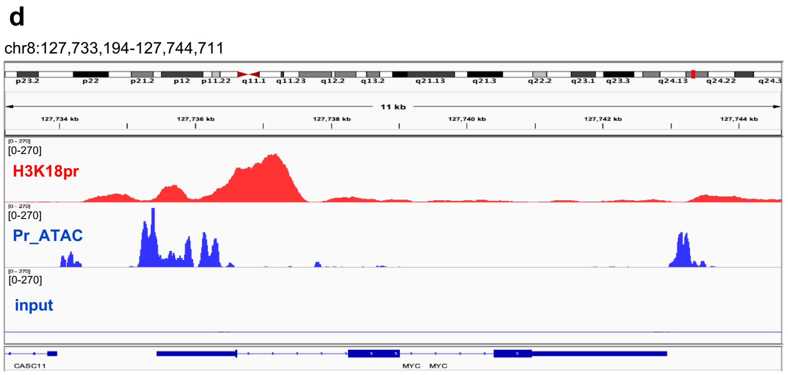


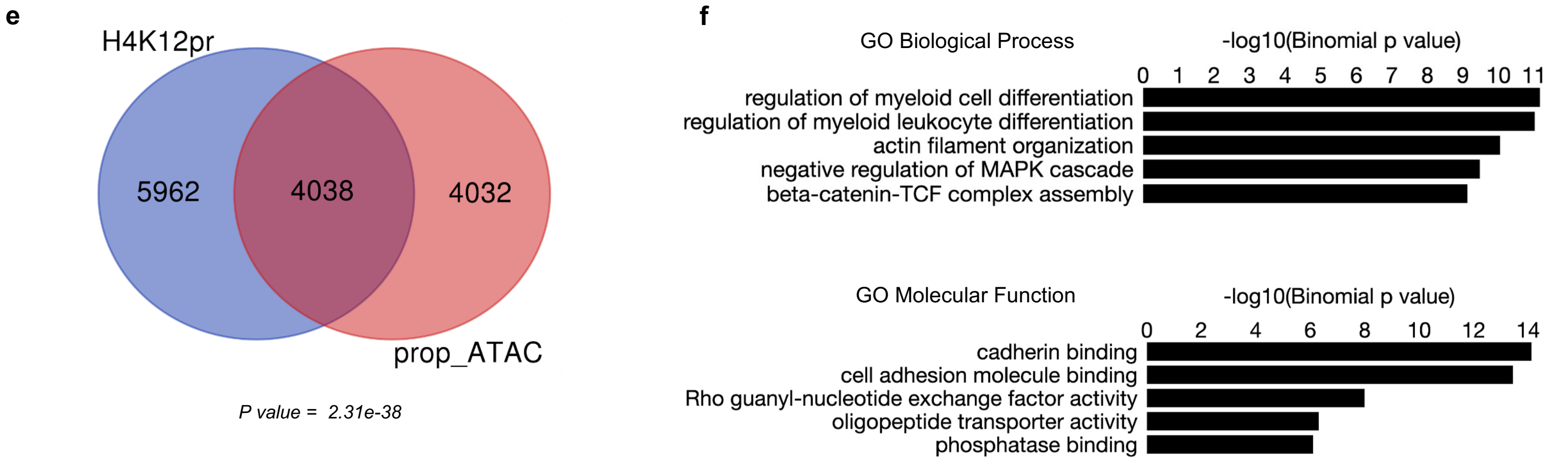


**g**

| H4K12pr ChIP CRC targets | | | Prop ATAC-seq | |
|---|---|---|---|---|
| symbol | fold | FDR | fold | FDR |
| ***CTNNB1*** | 3.17 | 5.90E-15 | 0.70 | 0.0086 |
| ***TCF20*** | 3.26 | 0.000136 | 2.24 | 1.124E-06 |
| ***LEF1*** | 2.01 | 0.0019 | 1.32 | 0.00243 |
| ***CCND1*** | 2.8 | 5.54E-14 | 1.48 | 5.224E-07 |
| ***MYC*** | 2.4 | 2.09E-11 | 0.81 | 0.000143 |
| ***FOS*** | 3.05 | 1.28E-07 | 1.46 | 8.17E-15 |
| ***JUN*** | 3.32 | 7.54E-18 | 1.89 | 1.76E-16 |

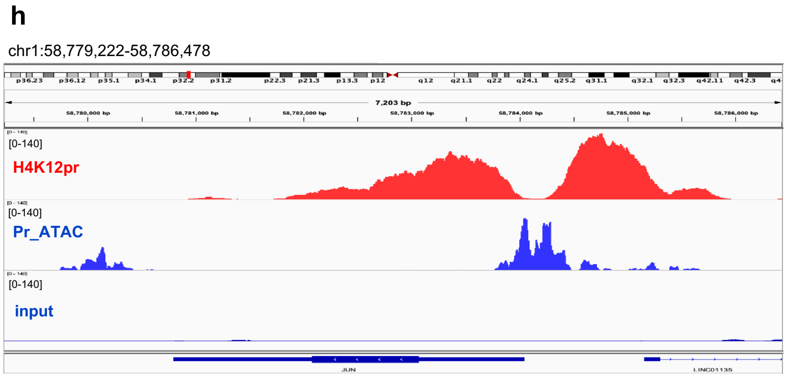


**Supplementary Fig. 6** | H3K18/H4K12pr ChIP-seq and propionyl ATAC-seq integration. **a, e** Overlap between TSS-proximal regions (+/- 1 Kb) for H3K18pr and H4K12pr differentially bound genes by ChIP-seq and differentially accessible genes following 10 mM propionate supplementation. Significance of overlap determined by hypergeometric test-generated *P* value. **b, f** GO 'Biological Process' and 'Molecular Function' pathway terms associated with H3K18pr and H4K12pr differentially bound genomic coordinates that are also present in ATAC-seq sorted by binomial *P* value. **c, g** Log2 fold change and FDR-adjusted *P* value in CRC-relevant gene targets associated with H3K18/H4K12pr ChIP-seq and propionyl ATAC-seq. **d, h** Signal tracks for *MYC* and *JUN* regions showing ChIP-seq and ATAC-seq profiles with input as background.

**a**

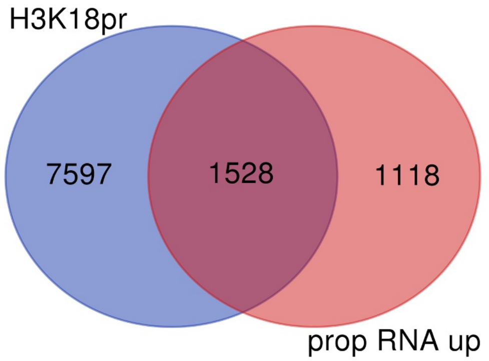


*P value = 7.10e-154*

**b**

| Enrichment FDR | nGenes | Fold Enrichment | Pathways |
|---|---|---|---|
| 2.1E-19 | 159 | 2.2 | Cell morphogenesis |
| 1.4E-20 | 214 | 2 | Generation of neurons |
| 9.3E-19 | 194 | 2 | Neuron differentiation |
| 2.0E-19 | 207 | 2 | Cell migration |
| 8.4E-21 | 228 | 2 | Neurogenesis |
| 9.3E-19 | 210 | 1.9 | Plasma membrane bounded cell projection organization |
| 2.0E-19 | 224 | 1.9 | Cell motility |
| 2.0E-19 | 224 | 1.9 | Localization of cell |
| 1.5E-18 | 213 | 1.9 | Cell projection organization |
| 1.1E-20 | 247 | 1.9 | Locomotion |

**c**

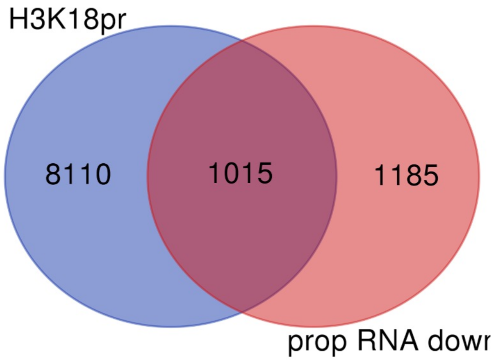


*P value = 3.29e-12*

**d**

| Enrichment FDR | nGenes | Fold Enrichment | Pathways |
|---|---|---|---|
| 4.7E-20 | 88 | 3.3 | Mitotic cell cycle phase transition |
| 4.4E-34 | 145 | 3.2 | Mitotic cell cycle proc. |
| 9.0E-35 | 152 | 3.2 | Chromosome organization |
| 9.4E-36 | 160 | 3.1 | Mitotic cell cycle |
| 9.9E-26 | 120 | 3.1 | Reg. of cell cycle proc. |
| 3.9E-26 | 131 | 2.9 | MRNA metabolic proc. |
| 8.4E-35 | 183 | 2.8 | Cell cycle proc. |
| 4.6E-27 | 147 | 2.8 | Reg. of cell cycle |
| 2.5E-38 | 226 | 2.6 | Cell cycle |
| 1.2E-20 | 184 | 2.1 | Pos. reg. of macromolecule biosynthetic proc. |

**e**

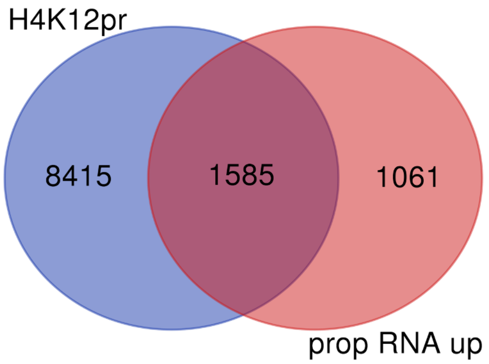


*P value = 1.69e-147*

**f**

| Enrichment FDR | nGenes | Fold Enrichment | Pathways |
|---|---|---|---|
| 5.2E-15 | 118 | 2.3 | Actin cytoskeleton organization |
| 7.8E-15 | 120 | 2.3 | Cell morphogenesis involved in differentiation |
| 5.4E-15 | 127 | 2.2 | Localization within membrane |
| 7.8E-15 | 127 | 2.2 | Actin filament-based proc. |
| 6.4E-18 | 161 | 2.2 | Cell morphogenesis |
| 5.4E-15 | 196 | 1.9 | Cytoskeleton organization |
| 9.7E-15 | 201 | 1.8 | Generation of neurons |
| 2.0E-15 | 217 | 1.8 | Neurogenesis |
| 3.0E-14 | 231 | 1.7 | Locomotion |
| 3.3E-14 | 230 | 1.7 | Cellular macromolecule localization |

**g**

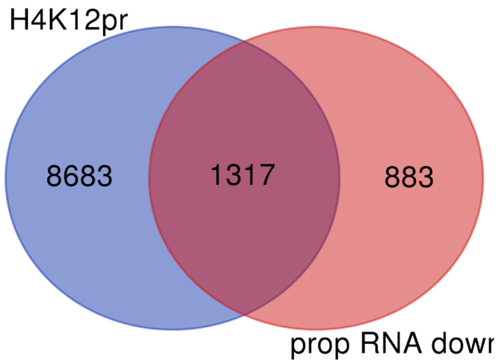


*P value = 1.35e-167*

**h**

| Enrichment FDR | nGenes | Fold Enrichment | Pathways |
|---|---|---|---|
| 2.4E-36 | 107 | 4.3 | Reg. of mRNA metabolic proc. |
| 4.2E-37 | 118 | 4 | RNA splicing |
| 1.0E-51 | 199 | 3.4 | MRNA metabolic proc. |
| 1.5E-42 | 191 | 3.1 | Chromosome organization |
| 4.2E-37 | 174 | 3 | Mitotic cell cycle proc. |
| 1.2E-40 | 195 | 2.9 | Mitotic cell cycle |
| 7.4E-38 | 186 | 2.9 | RNA processing |
| 7.1E-37 | 192 | 2.8 | Reg. of cell cycle |
| 2.2E-44 | 234 | 2.7 | Cell cycle proc. |
| 1.6E-47 | 286 | 2.5 | Cell cycle |

**Supplementary Fig. 7** | H3K18/H4K12pr ChIP-seq and propionyl RNA-seq integration. Overlap between TSS-proximal regions (+/- 1 Kb) for H3K18pr and H4K12pr associated genes by ChIP-seq, and upregulation of gene expression in 10 mM NaPr treated group **a**, **e** vs downregulation in the untreated group **c**, **g** by RNA-seq. Significance of overlap determined by hypergeometric test-generated *P* value. GO 'Biological Process' pathway terms associated with overlapping Kpr targets and upregulated genes **b**, **f** and downregulated genes **d**, **h** sorted by log2 fold enrichment.

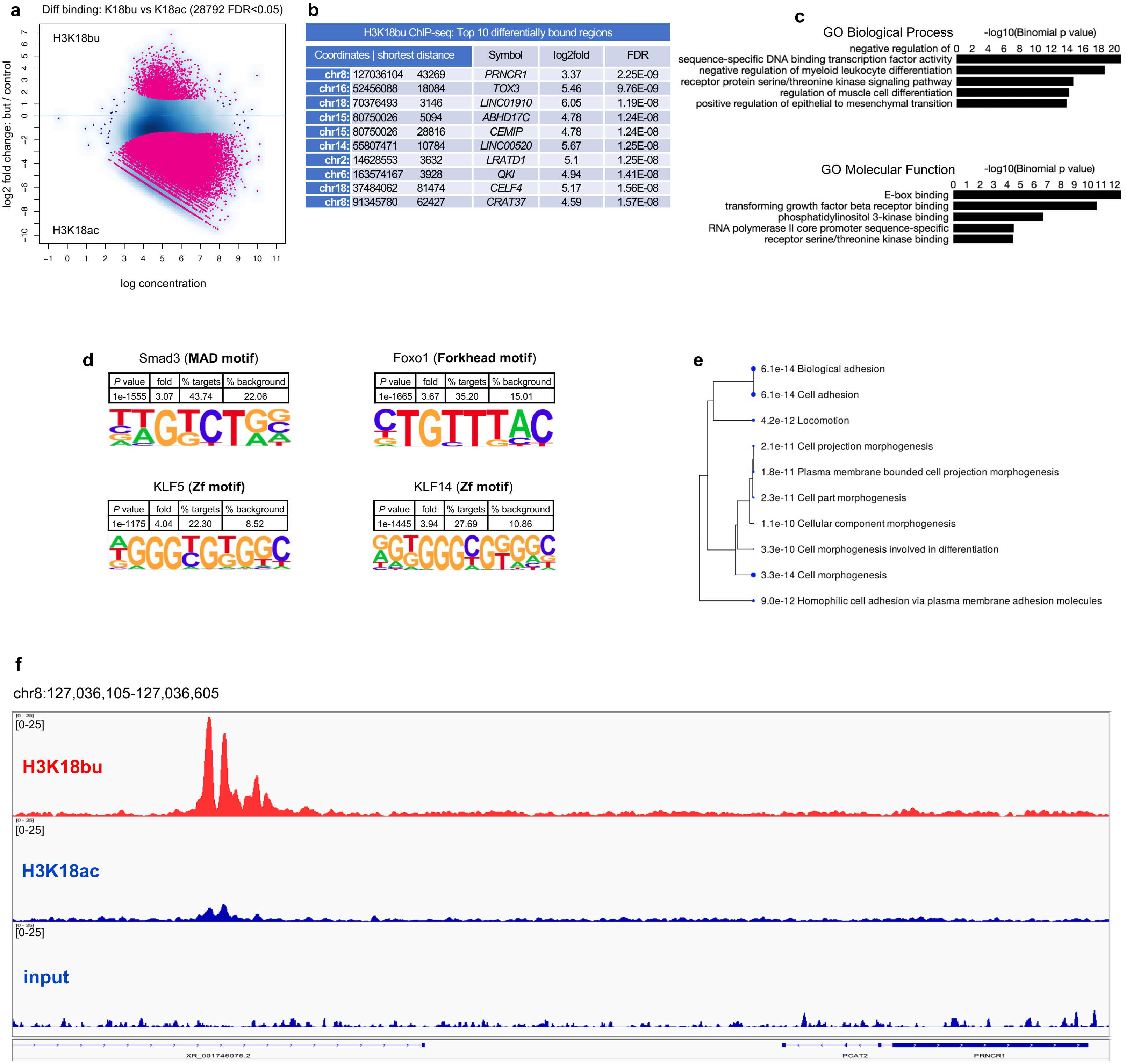


**Supplementary Fig. 8 |** Genome-wide H3K18bu distribution. **a** H3K18bu vs H3K18ac differential binding following 1 mM butyrate supplementation. Sites identified as significantly differentially bound are shown in red. Differential binding was performed by DiffBind package with DESeq2 using a two-sided tests for both increased and decreased binding affinity between conditions followed by by multiple hypothesis testing and FDR correction. **b** Top ten differentially bound regions associated with H3K18bu sorted by false-discovery rate adjusted *P* value (FDR < 0.05). **c** Top GO 'Biological Process' and 'Molecular Function' terms associated with H3K18bu-bound *cis*-regulatory elements determined by GREAT against a whole genome background using a binomial test over genomic regions, followed by multiple hypothesis testing using FDR corrected *P* values (FDR < 0.05). **d** Differential motif analysis of H3K18bu vs H3K18ac peaks was analyzed by HOMER, using a one-sided hypergeometric test for overrepresentation (enrichment) of motifs in the target sequences compared to the background, followed by multiple hypothesis testing and FDR correction. **e** Hierarchical clustering tree summary of correlations among significant pathways in H3K18pr-associated annotated genes. Hierarchical clustering of the pathways was performed using ShinyGO. Pathways were clustered together based on shared genes and gene enrichment analysis was performed using two-sided Fisher's exact test, and FDR correction was applied to adjust for multiple comparisons in the pathway analysis and hierarchical clustering. Size of dots indicates statistically significant FDR adjusted (FDR < 0.05) *P* values. **f** Signal tracks of 71 Kb-spanning *PRNCR1* region showing H3K18bu vs H3K18ac binding with input as background.

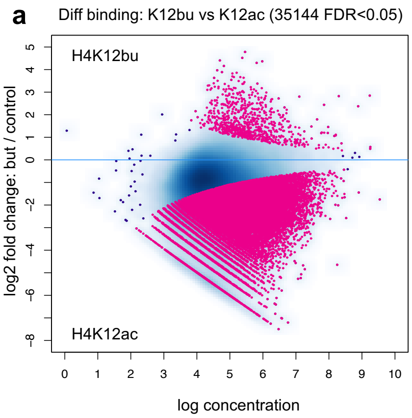


**b**

| H4K12bu ChIP-seq: Top 10 differentially bound regions | | | | |
|---|---|---|---|---|
| Coordinates \| shortest distance | | Symbol | log2fold | FDR |
| chr8: 127036130 | 43243 | *PRNCR1* | 2.85 | 3.14E-77 |
| chr7: 20217130 | 273 | *MACC1* | 2.8 | 6.53E-19 |
| chr1: 198621156 | 17014 | *PTPRC* | 2.98 | 4.36E-13 |
| chr15: 91345559 | 62648 | *CRAT37* | 4.52 | 6.85E-13 |
| chr6: 129491243 | 24825 | *LAMA2* | 2.69 | 1.15E-12 |
| chr8: 126842123 | 4431 | *PCAT1* | 2.83 | 1.57E-12 |
| chr3: 114706917 | 392417 | *ZBTB20* | 3.05 | 1.03E-11 |
| chr7: 138890353 | 58973 | *KIAA1549* | 2.71 | 1.70E-11 |
| chr2: 105583190 | 144678 | *FHL2* | 2.58 | 1.77E-11 |
| chr10: 62933853 | 13954 | *EGR2* | 2.68 | 4.19E-11 |

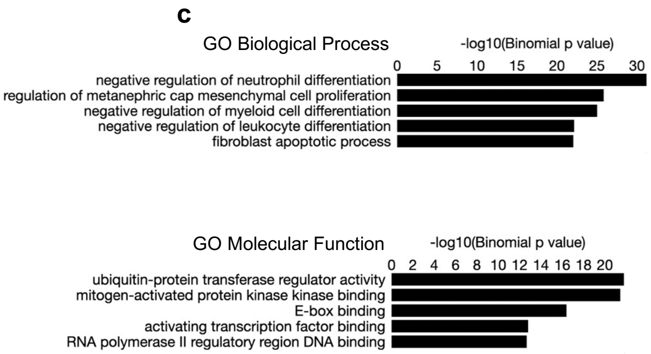


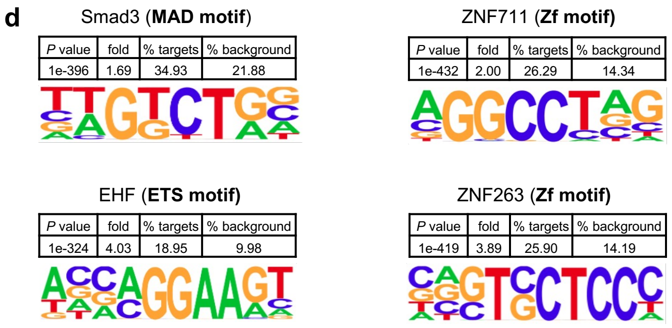


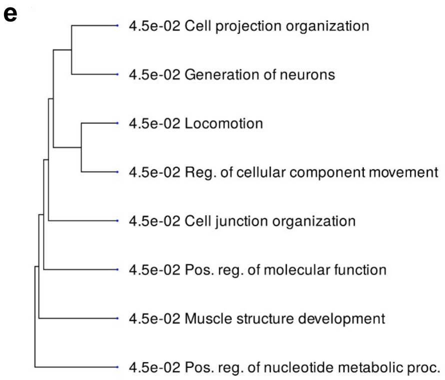


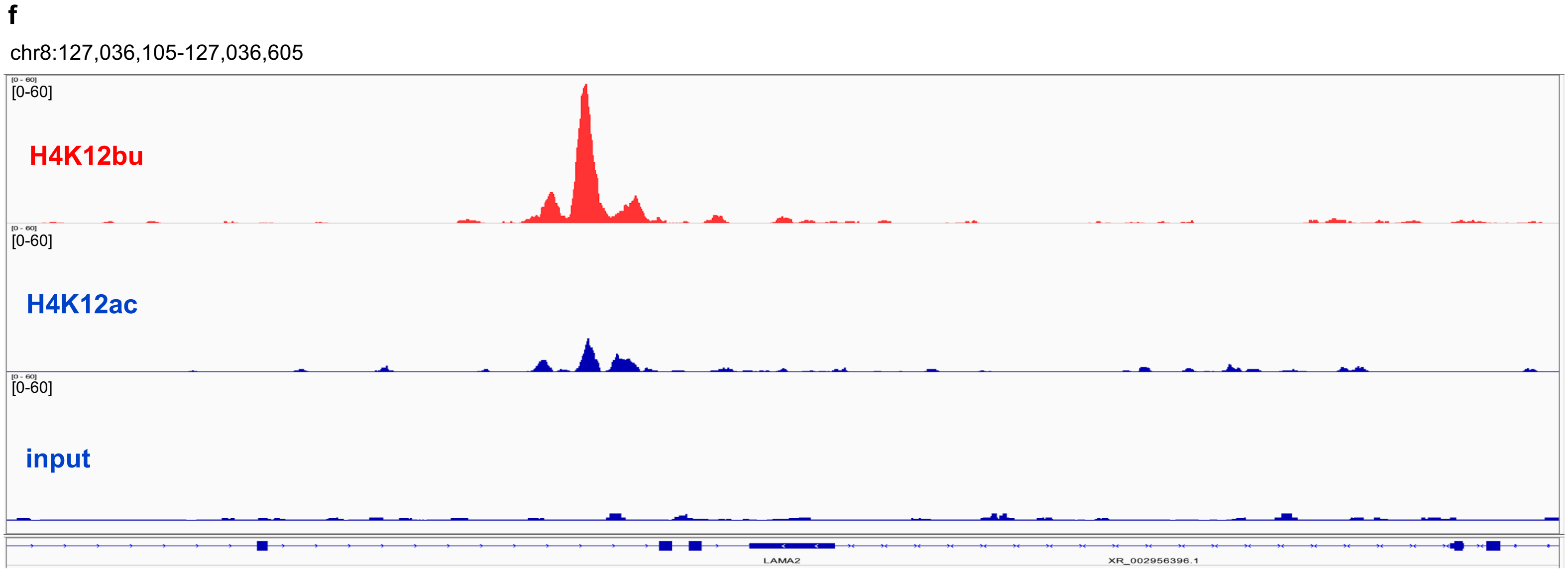


**Supplementary Fig. 9** | Genome-wide H4K12bu distribution. **a** H4K12bu vs H4K12ac differential binding following 1 mM butyrate supplementation. Sites identified as significantly differentially bound are shown in red. Differential binding was performed by DiffBind package with DESeq2 using a two-sided tests for both increased and decreased binding affinity between conditions followed by by multiple hypothesis testing and FDR correction. **b** Top ten differentially bound regions associated with H4K12bu sorted by false-discovery rate adjusted *P* value (FDR < 0.05). **c** Top GO 'Biological Process' and 'Molecular Function' terms associated with H3K18bu-bound *cis*-regulatory elements determined by GREAT against a whole genome background using a binomial test over genomic regions, followed by multiple hypothesis testing using FDR corrected *P* values (FDR < 0.05). **d** Differential motif analysis of H4K12bu vs H4K12ac peaks was analyzed by HOMER, using a one-sided hypergeometric test for overrepresentation (enrichment) of motifs in the target sequences compared to the background, followed by multiple hypothesis testing and FDR correction. **e** Hierarchical clustering tree summary of correlations among significant pathways in H4K12bu-associated annotated genes. Hierarchical clustering of the pathways was performed using ShinyGO. Pathways were clustered together based on shared genes and gene enrichment analysis was performed using two-sided Fisher's exact test, and FDR correction was applied to adjust for multiple comparisons in the pathway analysis and hierarchical clustering. Size of dots indicates statistically significant FDR adjusted (FDR < 0.05) *P* values. **f** Signal tracks of 20 Kb-spanning *LAMA* region showing H4K12bu vs H4K12ac binding with input as background.

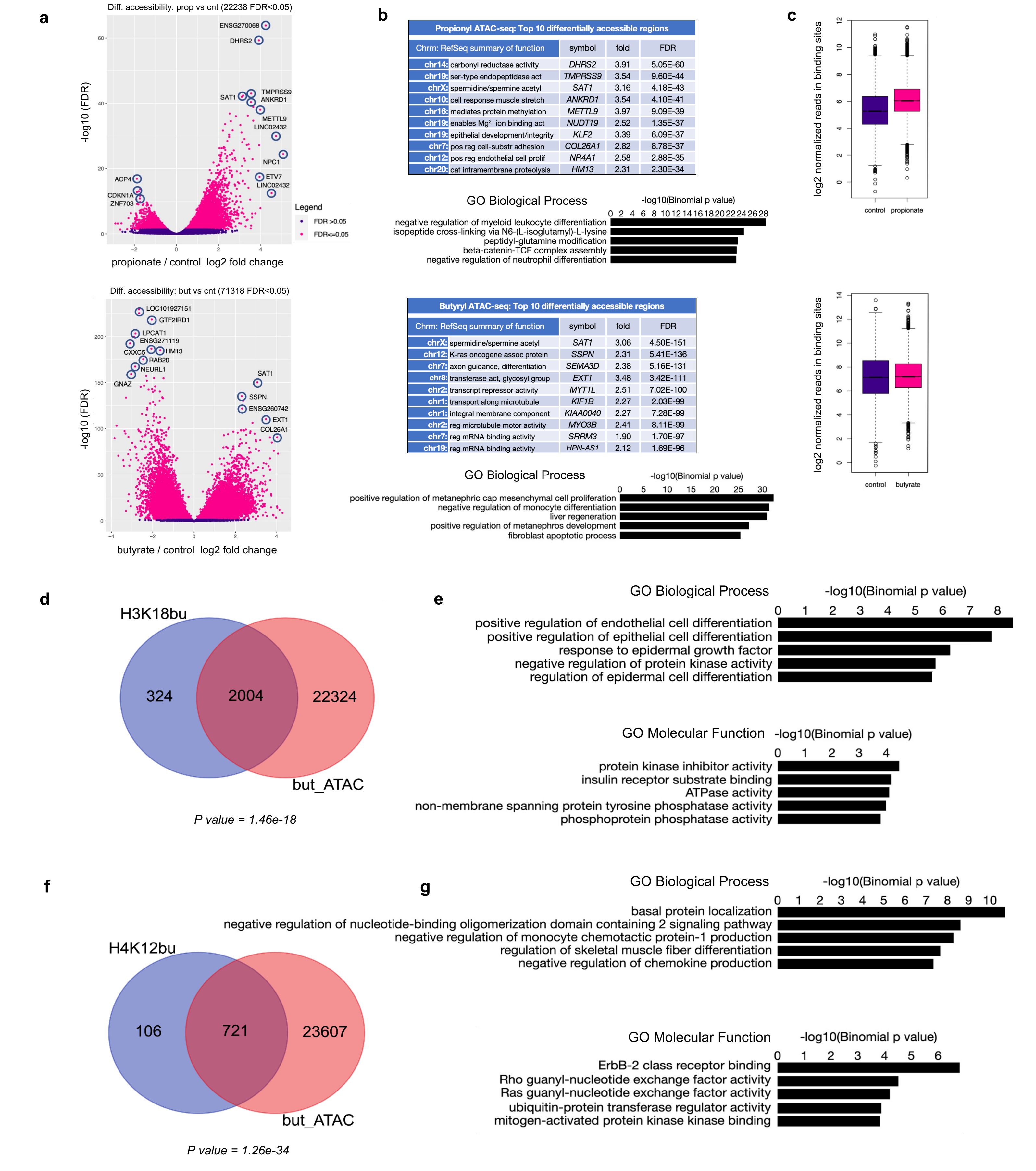


**Propionyl ATAC-seq: Top 10 differentially accessible regions**

| Chrm: RefSeq summary of function | symbol | fold | FDR |
|---|---|---|---|
| chr14: carbonyl reductase activity | *DHRS2* | 3.91 | 5.05E-60 |
| chr19: ser-type endopeptidase act | *TMPRSS9* | 3.54 | 9.60E-44 |
| chrX: spermidine/spermine acetyl | *SAT1* | 3.16 | 4.18E-43 |
| chr10: cell response muscle stretch | *ANKRD1* | 3.54 | 4.10E-41 |
| chr16: mediates protein methylation | *METTL9* | 3.97 | 9.09E-39 |
| chr19: enables $Mg^{2+}$ ion binding act | *NUDT19* | 2.52 | 1.35E-37 |
| chr19: epithelial development/integrity | *KLF2* | 3.39 | 6.09E-37 |
| chr7: pos reg cell-substr adhesion | *COL26A1* | 2.82 | 8.78E-37 |
| chr12: pos reg endothelial cell prolif | *NR4A1* | 2.58 | 2.88E-35 |
| chr20: cat intramembrane proteolysis | *HM13* | 2.31 | 2.30E-34 |

**Butyryl ATAC-seq: Top 10 differentially accessible regions**

| Chrm: RefSeq summary of function | symbol | fold | FDR |
|---|---|---|---|
| chrX: spermidine/spermine acetyl | *SAT1* | 3.06 | 4.50E-151 |
| chr12: K-ras oncogene assoc protein | *SSPN* | 2.31 | 5.41E-136 |
| chr7: axon guidance, differentiation | *SEMA3D* | 2.38 | 5.16E-131 |
| chr8: transferase act, glycosyl group | *EXT1* | 3.48 | 3.42E-111 |
| chr2: transcript repressor activity | *MYT1L* | 2.51 | 7.02E-100 |
| chr1: transport along microtubule | *KIF1B* | 2.27 | 2.03E-99 |
| chr1: integral membrane component | *KIAA0040* | 2.27 | 7.28E-99 |
| chr2: reg microtubule motor activity | *MYO3B* | 2.41 | 8.11E-99 |
| chr7: reg mRNA binding activity | *SRRM3* | 1.90 | 1.70E-97 |
| chr19: reg mRNA binding activity | *HPN-AS1* | 2.12 | 1.69E-96 |

**Supplementary Fig. 10** | Propionyl and butyryl ATAC-seq and Kbu ChIP-seq and ATAC-seq integration. **a** Differential accessibility following propionate and butyrate supplementation. Sites identified as significantly differentially accessible are shown in red. n = 3 technical replicates for each condition. Differential accessibility was performed by DiffBind package with DESeq2 using a two-sided tests for both increased and decreased binding affinity between conditions followed by by multiple hypothesis testing and FDR correction. **b** Top ten differentially bound regions associated with propionate and butyrate treatment sorted by false-discovery rate adjusted *P* value (FDR < 0.05) and top GO 'Biological Process' terms determined by GREAT against a whole genome background using a binomial test over genomic regions, followed by multiple hypothesis testing using FDR corrected *P* values (FDR < 0.05). **c** Normalized reads in accessible sites following propionate and butyrate treatment. Box plots display: The minimum, first quartile (Q1, 25th percentile), median, third quartile (Q3, 75th percentile), and maximum. The bottom of the box is Q1 and the top of the box is Q3. The line within the box represents the median (50th percentile) value. The whiskers extend to the most extreme data points within 1.5 times the IQR (interquartile range). **d, f** Overlap between Kbu bound genes and differentially accessible regions following 1 mM butyrate treatment. Significance of overlap determined by hypergeometric test-generated *P* value. **e, g** GO 'Biological Process' and 'Molecular Function' pathway terms associated with Kbu bound genomic coordinates that are also present in ATAC-seq data set sorted by binomial *P* value.
