## Supplementary material for "Short-chain fatty acid metabolites propionate and butyrate are unique epigenetic regulatory elements linking diet, metabolism and gene expression": Nshanian_et_al_Extended_Figures

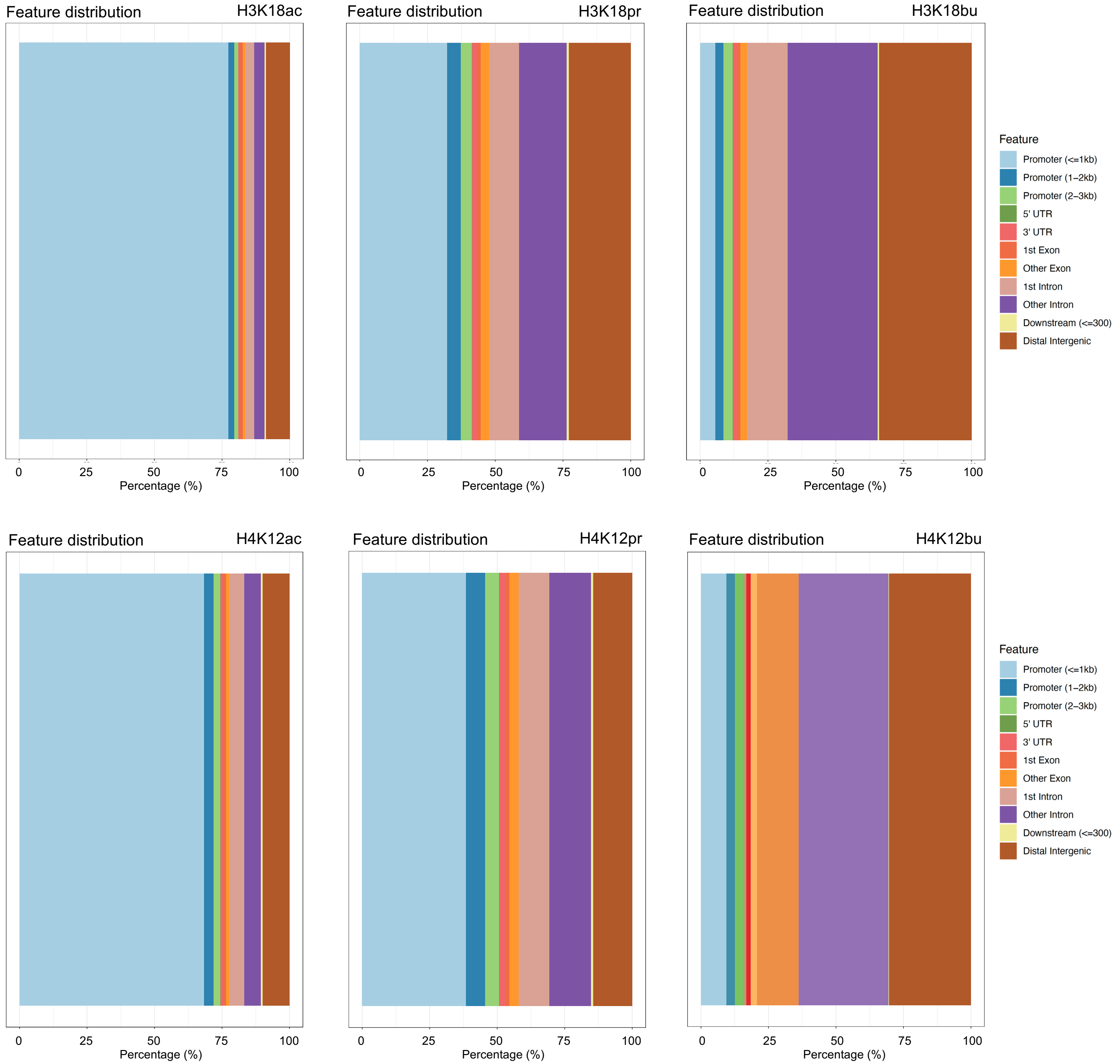


**Extended Fig. 1** | Feature distribution of H3K18ac/pr/bu (top panel) and H4K12ac/pr/bu (bottom panel) associated regions (+/- 3 Kb of TSS). H3K18ac/H4K12ac annotations were taken from ChIP-seq data that was generated without any treatment. H3K18pr/H4K12pr and H3K18bu/H4K12bu ChIP-seq experiments were performed following 10 mM NaPr, and 1 mM NaBu treatments, respectively. The x-axis provides the percentage of sites while colored regions represent distance from TSS. Results obtained using ChIPSeeker R package.

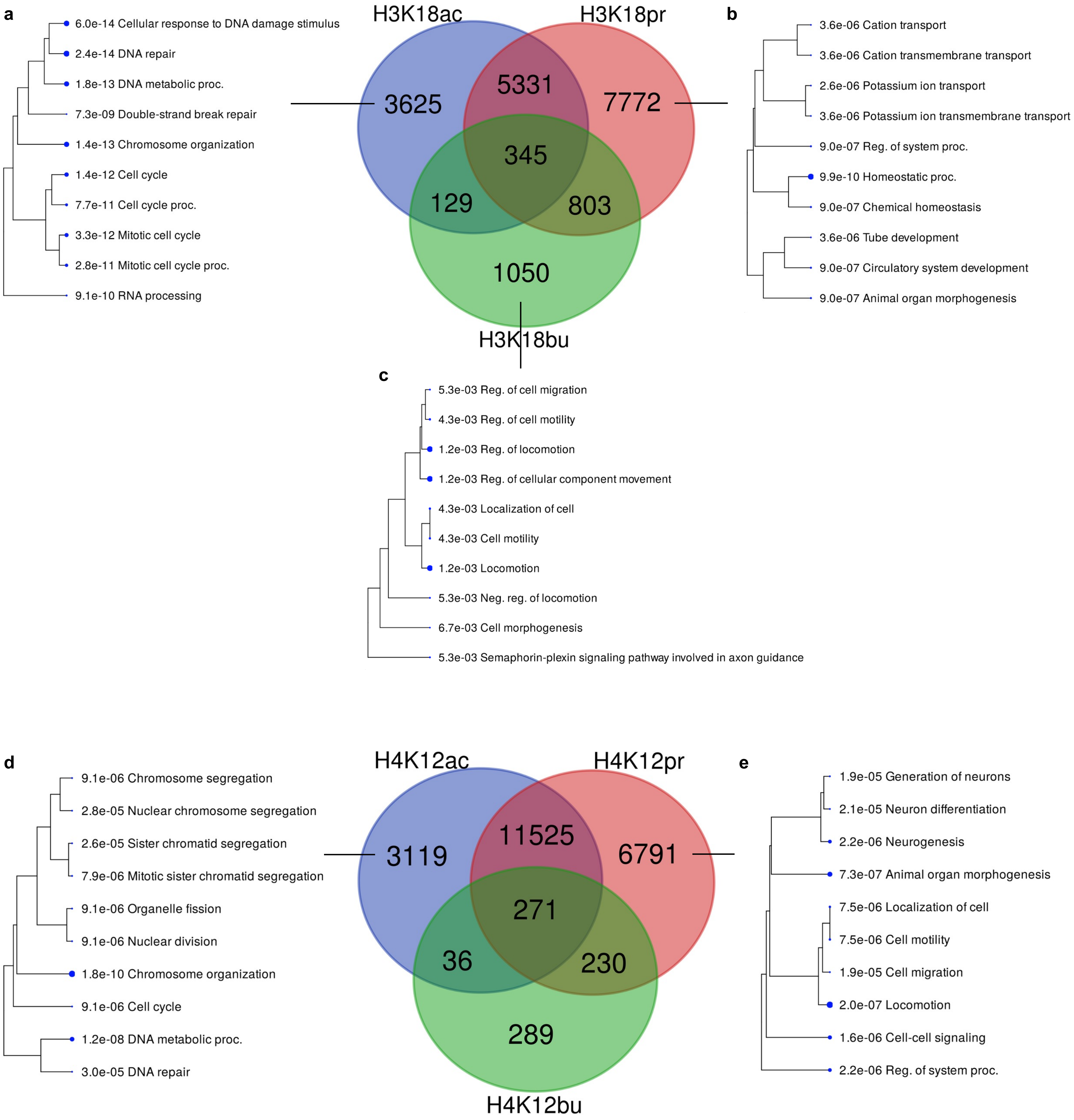


**Extended Fig. 2 |** Kac/pr/bu annotated features overlap. **a, d** Hierarchical clustering of GO 'Biological Process' terms associated with Kac but not Kpr or Kbu. **b, e** Hierarchical clustering of GO 'Biological Process' terms associated with Kpr but not Kac or Kbu. **c** Hierarchical clustering of GO 'Biological Process' terms associated with Kbu but not Kac or Kpr. Hierarchical clustering of the pathways was performed using ShinyGO. Pathways were clustered together based on shared genes and gene enrichment analysis was performed using two-sided Fisher's exact test, and FDR correction was applied to adjust for multiple comparisons in the pathway analysis and hierarchical clustering. Size of dots indicates statistically significant FDR adjusted (FDR < 0.05) *P* values.

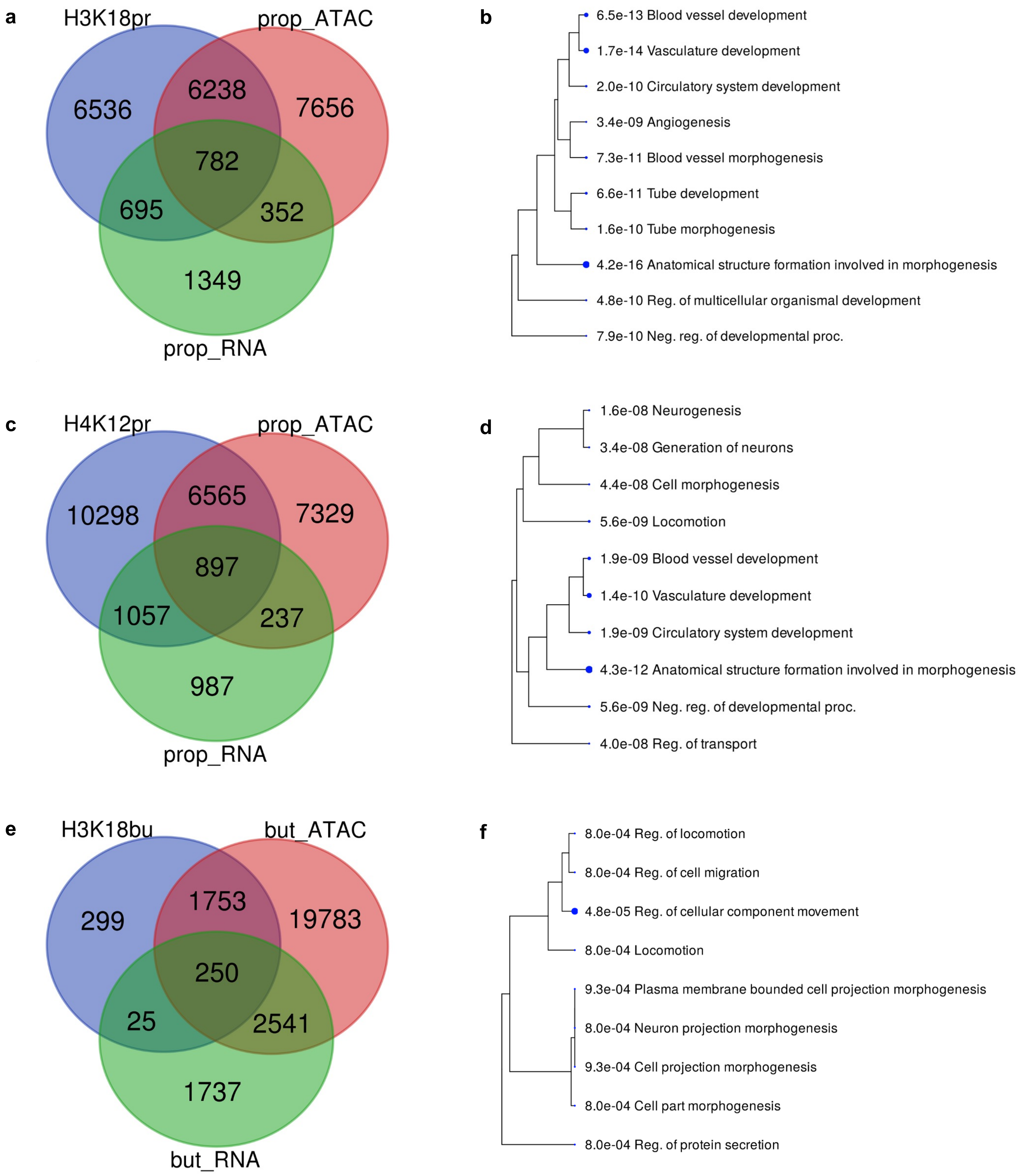


**Extended Fig. 3 |** Kpr/bu and ATAC/RNA-seq shared annotated features. **a** H3K18pr, propionyl ATAC/RNA-seq shared annotated features. **b** Hierarchical cluttering of GO ‘Biological Process’ terms associated 782 shared features. **c** H4K12pr, propionyl ATAC/RNA-seq shared features. **d** Hierarchical clustering of GO ‘Biological Process’ terms associated with 897 shared features. Hierarchical clustering of the pathways was performed using ShinyGO. Pathways were clustered together based on shared genes and gene enrichment analysis was performed using two-sided Fisher's exact test, and FDR correction was applied to adjust for multiple comparisons in the pathway analysis and hierarchical clustering. Size of dots indicates statistically significant FDR adjusted (FDR < 0.05) *P* values. **e** H3K18bu butyryl ATAC/RNA-seq shared features. **f** Hierarchical cluttering of GO ‘Biological Process’ terms associated with 250 shared features.

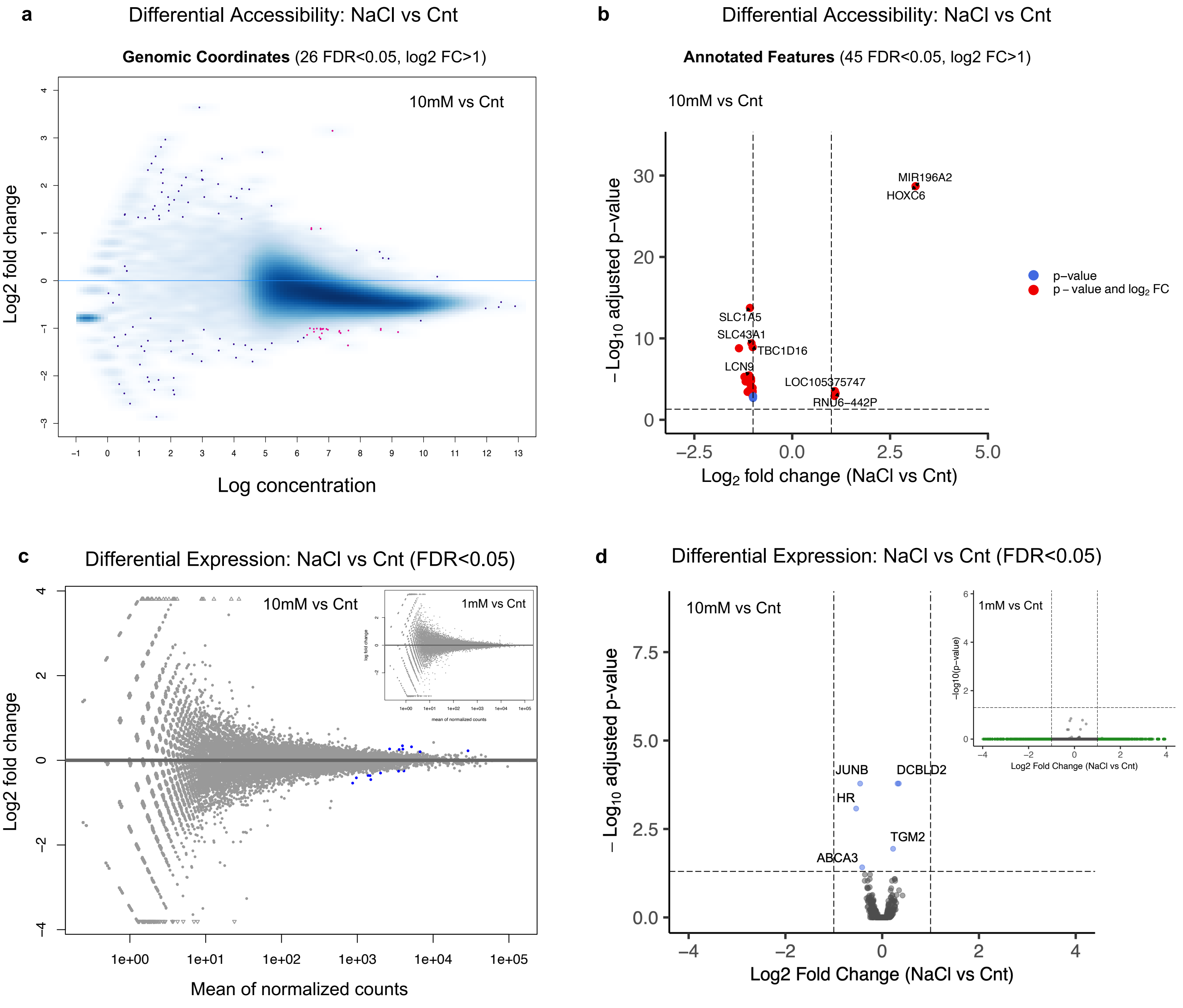


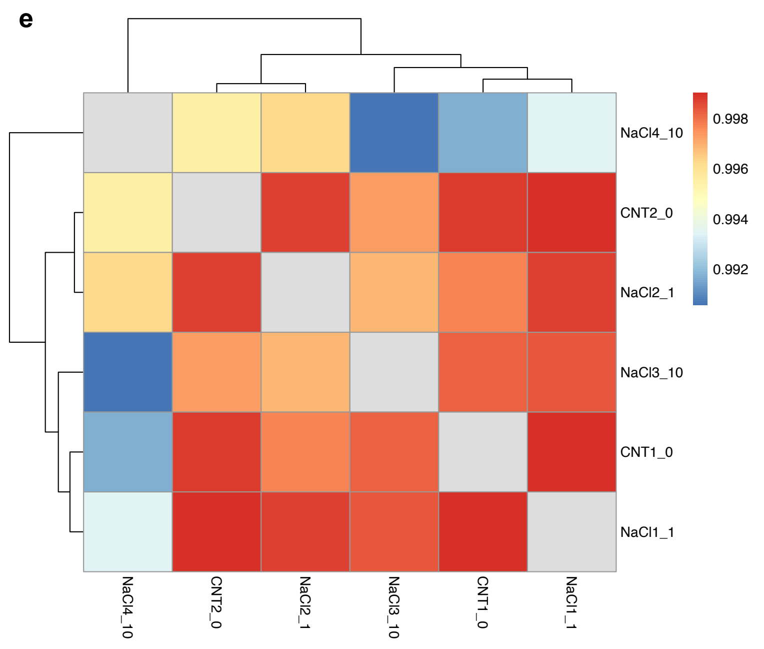


**Extended Fig. 4 |** Effect of $Na^+$ on differential accessibility and expression. **a** ATAC-seq MA plot of differential accessibility in 10 mM NaCl-treated vs untreated cells (SW480). Differential analysis was performed by DESeq2 using two-sided Wald tests to identify differentially expressed or accessible genes. The results were then adjusted for multiple comparisons using the FDR correction method. Sites identified as significantly differentially accessible (FDR<0.05, log2 FC>1) are shown in red. n = 3 experimental replicates for each condition. **b** Volcano plot of differential accessibility in 10 mM NaCl-treated vs untreated cells. **c** RNA-seq MA plot of differential expression in 10 mM NaCl treated vs untreated cells. Sites identified as significantly differentially expressed (FDR<0.05) are shown blue (*Insert*: 1 mM vs Cnt). **d** Volcano plot of differential expression in 10 mM NaCl-treated vs untreated cells. Sites identified as significantly differentially expressed by FDR only (FDR<0.05) are shown blue (*Insert*: 1 mM vs Cnt). **e** Correlation heatmap of RNA-seq data showing clustering replicates from 1 and 10 mM NaCl-treated vs untreated groups. Normalized measurement of the covariance between replicates is expressed by Pearson's correlation coefficient.

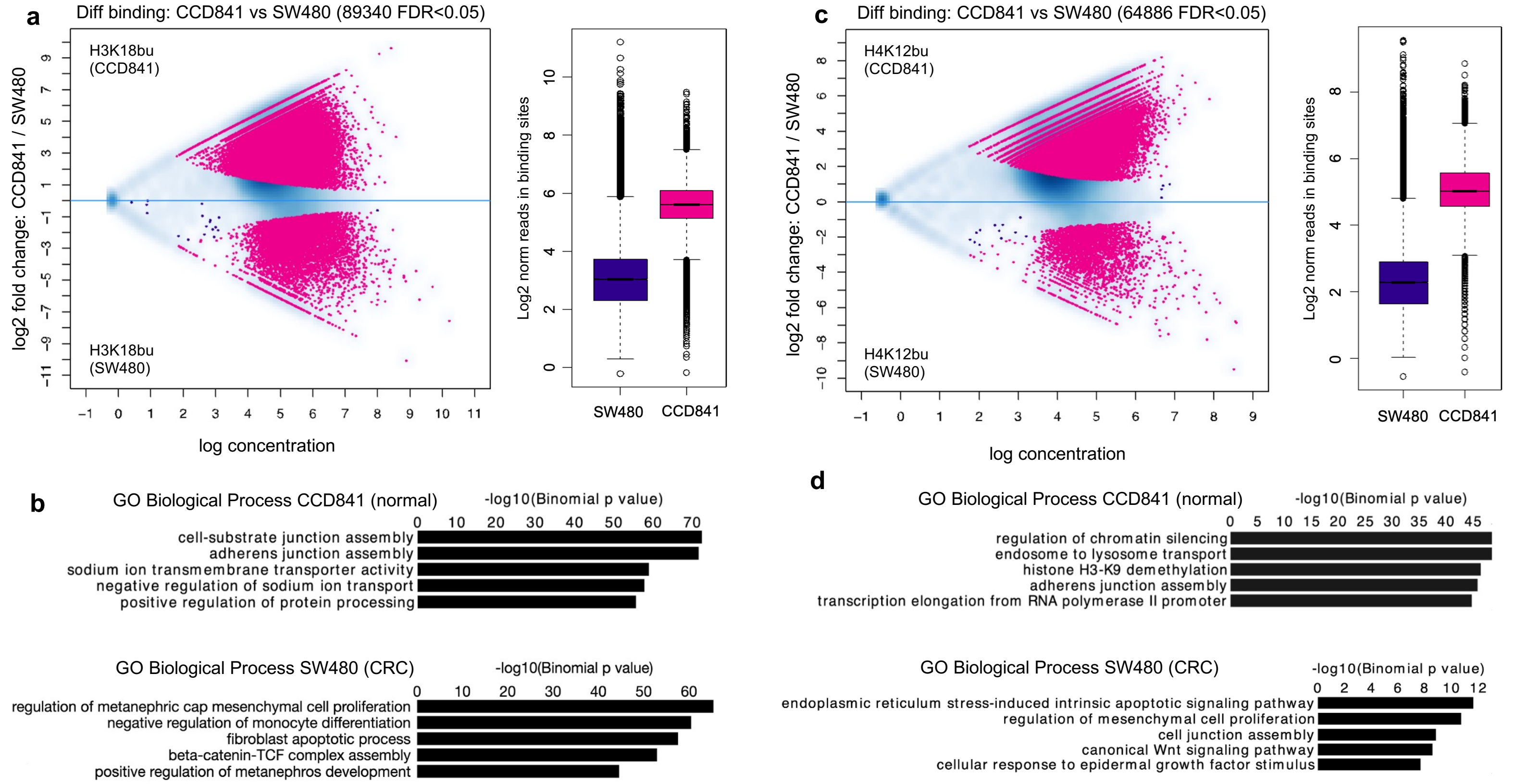


**e**

H3K18bu-assoc. CRC genes (CCD841)

| symbol | fold | FDR |
|---|---|---|
| *MYC* | -4.42 | 1.66E-29 |
| *FOSL1* | -3.71 | 2.99E-12 |
| *JUNB* | -3.08 | 1.15E-07 |
| *SMAD2* | -2.50 | 2.83E-04 |
| *CCDN1* | -6.85 | 9.32E-09 |
| *PDGFA* | -3.38 | 1.18E-08 |
| *TGFBI* | -3.22 | 8.64E-12 |
| *TCF4* | -3.79 | 2.82E-09 |
| *PRNCR1* | -7.56 | 6.69E-32 |
| *ZNF280D* | -3.49 | 6.82E-16 |

**f**

H4K12bu-assoc. CRC genes (CCD841)

| symbol | fold | FDR |
|---|---|---|
| *FOXK1* | -4.69 | 1.47E-09 |
| *TP63* | -4.16 | 2.00E-07 |
| *FOXQ1* | -2.35 | 1.22E-04 |
| *BCL6* | -3.11 | 1.22E-04 |
| *PDGFA* | -2.68 | 2.31E-04 |
| *FOSL1* | -3.85 | 5.50E-06 |
| *MYC* | -5.03 | 8.37E-13 |
| *TGFA* | -3.46 | 3.25E-05 |
| *CCND1* | -4.84 | 8.25E-07 |
| *PRNCR1* | -6.77 | 2.53E-27 |

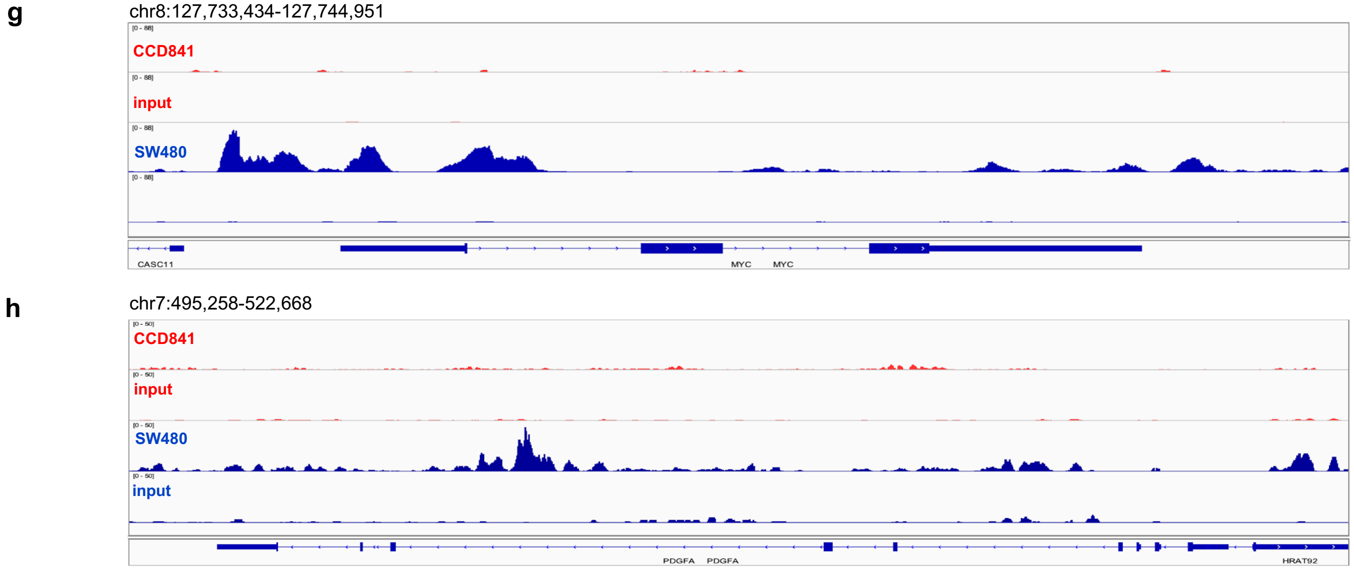


**Extended Fig. 5 | a** H3K18bu differential binding in cancer (SW480) vs normal (CCD841) cells and normalized reads in H3K18bu-associated binding sites following 1 mM NaBu treatment. Sites identified as significantly differentially bound are shown in red. n = 3 technical replicates for each condition. Differential binding was performed by DiffBind package with DESeq2 using a two-sided tests for both increased and decreased binding affinity between conditions followed by multiple hypothesis testing and FDR correction (FDR < 0.05). Box plots display: The minimum, first quartile (Q1, 25th percentile), median, third quartile (Q3, 75th percentile), and maximum. The bottom of the box is Q1 and the top of the box is Q3. The line within the box represents the median (50th percentile) value. The whiskers extend to the most extreme data points within 1.5 times the IQR (interquartile range). **b** Top GO Biological Process terms for H3K18bu-associated *cis*-regulatory elements in cancer vs normal cells determined by GREAT against a whole genome background using a binomial test over genomic regions, followed by multiple hypothesis testing using FDR corrected *P* values (FDR < 0.05). **c** H4K12bu differential binding in cancer vs normal cells and normalized reads in H4K12bu associated binding sites following 1 mM NaBu treatment. **d** Top GO Biological Process terms for H4K12bu-associated *cis*-regulatory elements in cancer vs normal cells. **e, f** CRC differentially bound genes associated with H3K18bu/H4K12bu. **g, h** Signal tracks showing differential binding in *MYC* and *PDGFA* regions in cancer vs normal cells.

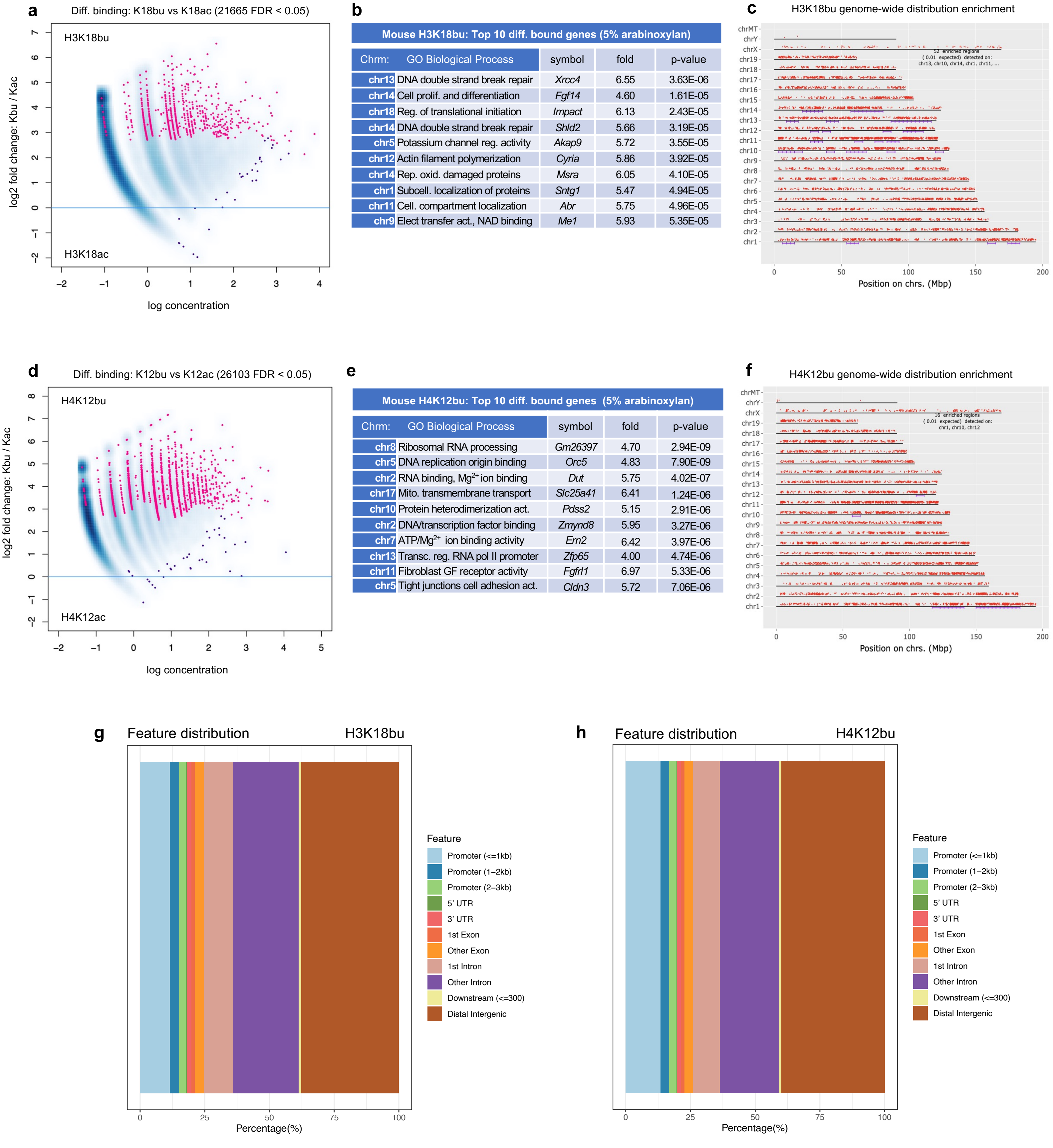


**b** Mouse H3K18bu: Top 10 diff. bound genes (5% arabinoxylan)

| Chrm: | GO Biological Process | symbol | fold | p-value |
|---|---|---|---|---|
| chr13 | DNA double strand break repair | *Xrcc4* | 6.55 | 3.63E-06 |
| chr14 | Cell prolif. and differentiation | *Fgf14* | 4.60 | 1.61E-05 |
| chr18 | Reg. of translational initiation | *Impact* | 6.13 | 2.43E-05 |
| chr14 | DNA double strand break repair | *Shld2* | 5.66 | 3.19E-05 |
| chr5 | Potassium channel reg. activity | *Akap9* | 5.72 | 3.55E-05 |
| chr12 | Actin filament polymerization | *Cyria* | 5.86 | 3.92E-05 |
| chr14 | Rep. oxid. damaged proteins | *Msra* | 6.05 | 4.10E-05 |
| chr1 | Subcell. localization of proteins | *Sntg1* | 5.47 | 4.94E-05 |
| chr11 | Cell. compartment localization | *Abr* | 5.75 | 4.96E-05 |
| chr9 | Elect transfer act., NAD binding | *Me1* | 5.93 | 5.35E-05 |

**e** Mouse H4K12bu: Top 10 diff. bound genes (5% arabinoxylan)

| Chrm: | GO Biological Process | symbol | fold | p-value |
|---|---|---|---|---|
| chr8 | Ribosomal RNA processing | *Gm26397* | 4.70 | 2.94E-09 |
| chr5 | DNA replication origin binding | *Orc5* | 4.83 | 7.90E-09 |
| chr2 | RNA binding, $Mg^{2+}$ ion binding | *Dut* | 5.75 | 4.02E-07 |
| chr17 | Mito. transmembrane transport | *Slc25a41* | 6.41 | 1.24E-06 |
| chr10 | Protein heterodimerization act. | *Pdss2* | 5.15 | 2.91E-06 |
| chr2 | DNA/transcription factor binding | *Zmynd8* | 5.95 | 3.27E-06 |
| chr7 | ATP/$Mg^{2+}$ ion binding activity | *Ern2* | 6.42 | 3.97E-06 |
| chr13 | Transc. reg. RNA pol II promoter | *Zfp65* | 4.00 | 4.74E-06 |
| chr11 | Fibroblast GF receptor activity | *Fgfrl1* | 6.97 | 5.33E-06 |
| chr5 | Tight junctions cell adhesion act. | *Cldn3* | 5.72 | 7.06E-06 |

**Extended Fig. 6 |** H3K18bu vs ac and H4K12bu vs ac differential binding from mouse intestine on HSF + 5 % arabinoxylan diet. **a, b, d, e** Sites identified as significantly differentially bound are shown in red. n = 3 biological replicates for each condition. Differential binding was performed by DiffBind package with DESeq2 using a two-sided tests for both increased and decreased binding affinity between conditions followed by multiple hypothesis testing and FDR correction. **c, f** Chromosomal positions of H3K18bu and H4K12bu-associated regions represented by red dots. Purple lines represent statistically significant enrichment compared to input. The genome was scanned with a sliding window (size 6 Mb) further subdivided into 2 equal-sized steps for sliding. Within each window a hypergeometric test was used to test for enrichment over all protein-coding genes in the genome. FDR adjusted *P* value cutoff for window was 1E-05. Gene chromosomal mapping performed using ShinyGO. **g, h** Feature distribution of H3K18bu and H4K12bu bound regions (+/- 3 Kb of TSS). The x-axis provides the percentage of sites while colored regions represent distance from TSS. Results obtained using ChIPSeeker R package.

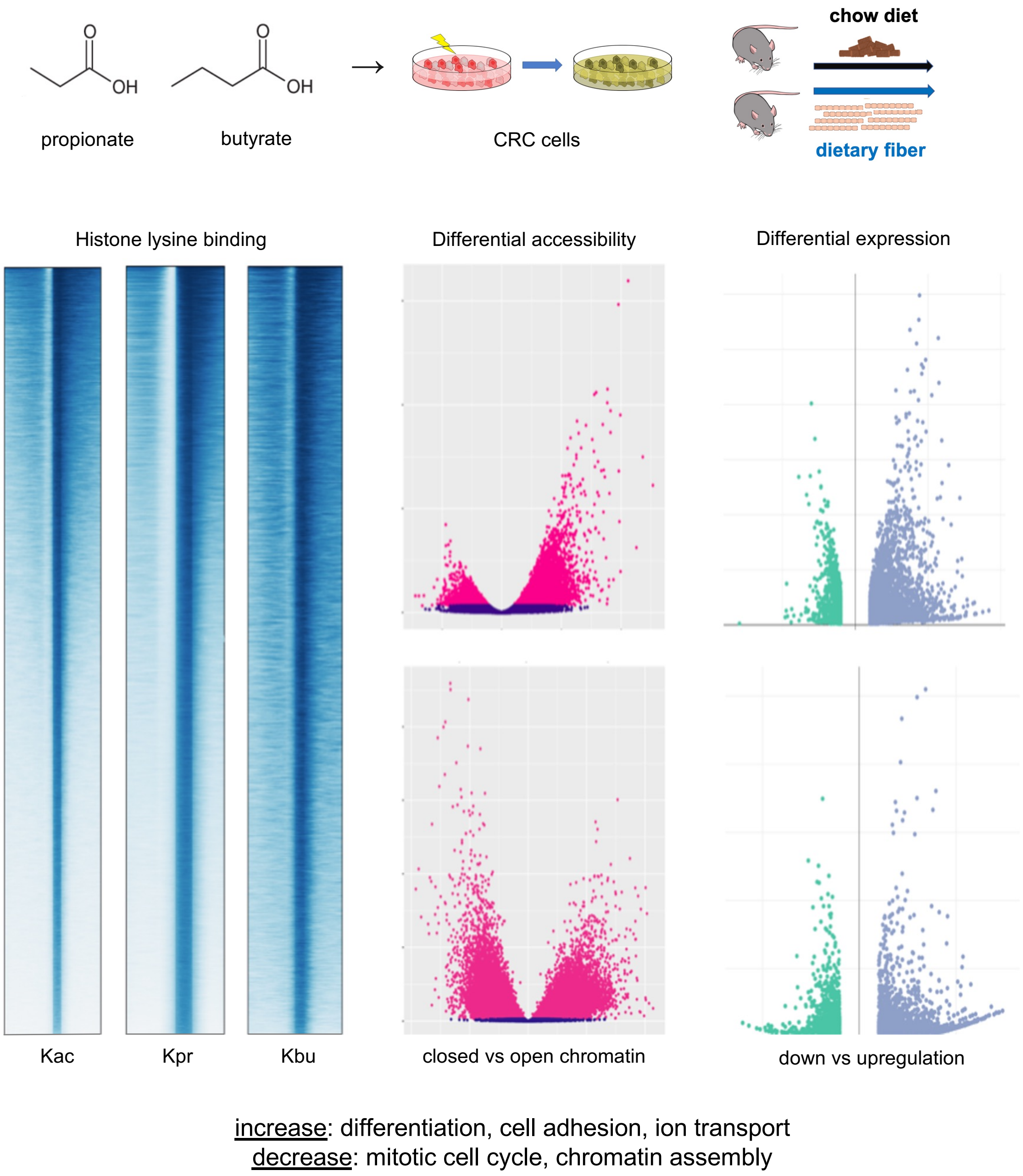


**Extended Fig. 7 |** Overview of SCFAs propionate and butyrate as regulatory elements affecting histone binding, chromatin accessibility and gene expression. Upper panels: *In vitro* cellular perturbations and *in vivo* fiber supplementation experimental designs. Bottom panels: Changes in TSS distribution profiles, differential chromatin accessibility and expression following SCFA supplementation. Top and Bottom volcano plots showing differential accessibility and expression following propionate and butyrate supplementation, respectively.
